# PhaseWY: A pipeline for haplotype phasing, sex chromosome identification and extraction of sex-limited sequences

**DOI:** 10.64898/2026.06.17.732863

**Authors:** Simon J Ellerstrand, Allison Churcher, Verena E. Kutschera, Bengt Hansson

**Affiliations:** Department of Biology, Lund University, Ecology Building, 223 62 Lund, Sweden; Department of Plant Physiology, National Bioinformatics Infrastructure Sweden, 901 87 Umeå, Sweden; Department of Biochemistry and Biophysics, National Bioinformatics Infrastructure Sweden, Science for Life Laboratory, Stockholm University, Box 1031, SE-17121 Solna, Sweden

**Keywords:** Snakemake, reproducible research, sex chromosome evolution, evolutionary strata

## Abstract

Sex chromosomes are central to many ecological and evolutionary processes. Evidence has accumulated that sex chromosome systems vary extensively in age, turnover and transitions, motivating renewed efforts to study the diversity of sex chromosome systems across the tree of life. However, successful genomic detection of sex chromosomes depends on several factors, including the size and divergence time, background genetic diversity, and the number of sequenced females and males. In addition, technical challenges associated with sequencing and analysing the sex-limited Y/W chromosome remain. Here, we present PhaseWY, an automated Snakemake pipeline that uses whole-genome sequencing data from multiple female and male individuals to identify sex-chromosomal regions and extract the corresponding Y/W sequences. PhaseWY (i) detects sex differences in alignment depth, (ii) applies read-based and statistical haplotype phasing, (iii) identifies sex-linked regions using haplotype clustering, and (iv) subsets autosomal, X/Z- and Y/W-linked variants for downstream analyses. We applied PhaseWY to simulated data to benchmark factors influencing sex-linkage detection and successful extraction of Y/W-linked variants. To demonstrate its practical utility, we further applied PhaseWY to the neo-sex chromosome system in *Alauda* larks (Alaudidae) and performed a range of downstream analyses demonstrating the scope of applications of the PhaseWY output. We conclude that PhaseWY provides an easy-to-use and reproducible tool for population-genomic analyses in non-model organisms, with particular importance for advancing our understanding of sex-chromosome evolution.

## Background

Sex chromosomes often evolve from homologous pairs of autosomes through the acquisition of one or more sex-determining mutations. The sex-linked region can subsequently expand via the formation of non-recombining haplotypes that associate alleles across loci, a process that can occur gradually or instantly over larger chromosomal regions through structural mutations such as inversions (Ohno, 1967). These events generate sex-specific haplotypes for each sex chromosome and are hypothesised to be primarily driven by selection favouring linkage between sexually antagonistic loci and the sex-determining locus (Bull, 1983; Fisher, 1930; Rice, 1987). Regardless of whether such expansions occur, a short recombining region often remains; this region, referred to as the pseudoautosomal region (PAR), enables proper meiotic paring and segregation (Graves et al., 1998; Otto et al., 2011; Zhou et al., 2014).

As evolutionary time progresses, the non-recombining regions accumulate sequence differences and diverge. Eventually, the sex-limited chromosome (the Y in XY males or W in ZW females, i.e. the heterogametic sex) often degenerate as a consequence of the accumulation of deleterious mutations and inefficient selection (Bachtrog, 2008). Degeneration of the sex-limited chromosome can negatively affect fitness in the heterogametic sex, e.g. through unbalanced gene expression (the dosage hypothesis; Chandler, 2017; Ohno, 1967; Zhu et al., 2025) or through exposing recessive deleterious mutations segregating on the X/Z chromosome (the unguarded X hypothesis; Robert, 1985; Xirocostas et al., 2020). In addition, repetitive content and deleterious transposable elements may accumulate on the sex-limited sex chromosome, which may not only reduce heterogametic viability , but also act as a barrier to geneflow and hybridization by contributing to hybrid dysfunction in the heterogametic sex when introduced into a naïve genomic environment, consistent with Haldane’s rule (Haldane, 1922).

Over the past decades extensive sex chromosome characterisation initiatives of non-model organisms have accumulated evidence that sex chromosome systems across the tree of life can vary extensively in age, turnover, evolutionary transitions and heteromorphy, even within systems otherwise considered relatively stable, such as mammals and birds (Bachtrog et al., 2014; Cortez et al., 2014; Jeffries et al., 2018; Sigeman et al., 2019; Xu et al., 2019). This has spurred the development of several methods for detecting sex-linked genomic regions. While sex-specific karyotyping has traditionally been used to identify sex chromosomes through differences in chromosome number or heteromorphy, such methods are less precise for revealing young sex chromosomes, which often remain homomorphic in size (Valenzuela et al., 2003). Sex chromosomes can likewise be identified and assembled through various high-resolution sequencing techniques, such as long-read approaches. However, these may still encounter problems due to low haplotype divergence or biased accumulation of repetitive content on the sex-limited chromosome (Tomaszkiewicz et al., 2017). To overcome these limitations, a commonly used approach nowadays is to identify sex chromosomes using short-read sequencing of males and females to detect sex-specific genomic signatures, such as FindZX (Sigeman et al., 2022), SexFindR (Grayson et al., 2022), SEXDETector (Muyle et al., 2016), and more (see Palmer et al., 2019 for a detailed overview).

By alignment of short-reads of both sexes to a homogametic reference assembly, it is possible to detect sex differences in alignment depth (indicating older and degenerated Y/W regions that do not align to the homogametic reference; FindZX and SexFindR) or inflated heterozygosity/SNP density in the heterogametic sex (younger Y/W regions with low sequence divergence align; FindZX and SexFindR). With several females and males, it is possible to apply methods such as sex-specific differentiation with F_ST_ (younger regions which have accumulated sequence divergence; SexFindR), or sex-specific GWAS/kmer GWAS (which can detect very small and young regions with enough samples; SexFindR). Even with a fragmented assembly, liftover techniques can be applied to visualise the sex-linked region in the context of chromosome-level assembly of a closely related species (FindZX). However, the accuracy of these techniques in detecting sex-linked regions may depend on a number of factors, such as study design (the number of sequenced females and males), divergence time of sex-linked regions and effective population sizes (e.g. noise from background genetic diversity or increased divergence through drift). The impact of such factors on accurate sex-chromosome detection has not been well benchmarked, and several methods are often applied in conjunction in unknown sex determination systems to improve detection of sex-linked regions.

While identification of sex chromosomes can often be successful with relatively few resources, technical difficulties remain in acquiring and studying variation segregating on the sex chromosomes, since none of the methods mentioned above extract sequences from the sex-limited Y/W chromosomes. Methods employed for such tasks are segregation analysis (SEXDETector) or linkage mapping. Both require pedigree data, which may not be easily attained in non-model organisms. While more sophisticated sequencing techniques can offer solutions – such as long-read, Hi-C and 10x Chromium sequencing – they put a higher demand on sample quality and can be costly to apply to numerous individuals.

Due to the technical challenges associated with sex-limited chromosomes, sequencing projects often focus on the homogametic sex to avoid confounding signals resulting from spurious alignments of homologous Y/W sequences (Tomaszkiewicz et al., 2017). This is unfortunate, since sex chromosomes can play a disproportionately large role in evolutionary processes (Haldane, 1922; Meisel, 2022), and studying their genetic diversity can inform us about species ecology and life history traits (Ellegren and Galtier, 2016). The knowledge gained from observations of X/Z chromosomes is therefore poorer without the context of the evolution of their Y/W homologs.

A more accessible method for separating and extracting sequences from both sex chromosomes would enable the study of open questions in evolutionary biology broadly, and the evolution of sex chromosomes in particular, including exploring the variation segregating within sex-limited haplotypes (as done in guppies; Almeida et al., 2021), gametolog divergence (as done in grasshoppers and passerines; Jayaprasad et al., 2024; Sigeman et al., 2019), degeneration and selective dynamics of gametologs (as done in larks; Ellerstrand and Hansson, 2026), and co-segregation of the maternally inherited W chromosome and mitochondrion (as done in birds; Berlin and Ellegren, 2001; Gu et al., 2023; Smeds et al., 2015). Such approaches would also facilitate the study of sexually antagonistic mutations segregating on the shared X/Z chromosomes (Eyer et al., 2019; Ruzicka et al., 2020). While Almeida et al., (2021) performed analyses of variation on the sex-limited chromosome using an elaborate approach, their framework is not published in a format that is easily reproducible in other projects.

We have previously applied an early version of our bioinformatic pipeline PhaseWY (https://github.com/sjellerstrand/PhaseWY) to extract W sequences and analyse gametolog dynamics on sex chromosomes in larks (*Alauda*, Alaudidae; Ellerstrand and Hansson, 2026), and in XO and XY grasshoppers (Orthoptera; Jayaprasad et al., 2024). Here, we present PhaseWY as an optimised, user-friendly Snakemake workflow (available on GitHub: https://github.com/sjellerstrand/Snakemake_PhaseWY). PhaseWY uses short-read sequencing data of several females and males and classifies callable genomic regions as sex-linked or autosomal. Highly degenerated regions are classified based on sex differences in alignment depth, where heterogametic genotypes are recoded as haploid for the shared sex chromosome. Non-degenerated regions are classified using a novel approach, employing phasing and haplotype clustering, which has previously been applied by Almeida et al., (2021). Since the number of samples from each sex is known, the expected proportion of sex chromosome haplotypes in the dataset is also known, which can be used to infer sex-linkage. Within these regions, the X/Z and Y/W haplotypes are extracted separately, and genotypes are recoded as haploid in heterogametic individuals. What makes our approach unique is the strict and explicit classification, extraction, subsetting and ploidy-correction of sex-linked genotypes. The output is summarised in vcf and bed format, which can be directly used in downstream analyses with commonly used software packages in population genomics and phylogenomics. While strict classification of a sequence may be considered controversial, the method is transparent and outputs detailed information on how each genomic region was classified.

We generate simulated data to benchmark the accuracy of PhaseYW in relation to several factors, such as biological parameters (time since divergence of the sex chromosomes and effective population size), study design (number of sequenced females and males used to infer haplotypes) and parameter settings for the identification of sex-linkage through haplotype clustering (including window size and minor allele frequency filter, and choice of haplotype clustering model). We then evaluate the pipeline on the Skylark (*Alauda arvensis*) and Raso lark (*A. razae*), two species with several strata of various ages, as well as extreme differences in demographic history and background genomic diversity. Finally, we use the empirical datasets to showcase information outputted by the pipeline, as well as illustrate how the output can be applied to downstream analyses.

## Implementation

PhaseWY is written as an automated Snakemake workflow (Mölder et al., 2021; **Figure 1**). To run the pipeline, the sex-determination system (XY or ZW) and the sex of each sequenced individual is required. If this is unknown, pipelines such as FindZX can be used to infer the sex-determination system, the sex of respective individual, and the genomic regions of interest. PhaseWY is also a dedicated phasing pipeline and can be used as such without identifying sex-linkage. PhaseWY can also be used only for inferring callable regions of the genome and/or as a general phasing pipeline (without identifying sex-linkage). If only a subset of contigs is of interest, it is possible to limit the analysis to those by providing a list of contigs.

**Figure 1.**
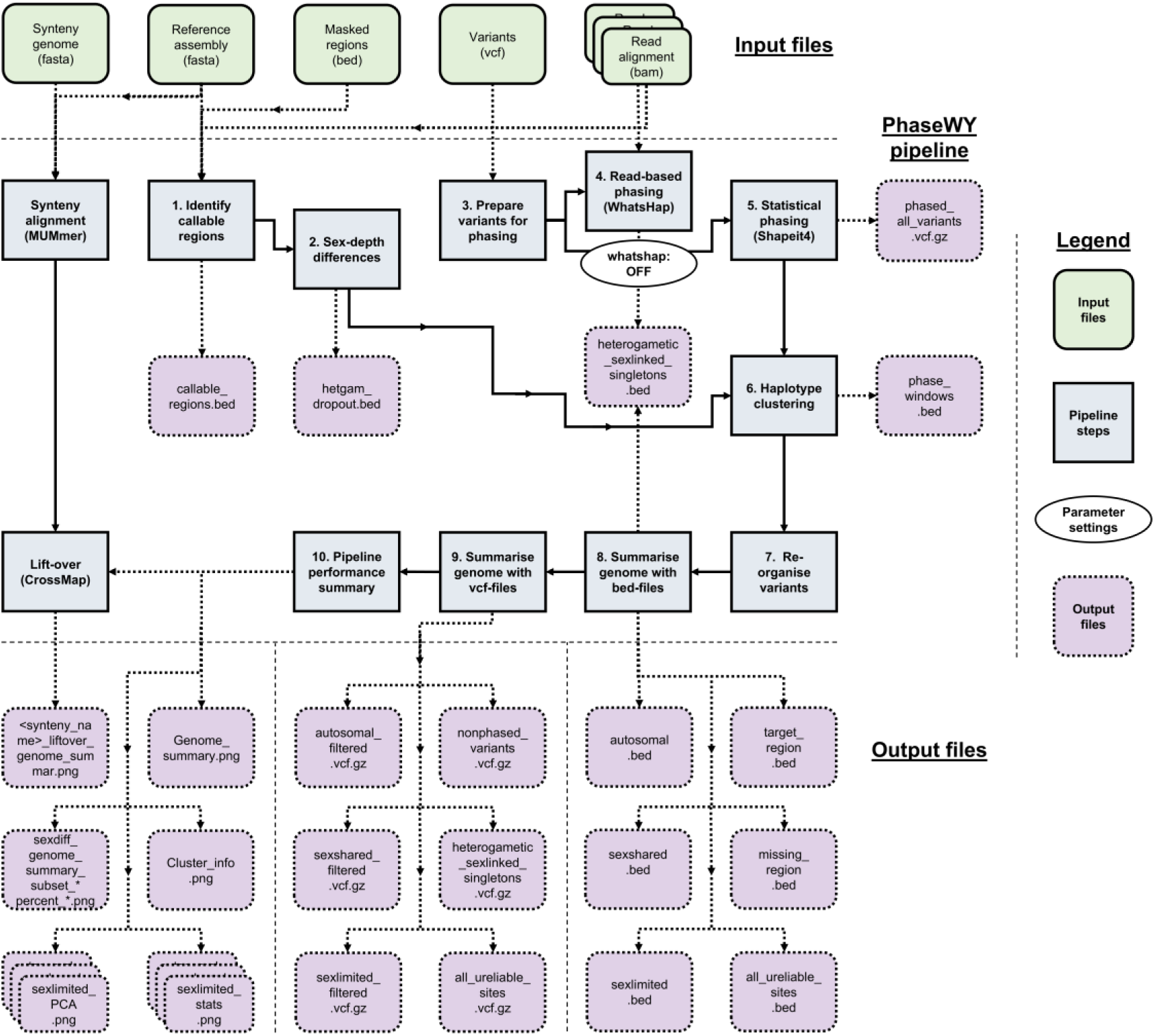
PhaseWY flowchart. An overview of the PhaseWY workflow. The input required include filtered variants of females and males (multi-individual vcf), corresponding read alignments (bam) and genome assembly (fasta). In addition, a file of sites or regions to mask and exclude can be provided (bed). For visualisation of classification along the genome of another species, a synteny genome can be provided. The pipeline performs 10 key steps. Read-based phasing in Step 4 can be disabled. The final output includes a summary of classification across the genome in bed-files, autosomal, X/Z, Y/W and unreliable variants in vcf-files. Furthermore, the classification along the genome is visualised in various plots output in png-format. Once the pipeline is configured, it is automated and can be run with Snakemake using a single command. Furthermore, the user can run a subset of the pipeline to retain callable regions in the genome (Step 1), to retain sex-depth differences across the genome (Step 1-2), or to produced phased data (Step 3-5). Several steps will parallelise over contigs or individuals in an HPC environment. For details on each step, rule and file, see supplementary material.

## Input data

### Reference assembly and contig list

PhaseWY relies on Y/W sequences being aligned to their homologous X/Z reference. Therefore, the reference assembly and alignment should be based on the homogametic sex, or Y/W contigs should be excluded prior to alignment. Users may optionally provide a list of contigs to be analysed. This allows, for example, restriction to known or suspected sex-linked contigs, exclusion of very short contigs, or inclusion of all contigs for a genome-wide exploratory analysis (see also “Masked regions”).

### Read alignments

Bam-files are used to assess callable regions, infer sex differences in alignment depth, and perform read-based phasing for each individual. Bam-files must contain an @RG header with an SM-tag matching the corresponding sample name. The pipeline uses all reads present in the bam-files. Therefore, users are advised to filter the bam files to remove duplicate reads, low-quality, or improper paired reads (e.g. using samtools -f2 -F260 -q20).

### Variants

The vcf-file contains all individuals to be included in the analysis, and all variants to be classified. Biallelic and multiallelic SNPs, indels and MNPs are supported, as are monomorphic sites. Because the provided variants are used to infer phase and haplotype clustering, and because more variants take longer to process, a properly filtered vcf-file is recommended before running the pipeline. However, the filtering step needs consideration, because it might affect the outcome of the analysis. For example, an allelic-balance filter may not be suitable, since it is likely that alignment biases occur between X/Z and Y/W sequence to the homologous X/Z reference. Likewise, excess heterozygosity is expected in the sex-linked region of the heterogametic sex, and too stringent filters on heterozygosity or HWE should therefore be avoided. Furthermore, the heterogametic sex will experience half the coverage in old, highly degenerated sex-linked regions. Therefore, too high minimum depth filters might filter out X/Z sequences, which could otherwise be extracted for analysis. Missing genotypes will be imputed during statistical phasing. Imputation might be considered controversial in small datasets, or in datasets with high missingness. Therefore, the user needs to consider wether to apply strict missingness filters or not. Finally, even if specific genomic elements such as exons are of interest, flanking regions may still contain information utilised for accurate phasing and should therefore not be filtered until after running the pipeline.

### Sample information

The pipeline requires metadata for each individual included in the analysis. To infer sex-specific patterns, individuals must be annotated as homogametic or heterogametic. PhaseWY accepts only diploid individuals and is not designed to accommodate aneuploidies. Mean alignment depth per individual is used to normalize depth when inferring sex differences in coverage. In addition, file paths to individual bam-files are provided to allow efficient access throughout multiple steps of the pipeline.

### Callable filtering parameters

Callable regions across the genome are identified and retained if meeting a set of criteria: minimum depth, minimum mean depth, maximum mean depth, and missingness. We suggest using the same parameters used for variant filtering, since these filters will likewise be applied to variants provided in the vcf.

### Masked regions (optional)

The pipeline allows masking regions of the genome. For example, repetitive regions might be unreliable and can be inferred with repeatmasker (https://www.repeatmasker.org/). These regions will not be considered callable and will not be analysed for sex differences in alignment depth. Neither will any variants present in such regions be included in the output.

### Syntenic reference genome (optional)

If the genome assembly of the study species is highly fragmented, it is possible to visualise the results across complete chromosomes by providing an available chromosome-level assembly of a closely related species. If a syntenic reference genome from a related species is provided, a genome alignment and lift over of genome classification will be performed. Note that the quality of a cross-species alignment is highly dependent on the genome divergence between the study species and the syntenic species.

### Pipeline steps

Various steps of the pipeline alter between parallel and separate operations per contig or individual and goes through several key steps (see supplementary material for details).

### Step 1: Identify callable regions

The alignment depth of each individual and at every site of the contig is evaluated, and sites are retained as callable if passing the filtering criteria thresholds (minimum depth, minimum and maximum mean depth, and proportion of missing data). The sites are then merged into coherent regions, and any region provided in the mask file is excluded.

### Step 2: Identify sex-linked regions based on differences in sequence depth

In highly degenerated sex-limited regions, few or no Y/W reads will align to the homologous reference. In such situations, the heterogametic alignment depth is expected to be about half the mean individual depth across non-sex-linked regions of the genome. The read depth in every individual and at every callable site of the contig is divided by the mean depth of the individual to produce within-individual normalised depths at each site. The average normalised depth at each site is then calculated for each sex. In autosomal regions, the depth of heterogametic individuals divided by the depth of homogametic individuals should be close to 1. In drop-out regions, it should be close to 0.5. Noise in alignment will create two overlapping distributions around those values. We use a threshold of 0.75, and any site with a ratio lower than this value will be classified as sex-linked. These values are also averaged in sliding windows for plotting purposes (same parameter settings as in Step 6). If the threshold needs adjustment, this can be set by the user. However, this requires a complete rerun of the pipeline. A subset of 0.01 % of callable sites across the genome will be retained to produce distribution plots of this ratio in genomic regions of various classifications. The number of sites to subsample can be set by the user.

### Step 3: Prepare variants for phasing

The input vcf is filtered based on the provided contig list, depth and missing-data thresholds, and any variants specified in the masking file. Multiallelic sites are split into biallelic records to ensure compatibility with downstream phasing software. If a contig does not contain at least one biallelic sites with more than one heterozygous genotype (an issue that may occur for small contigs in highly fragmented assemblies), that contig is excluded from further analysis, as it cannot be statistically phased. Variants located on such contigs are retained in the output but are reported as unphased.

### Step 4: Read-based phasing

Variants are first subset into individual vcf-files, after which read-based phasing is performed for each individual using WhatsHap (Martin et al., 2016). Bam-files are used identify reads and read-pairs spanning at least two heterozygous variants within an individual, resulting in phase-sets that span as many variants as can be observed in physically link on the same contig. These phase-sets are subsequently used to inform statistical phasing. Read-based phasing can therefore improve phasing accuracy in small datasets and enables phasing of singleton alleles when they occur within a phase-set. After read-based phasing, all individuals are merged into a single vcf-file. Because singletons not included in any phase-set are randomly assigned to haplotypes during statistical phasing, such sites are considered unreliable. Consequently, heterogametic-specific singletons lacking phase-set information and located in regions classified as sex-linked are recorded for posterity.

This step is the most computationally demanding component of the pipeline and is strongly influenced by bam-file size, contig length and overall genome size, and the level of heterozygosity (whether in autosomes or inflated in sex-linked regions of heterogametic individuals). Nevertheless, read-based phasing can be critical for inferring phase in small datasets, resolve haplotypes in populations with large effective population sizes, and provides the only opportunity to correctly phase singletons. Users may therefore choose to disable read-based phasing as a strategic option for larger datasets. If read-based phasing is not applied, all heterogametic singletons in sex-linked regions are treated as unreliable.

### Step 5: Statistical phasing

Statistical phasing is performed using SHAPEIT4 (Delaneau et al., 2019), which incorporates phase-sets generated by WhatsHap to inform haplotype inference. After phasing, multiallelic variants are reconstructed from split biallelic records into their original multiallelic representation.

SHAPEIT4 imputes missing genotypes based on haplotype information. Because imputation can be undesirable in small datasets or datasets with high level of missingness, users should consider applying appropriately stringent missingness filters prior to phasing, depending on their study design. Finally, singletons that are not included in any phase-sets are randomly assigned to haplotypes during statistical phasing and are therefore considered unreliable (see Step 4).

### Step 6. Haplotype clustering based on sex-linkage

If phasing of sex-linked regions is successful, haplotypes are expected to cluster into X/Z and Y/W haplotypes. Given the number of females and males in the dataset, there is a predefined expectation for the number of haplotypes forming a larger X/Z cluster and a smaller Y/W cluster. This expectation allows inference of whether a genomic region is sex-linked or autosomal based on the haplotype-cluster composition. For regions inferred as sex-linked, the corresponding sex-limited Y/W sequence can subsequently be extracted.

Only some biallelic variants are used for haplotype clustering. Singletons should be excluded (minor allele count = 1), as they provide no information for clustering, are more likely to represent erroneous calls, and may not have been correctly phased (see Step 4). Optionally, a stricter minor-allele count threshold can be applied to match the expected frequency of heterogametic chromosomes in the dataset, which may improve accuracy in datasets with high background genomic diversity. Note that variants excluded from clustering are not removed from the final output, as their phase information is retained. They are thereby extracted later together with their corresponding haplotypes.

Because phase-switch errors can occur during phasing and because sex-linkage may not span entire contigs (e.g., at PAR borders), sex-linkage is evaluated using a sliding-windows approach across the genome. For each window containing at least one variant, haplotypes are forced into two clusters using k-means clustering with 10 random sets. As there are always fewer Y/W than X/Z chromosomes when both sexes are present, haplotypes in the smallest cluster are inferred as putative Y/W haplotypes. A window is classified as sex-linked only if the following criteria are met, evaluated sequentially: 1) only haplotypes from heterogametic individuals are present in the smallest cluster; 2) at most one haplotype per heterogametic individual is present in the smallest cluster; and 3) the size of the smallest cluster matches the expected number of Y/W haplotypes. If these criteria are met, haplotypes in the smallest cluster are annotated as sex-limited Y/W. By default, clustering is based on absolute (Hamming) distances between haplotypes. Alternatively, distances can be weighted by the inverse minor-allele frequency, increasing the influence of rare variants that are more likely to share recent identity by decent.

Uncertainty may arise when an autosomal window borders a sex-linked window, making it ambiguous whether variants are autosomal or sex-linked, and where the exact border between autosomal and sex-linkage should be placed. In such cases, the putatively autosomal window is evaluated for sex-specific depth difference to verify that it does not represent a X/Z region lacking a Y/W sequence. Similar uncertainty can also arise from phase-switch errors affecting the sex-limited haplotype within a series of sex-linked windows. Because a sliding-window approach is applied, all windows overlapping regions of uncertainty are recorded for posterity. Variants located within individual-specific phase-switch regions are not extracted for that individual, as their haplotype assignment cannot be reliably inferred.

The default window size is 10 000 bp with a step size of 2 500 bp, both of which can be adjusted by the user. Larger windows reduce runtime but may be problematic in the presence of frequent phase-switch errors and provide less precise localisation of sex-linkage boundaries. Conversely, very small windows may fail to capture weak signals and can result in fragmented and spurious autosomal classifications.

Two primary output files are generated. The first contains per-window statistics, including classification (sex-linked or autosomal, and the criterion by which it was classified), the number of haplotypes present in the smallest and largest cluster (**Figure 2B**), total and between-cluster sums of squares (**Figure 2C**), cluster-specific sums of squares of the smallest and largest cluster, variant density, sex differences in alignment depth (**Figure 2D**) and heterozygosity (**Figure 2E**; not used for classification of sex-linked variants), and the inferred Y/W haplotype for each heterogametic individual (or “Unknown” if autosomal). The second file provides a simplified summary of coherent regions classified as autosomal, sex-linked or unknown. In addition, two bed-files are produced for each heterogametic individual, specifying which region of the genome correspond to the sex-limited haplotype (“left” or “right” side of the pipe in a phased genotype).

**Figure 2.**
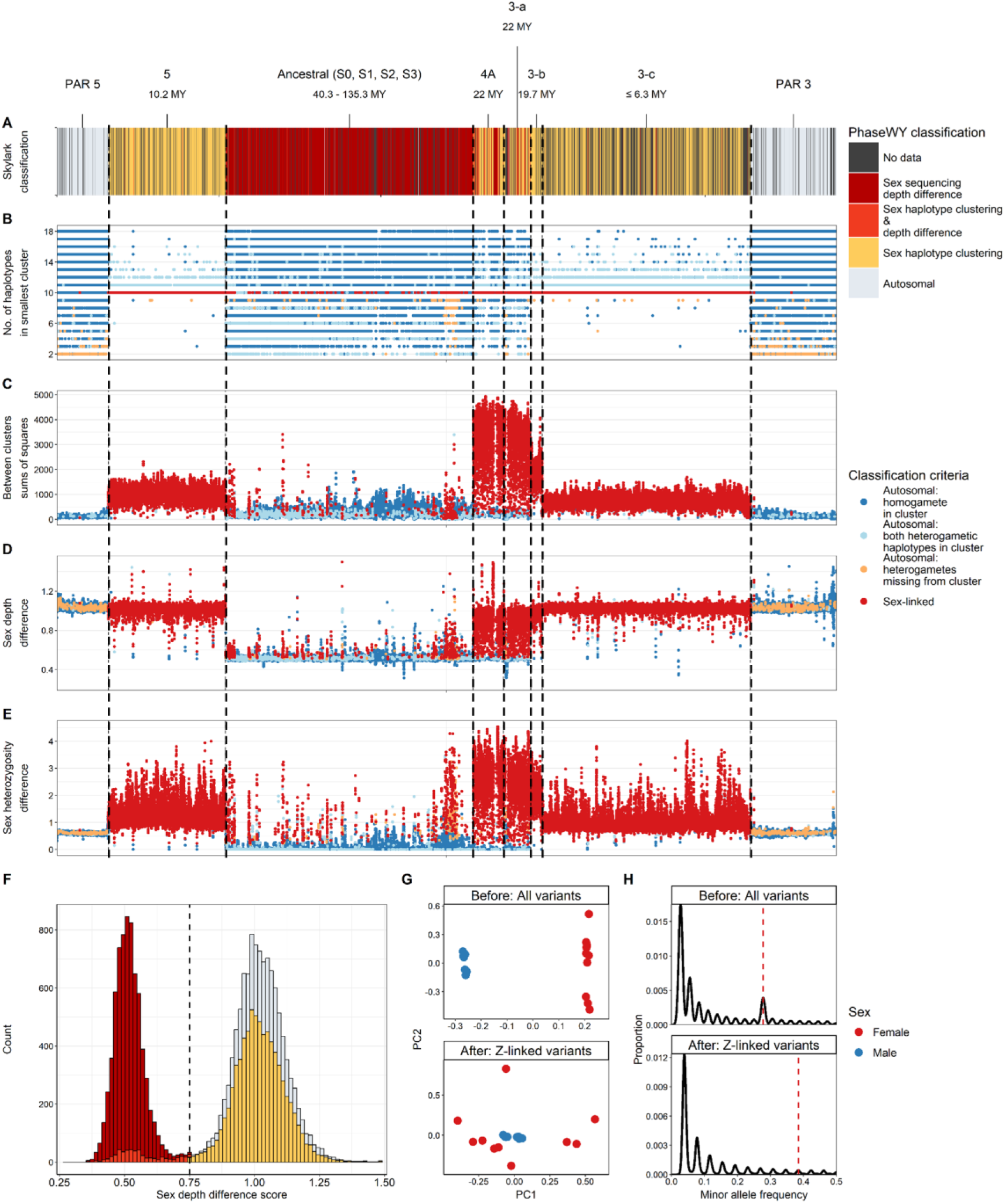
PhaseWY classification statistics. PhaseWY outputs several statistics and plots for interpretation and troubleshooting of the classification of sex-linkage. Haplotypes in a window are forced into two clusters with k-means, and are then evaluated against three key criteria to classify the window as sex-linked: i) homogametic individuals are not allowed in the smallest cluster, ii) two haplotypes are not allowed from the same individual in the smallest cluster, and iii) all heterogametic individuals must be present in the smallest cluster. **(A)** Final classification along the Skylark sex chromosome. **(B)** The number of haplotypes in the smallest cluster. **(C)** Between cluster sums of squares, representing the genetic distance between X/Z and Y/W haplotype clusters. **(D)** Sex difference in alignment depth (female/male). In windows with sex-depth differences, classification is often not possible due to an absence of Y/W sequences. Therefore, Y/W sequences cannot be extracted from these regions, and only recoded, haploid X/Z genotypes in the heterogametic sex will be available. **(E)** Sex differences in heterozygosity (female/male). Not part of the classification process, but presented due to its high sensitivity for detecting young sex-linked regions. **(F)** Distribution of sex depth difference scores (female/male). The distribution of each classification category is output with the user set threshold for sex-depth differences (default 0.75). **(G)** Principal Component Analysis (PCA) results showing PC1 and PC2 for males and females based on all variants before applying PhaseWY, and of Z-linked variants after classification and data subsetting with PhaseWY. **(H)** Minor allele frequency of Skylarks (sexes combined) based on all variants prior to PhaseWY processing and of Z-linked variants following genome classification and data subsetting with PhaseWY. Dashed red line indicates the frequency where W-linked variants may inflate the distribution if present in the dataset, as is observed in the dataset including all variants prior to PhaseWY processing.

Because several parameters can be adjusted in this step – including minor-allele count filter, clustering model, and window and step size – some optimisation may be required. Provided that intermediate files are retained, this step and all subsequent steps can be rerun using alternative parameter settings, with output written to separate folders.

Finally, the presence of homozygous genotypes for both alleles among heterogametic individuals in sex-linked regions may indicate X/Z-linked polymorphisms located in a degenerated region (see Step 2). Alternatively, it may reflect incomplete lineage sorting between the sex chromosomes. As such sites may be unreliable, they are recorded for posterity.

### Step 7. Re-organise genotypes according to genomic region

In this step, all variants are reorganised and re-genotyped according to their inferred genomic classification. Variants located in autosomal regions are output unchanged. Variants in sex-linked regions are separated into two files representing the shared sex chromosome (X/Z) and the sex-limited chromosome (Y/W). For the shared sex chromosome, homogametic individuals are reported as diploid, whereas the heterogametic individuals are reported as haploid. For the sex-limited chromosome, heterogametic individuals are reported as haploid. Sex-linked regions are handled differently depending on whether they were identified through haplotype clustering or though sex-difference in alignment depth.

For regions detected through haplotype clustering, genotypes for each heterogametic individual are first split into two vcf-files corresponding to the “left” and the “right” haplotypes. Information from the haplotype-clustering step is then used to reorganize the genotypes into one file per sex chromosome. Any genotypes that were also detected through sex-specific differences in depth are considered unreliable and are therefore excluded from the sex-limited (Y/W) file.

In regions detected exclusively through a sex-specific difference in alignment depth, homozygous genotypes observed in heterogametic individuals are assumed to represent haploid genotypes, whereas heterozygous genotypes are assumed to be erroneous. Accordingly, homozygous genotypes are encoded as haploid, and heterozygous genotypes are set to missing.

Finally, if identical variants are observed on both chromosomes, this may indicate unsuccessful phasing or reflect incomplete lineage sorting between the sex chromosomes. Because such sites may be unreliable, they are recorded for posterity.

### Pipeline output

The pipeline outputs classified and reorganised variants and reports statistics from the analysis in multiple formats.

### Step 8. Summarise genome with bed-files

The pipeline primarily summarises genomic information in bed-format. These files contain information on which regions are callable or not, and whether they are classified as sex-linked, autosomal or unknown. Sex-linked regions are additionally reported in separate files according to how they were detected. An additional file provides per-window output from the haplotype clustering analysis, including the number of variants analysed per window, cluster sizes inferred from k-means, within- and between-cluster sums of squares, and whether a transition between autosomal and sex-linked region was detected. For windows classified as sex-linked, the inferred sex-limited haplotype is reported for each heterogametic individual. For windows classified as autosomal, the criterion supporting this classification is recorded. Finally, all potentially unreliable sites are reported in separate files according to the criterion by which they were flagged.

### Step 9. Summarise genome with vcf-files

The pipeline summarises genomic variants in vcf-format. One output file contains all phased variants without any further modifications or filters. This file may be informative for investigating the PAR boundary, since they will span the both the sex-linked and PAR with individual haplotype information. However, the correspondence between haplotypes and specific sex chromosomes is not explicit in this file, but can be inferred from the output of Step 6. Additional files generated contain autosomal variants, variants from the shared sex chromosome (with heterogametic individuals as haploids), and variants from the sex-limited chromosome (with heterogametic individuals as haploids). Potentially unreliable sites are reported in separate files according to the criterion by which they were flagged. Note that the pipeline outputs both unfiltered files (including all sites) and files filtered to exclude all flagged unreliable sites. The choice of which file to use for downstream analyses, or whether to exclude only specific classes of unreliable sites, is left to the user. Such a decision may be guided by the pipeline statistics reported in Step 10 for each output file.

### Step 10. Pipeline accuracy summary

For each generated vcf-file, several statistics are produced and visualized to facilitate evaluation of pipeline accuracy. These include the proportion of missing data per site, and minor allele frequency (folded Site Frequency Spectrum; **Figure 2H**). Distributions of individual-specific metrics are also generated, including the inbreeding coefficient (F_IS_; **Figure S1**), and the proportion of missing data per individual. Because some genotypes are haploid, certain statistics – for example inbreeding coefficient– are not applicable to all files. Plink v.1.90b4.9 (Purcell et al., 2007) is used to linkage-prune the data using 50 kb windows, a step sizes of 10 kb, and an r-squared value of 0.1, and principal components are then calculated from the remaining pruned set of independent variants (**Figure 2G**).

The success of variant classification can be evaluated by inspecting diagnostic patterns. For example, if the vcf file containing all variants shows sex-specific clustering in the PCA, this pattern should not be present in the PCA of autosomal variants, nor in that of the X/Z variants (although haploids are expected to show increased dispersion in PC space due to higher variance compared to diploids; Zabel et al., 2025; **Figure 2G**). Similarly, the folded Site Frequency Spectrum may exhibit an excess of variants corresponding to the frequency of the sex-limited chromosome; a pattern that should not be present after classification and subsetting of regions (**Figure 2H**). In addition, sex-bias patterns may be observed in the inbreeding coefficient, as heterogametic individuals tend to have inflated heterozygosity and thus reduced F_IS_. Such bias should not be evident for autosomal variants (**Figure S1**). All these summary plots are produced for each category of potentially unreliable variants. Users may therefore inspect these plots for signatures of sex-linkage and decide whether specific classes of unreliable sites should still be retained for downstream analyses.

A summary of the haplotype clustering performed in Step 6 is presented as Manhattan plots (**Figure 2B-E**). These plots display sums of squares of clusters (total, between clusters, largest-and smallest cluster), the proportion of haplotypes assigned to the smallest cluster, sex difference in heterozygosity and alignment depth, and variant density. Data points are coloured according to their classification as sex-linked or by the criterion under which they were classified as autosomal. Note that sex-linked regions identified solely through sex-specific depth differences will typically appear as autosomal in these plots.

In addition, a single bed-file is produced spanning the analysed genome, in which each region is classified as “Autosomal”, “Sex depth difference”, “Sex haplotype clustering”, “Sex haplotype clustering & depth difference”, or ”Missing data”. This file is used to visualise classification across the reference assembly (**Figure 2A**). If a synteny genome was provided, an equivalent plot is generated in the coordinate system of the synteny species, based on synteny alignment and lift-over of regional classifications; non-aligned regions are also visualised in this plot. Finally, sites in regions of different classifications are visualised as distributions of sex-depth ratios inferred in Step 2 (**Figure 2F**).

### Parameter refinement and troubleshooting

Several parameters can be adjusted to improve pipeline accuracy, including the threshold used to classify sex-linked regions based on sequence-depth difference in Step 2, whether read-based phasing is enabled or disabled in Step 4, and multiple parameters governing haplotype clustering in Step 6. In particular, users may wish to explore alternative parameter setting for the haplotype clustering step. For this reason, Step 6 and all subsequent steps can be rerun with different parameter settings and output written to separate folders, provided that intermediate files from previous steps are retained. However, Steps 1–5 are relatively computationally intensive, and any changes to parameters affecting these steps require a complete rerun of the pipeline (see supplementary section “Parameter refinement” for more details on these parameter choices).

It is also important to note that, whereas sex-linkage inference based on sequence-depth differences is relatively robust to misclassification of sex or aneuploidy in a small number of individuals, haplotype clustering is not and requires accurate knowledge of the frequency of sex chromosomes in the dataset. Consequently, if known sex-linked regions are not identified as expected, this issue can be possible to troubleshoot by inspecting the pipeline summaries from Step 10 (see supplementary section “Troubleshooting” for suggestions on how to troubleshoot common issues).

## Results and discussion

### Simulated datasets

We performed individual-based forward simulations using SLiM 4.2.2 (Haller and Messer, 2023). We simulated 200 kb neutrally evolving autosomes in a diploid population with constant population size and non-overlapping generations. In the first generation, half of the chromosome was converted into a XY sex-linked through complete recombination suppression. Simulations were ran at a range of different constant effective population sizes (Ne: 100, 1,000, 10,000, 100,000, 1,000,000). To investigate the effect of time since recombination cessation, samples were taken after different numbers of generations within the same simulation (Generation: 1,000, 10,000, 100,000, 1,000,000, 10,000,000).

For each simulation, we simulated short-read sequencing data for 10 females and 10 males using NGSNGS v.0.9.2 (Henriksen et al., 2023). Reads were aligned to a separate homogametic female reference sequence, after which variants were called and quality filtered. Each simulation was subsequently analysed with PhaseWY to benchmark accuracy across a range of parameter settings. These included various window sizes for the haplotype clustering algorithm (100, 1,000, 10,000), filtering the input vcf by minor allele count prior to cluster (1, 5, 9), enabling or disabling read-based phasing with WhatsHap (OFF, ON), using alternative distance metrics between haplotypes for the k-means clustering model (Hamming, Inverse MAF), and varying the number of female–male pairs (2, 4, 6, 8, 10).

We then quantified the proportion of sites within the sex-linked region showing a sex-depth difference below 0.75, as well as window-averaged differences in sex-specific heterozygosity in both the pseudoautosomal and sex-linked region. In addition, we calculated the total number of autosomal, X-linked, and Y-linked genotypes that were successfully and falsely extracted by PhaseWY. These did not include Y-linked variants at sites with a sex-depth difference below 0.75.

The accuracy of the pipeline was evaluated as the proportion of variants classified as true positives (Sensitivity), and the proportion of true positives out of all both true and false positives (Precision; see supplementary section 2.1 for details). Rather than focusing on individual parameter estimates, we summarise the dominant effects and their practical implications below. An overview of tested factors and their main effects on sensitivity and precision at realistic parameter space is provided in **Figure 3, S13**, with full statistical details reported in **Figure S4, S8, S9, S12** and **Table S1-S6**.

**Figure 3.**
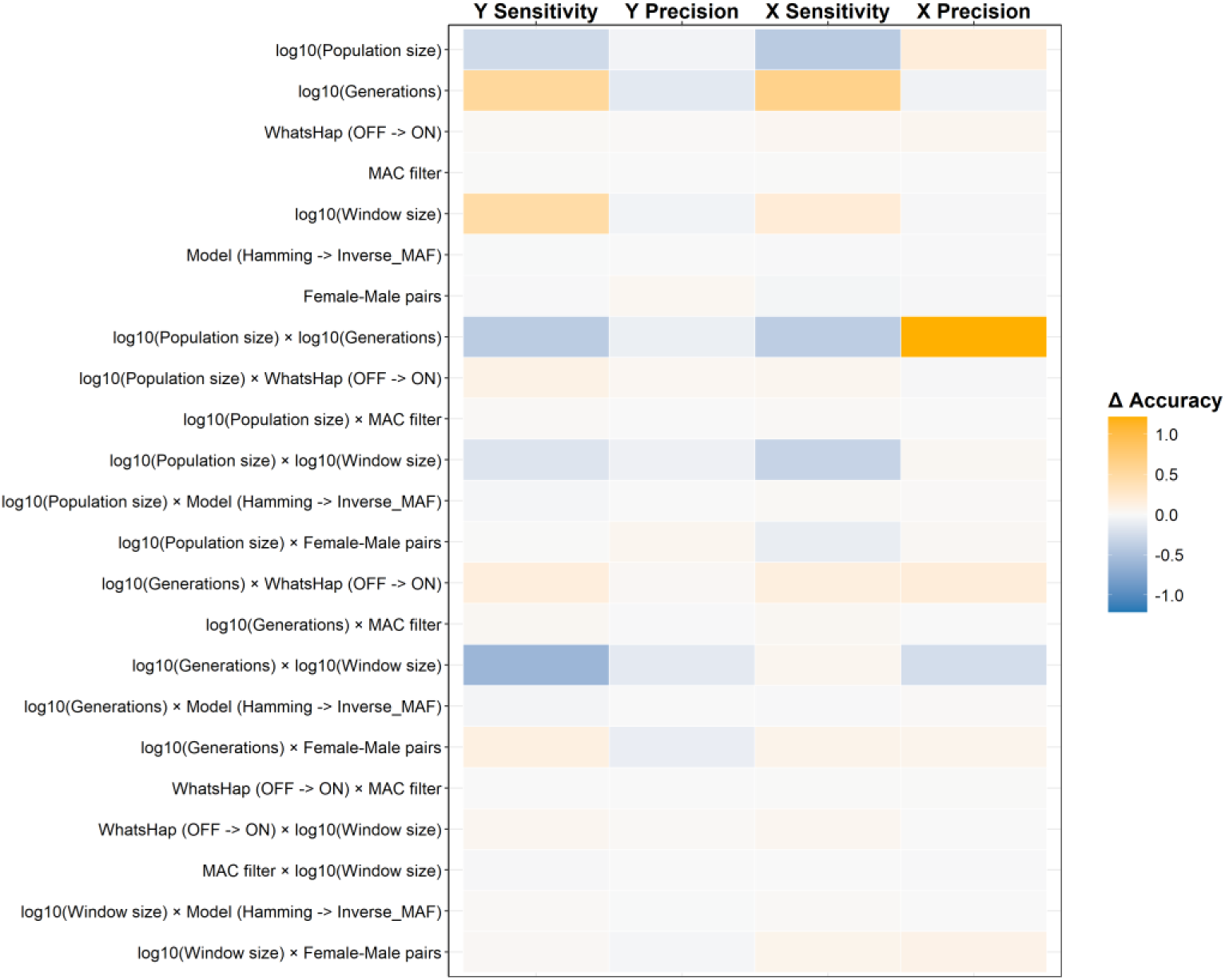
Average predictive contrasts on simulated data. The average change in accuracy between the minimum and maximum value tested for respective parameter. Orange values represent an increase in accuracy, and blue values represent a decrease. Accuracy is shown for Y and X linked markers, and measured as the proportion of variants classified as true positives (Sensitivity), and the proportion of true positives out of all both true and false positives (Precision).

### Detecting sex linkage across evolutionary time

Different signals of sex linkage were informative at different evolutionary timescales. Sex-specific differences in heterozygosity emerged relatively early after recombination suppression and were detectable after approximately 10,000 generations. However, in populations with large effective population sizes, this signal was delayed and only became apparent after substantially longer evolutionary times, reflecting masking by high background genetic diversity (**Figure 4A, S2; Table S1**). In contrast, sex-specific differences in alignment depth required much longer timescales to become detectable, typically only after 1,000,000-10,000,000 generations. These depth-based signals were therefore most informative for older, more degenerated sex-chromosome systems (**Figure S3; Table S2**). Together, these results indicate that heterozygosity-based signals are most powerful for identifying very young sex chromosomes, whereas depth-based signals are better suited for older systems.

**Figure 4.**
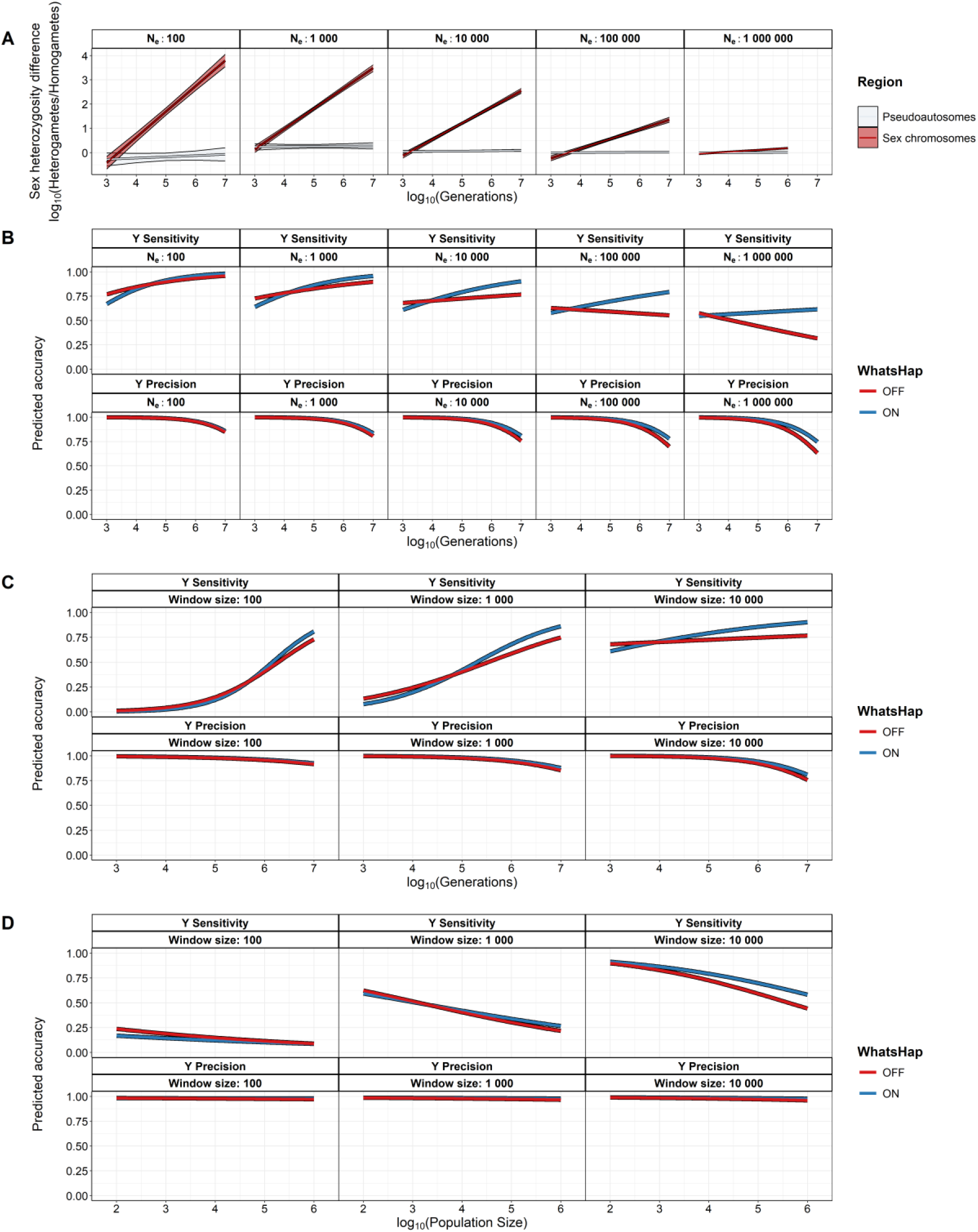
Statistical analyses on simulated data. **(A)** Sex difference in heterozygosity at pseudoautosmal and sex chromosomal regions as a function of generations and effective population size. **(B)** Sensitivity and precision in retaining Y-linked variants as a function of population size, generations, and enabling or disabling WhatsHap for read-based phasing. Values are predicted with window size = 10 000, and MAC = 1. **(C)** Sensitivity and precision in retaining Y-linked variants as a function of window size, generations, and enabling or disabling WhatsHap for read-based phasing. Values are predicted with population size = 10 000, and MAC = 1. **(D)** Sensitivity and precision in retaining Y-linked variants as a function of window size, population size, and enabling or disabling WhatsHap for read-based phasing. Values are predicted with generation = 100 000, and MAC = 1.

### Effective population size

Larger effective population sizes reduce Y sensitivity (**Figure 4B**). Larger populations also reduce X sensitivity, but X precision improves in large populations, especially in old sex chromosome systems (**Figure S6**). Enabling read-based phasing with WhatsHap generally improve accuracy in large populations with rapid linkage decay (**Figure 4B, 4D, S5, S7**). Increasing window size can reduce overall accuracy in large populations, likely due to inefficient statistical phasing when linkage decays quickly rather than because of high background genetic diversity. However, WhatsHap improves the construction of longer, more coherent haplotypes, thereby allowing the use of larger window sizes in larger populations (**Figure 4D, S7**).

### Time since recombination cessation

Evolutionary time generally improves the sensitivity in Y linked markers, but reduces Y precision in very old sex chromosome systems (**Figure 4B-C**). Likewise, X sensitivity improves, but precision is generally reduced with time (**Figure S5, S6**). Enabling read-based phasing with WhatsHap generally improve accuracy in older sex chromosome systems (**Figure 4B-C, S5, S6**). Increasing window size improves Y sensitivity in young systems, and X sensitivity overall, while it somewhat reduces Y and X precision in old systems (**Figure 4C, S6**), likely due to increased noise from patchy alignments in degenerated regions.

### Read-based phasing

Enabling read-based phasing with WhatsHap appears to overall improve accuracy, especially in old systems, and enables the use of larger window sizes through the construction of longer, more coherent haplotypes (**Figure 4C-D, S6, S7**). Notably, in large populations, Y sensitivity appears to improve with time when WhatsHap is enabled (**Figure 4B**).

### Window size

Larger window sizes generally improve accuracy. This effect is especially seen in Y and X sensitivity in small populations, while it has a little effect in large populations (**Figure 4C-D, S6, S7**). However, larger window sizes appear to slightly reduce Y precision in old systems (**Figure 4C**). Moreover, read-based phasing with WhatsHap appear to benefit by larger window sizes (**Figure 4C-D, S6, S7)**.

### Very small sample sizes

We further evaluated accuracy when only two or three individuals were available (**Figure S12, S13**). PhaseWY requires at least three individuals for haplotype-based classification, whereas a simpler companion Python script can be used to extract Y-linked variants from a single female–male pair when sex-linked regions are known a priori. Y sensitivity was greatly improved in young sex-chromosome systems with the simpler extraction approach relative to PhaseWY with three individuals, while both approaches performed similarly in old systems. PhaseWY performs best in young systems when using one additional heterogametic male, while older systems benefit from using one additional homogametic female (**Figure S14, S15**).

Accuracy for X-linked variants was more context-dependent on population size and evolutionary time. Overall, the simpler extraction approach had higher X sensitivity in young sex-chromosome systems, and higher X precision in small to medium population sizes. Importantly, PhaseWY greatly outperforms the simple extraction approach in terms of X sensitivity in older systems, especially using one additional female. Again, PhaseWY performs best in young systems when using one additional heterogametic male, while older systems benefit from using one additional homogametic female (**Figure 4D**).

### Non-influential parameters

Three of the tested parameters had little to no impact on accuracy. Minor allele count filter and clustering model appears to have no impact on any marker or accuracy metric. Increasing the number of female–male pairs from two to ten has a relatively minor impact on pipeline accuracy, but a slight increase in Y precision (**Figure S11**). Increasing the number of female–male pairs improved the sensitivity of Y and X in old systems and large window sizes, as well as slightly improved X precision when larger window sizes were used (**Figure S10, S11**).

### Practical recommendations

Taken together, these simulations highlight that evolutionary and demographic context dominate accuracy, whereas most user-defined parameters have comparatively minor effects. However, larger window sizes and read-based phasing are generally recommended. With limited individuals, additional heterogametic individuals are preferable in young systems, whereas additional homogametic individuals can be advantageous in older systems. However, our simulations suggest that increasing the number of female–male pairs from two to ten has a relatively minor impact on pipeline accuracy. Consequently, user may wish to base decisions about sample size primarily on other considerations, such as the number of individuals or haplotypes required for downstream analyses.

### Empirical datasets

The largest known avian sex chromosomes have been described in the widespread Eurasian Skylark (*A. arvensis*) and the critically endangered Raso lark (*A. razae;* Sigeman et al., 2019)). Larks possess a ZW sex chromosome system comprising several evolutionary strata that vary widely in divergence time, ranging from approximately 6 to 135 MYA (corresponding to ca 3–35 million generations; Ellerstrand and Hansson, 2026). The Skylark and Raso lark diverged around 6.3 MYA, yet differ markedly in geographic range, demographic history and levels of background nucleotide diversity (0.016 vs. 0.0013 in Skylark and Raso lark, respectively). This system therefore provides an informative contrast for assessing how divergence time and background genomic diversity jointly influence phasing and haplotype clustering accuracy. Inspecting the classification output revealed that in the two youngest strata (5 and 3-c), high background genetic diversity in the Skylark interfered with reliable classification in regions of low sex-chromosome divergence, a pattern not evident in the severely bottlenecked Raso lark (**Figure 5**).

**Figure 5.**
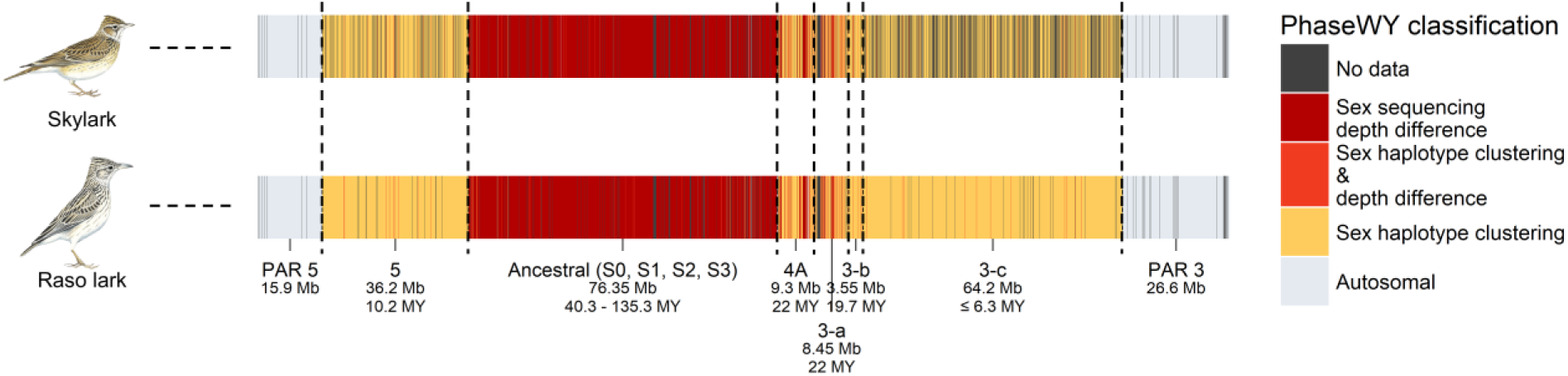
PhaseWY classification of sex-linkage in empirical datasets. Regions are classified as autosomal (light grey), sex-linked (sex depth differences as red; haplotype clustering as yellow; both as orange), no data (shown as black; including non-callable regions, repetitive regions, uncertain phasing), or unsuccessful synteny alignment (shown as white). **(A)** The sex-chromosomes of the Skylark and Raso lark. Strata are annotated by name, size, and age. The Skylark is a widespread and genetically diverse species, while the Raso lark is severely bottleneck and retains little genetic diversity. Despite both datasets being aligned to the Skylark reference assembly, it is evident in the two youngest strata - 5 and 3-c - that high levels of background genetic diversity interferes with classification in low-divergence regions. Illustrations by Tim Worfolk (Birds of the World).

### Downstream applications

We performed a series of downstream analyses using on the lark datasets to illustrate how the pipeline output can be applied in practice **(Figure 6**).

**Figure 6.**
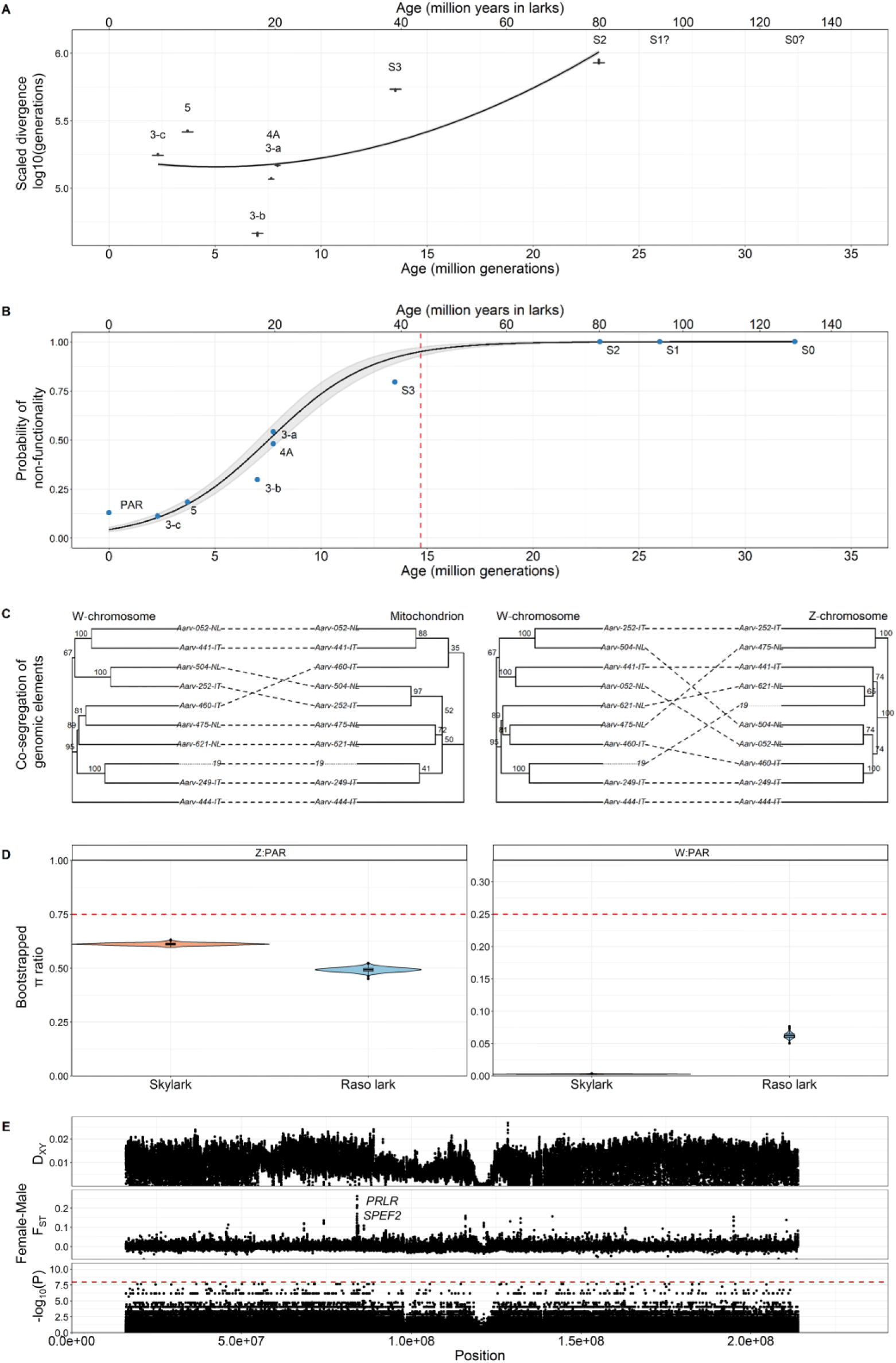
Examples of downstream applications of PhaseWY output. **(A)** 1,000 ultrafast bootstrap estimates of gametolog divergence in the Skylark, halved and scaled by the Great reed warbler mutation rate (µ=7.16×10^−9^ m/s/g) to achieve divergence in generations. Boxplots are ordered by putative age of respective stratum (MG=million generations). The line corresponds to a linear model fit (y∼poly(x,2)). **(B)** Predicted probability of W gametolog non-functionality (± 95% confidence intervals) as a function of stratum age (in million generations, MG). The equivalent age in years is shown on top axis. Blue points represent the observed proportion of non-functional genes on each stratum (and PAR 3 and 5 combined at age zero). The red dashed line indicates the estimated age at which W gametologs reach 95% probability of non-functionality. Reproduced from Ellerstrand and Hansson (2026). **(C)** Co-segregation of Skylark W haplotypes and mitochondrion, including the Z-chromosome as control. **(D)** Bootstrapped nucleotide diversity sex chromosome–PAR ratios (Z:autosomes and W:autosomes, respectively) at “neutral” sites located ≥ 20 kb from any exon for Raso lark (blue) and Skylark (peach). Red dashed lines indicate the expected ratios under mutation–drift equilibrium, even sex ratios and neutral evolution. **(E)** Investigation of sexually antagonistic polymorphisms on Skylark Z using female-male pairwise D_XY_, F_ST_ and GWAS. The scans were performed using a windows size of 10,000 bp and a step size of 2,500 bp. Red dashed line corresponds to the Bonferroni-corrected 0.05 p-value threshold for multiple testing.

### Gametolog divergence in strata of various ages

We investigated the gametolog divergence across strata of different ages by constructing phylogenetic trees for each sex chromosome stratum in the Skylark (**Figure 6A**). Analyses were restricted to CDS regions that were callable on both Z and W, as inferred by the pipeline (i.e., no sex-depth difference). Genes located on the ancestral Z chromosome were further assigned to evolutionary strata by cross-referencing gene names to Xu et al. (2019). Alignments included both variable and invariant sites within callable regions.

For each stratum, we inferred maximum-likelihood phylogenetic trees using IQ-TREE v.2.4.0 (Minh et al., 2020), with 1 000 ultrafast bootstrap replicates (Hoang et al., 2018). Branch lengths from each bootstrap tree were halved to obtain estimates of divergence time and associated confidence, and subsequently scaled using a mutation rate of 7.16×10^−9^ m/s/g, as estimated for the Great reed warbler (Zhang et al., 2023). For the oldest stratum, S0 (32.3 MG/135.3 MY), we did not retrieve sufficient callable W-linked regions nor enough variants to construct a phylogeny. Similarly, we excluded the second oldest stratum, S1 (26.0 MG/92.0 MY), due to limited amount of informative data recovered (19 phylogenetically informative sites and 123 constant sites). This scarcity of data likely reflects extensive Z–W divergence and degeneration of W gametologs over long evolutionary timescales (**Figure 6B**). Overall, estimated divergence times increased with stratum age, although some relative inconsistencies were observed among neo-sex chromosomal strata. The inferred order of sex-linkage across strata is well supported by comparative species evidence (Dierickx et al., 2020; Ellerstrand and Hansson, 2026; Sigeman et al., 2019), but it is important to note that the absolute ages of individual strata are only approximately estimated from the literature (Ellerstrand and Hansson, 2026, supplementary material). Across strata, the scaled divergence estimates were one to two orders of magnitudes lower (10-150x) than the estimated accepted ages of respective stratum. Because these estimates were based solely on CDS regions, they are likely strongly influenced by varying intensities of purifying selection. For example, stratum 3-b exhibited particularly low divergence and retained a relatively high number of functional W gametologs compared to other strata (**Figure 6B**), potentially reflecting relatively stronger purifying selection acting on genes within this stratum.

### Loss of W gametologs over time

We reproduced a subfigure from Ellerstrand and Hansson (2026) in which the probability of W gene-loss over time was modelled (**Figure 6C**). These analyses were conducted using the same individuals as in the present study, but were based on a scaffold-level assembly and an earlier, non-Snakemake implementation of PhaseWY.

Briefly, W gametologs were classified as non-functional based on sex-depth differences and predicted mutational impacts inferred by SnpEff v.4.3t (Cingolani et al., 2012). Gametologs were considered non-functional if they had lost at least one exon, or if an otherwise intact transcript was fixed for one or more loss-of-function mutation (i.e. mutations causing loss of start codons, gain or loss of stop codons, or frameshifts). We then used glmmTMB v.1.1.10 (Brooks et al., 2017) to make generalized linear model (GLMs) with a binomial error distribution and logit link function to test whether the probability of W non-functionality varied with stratum age (million generations, MG), species (Skylark vs. Raso lark), gene haploinsufficiency (where high haploinsufficiency indicates that two copies are required for normal gene function; Collins et al., 2022), gene length, and their interactions. We found that the probability of W non-functionality increased with both stratum age and gene length, whereas haploinsufficiency mitigated the effect of gene length on gene loss. Using median predictor values and averaging across species, we estimated that a typical W gametolog reaches a 95% probability of being non-functional after ∼14.7 MG, corresponding to ∼43.5 MY in passerines and larks.

### Co-segregation of the W chromosome and mitochondrion

We investigated maternal co-segregation of the Skylark W chromosome and mitochondrion, using the Z chromosome as a control (**Figure 6C**). Maximum-likelihood trees were inferred using IQ-TREE v.2.4.0 (Minh et al., 2020), including model selection (Kalyaanamoorthy et al., 2017) with ascertainment-bias correction and 1,000 ultrafast bootstrap replicates. Resulting trees were converted to ultrametric trees using chronos from ape v.5.8.1 (Paradis and Schliep, 2019). Co-segregation patterns were visualised using cophylo from phytools v.2.4.4 (Revell, 2024).

Overall, we observed strong concordance between the W chromosome and mitochondrial phylogenies, consistent with maternal inheritance, although a small number of individuals deviated from this pattern. Such deviations may have biological meaningful explanations. For example, mitochondrial heteroplasmy within an individual – the presence of multiple mitochondrial haploptypes within an individual – can arise through paternal leakage, or potentially through recombination involving either the W chromosome or the mitochondrial genome (Smeds et al., 2015). Alternatively, the observed discrepancies may reflect technical limitations, including limited resolution of within-species phylogenetic relationships. This interpretation is supported by low bootstrap support on some branches, potentially due to limited variation on the W chromosome and the mitochondrion.

### Deviations in nucleotide diversity between autosomes and sex chromosomes

We investigated deviations from the expected ratio of nucleotide diversity (π) between autosomes, the Z chromosome and the W chromosome under neutral evolution, random mating and even sex ratios (A:Z:W = 4:3:1; **Figure 6D**). Nucleotide diversity was estimated for the PAR (pseudoautosomal region), Z and W regions using non-overlapping 10 kb-windows. We applied predefined windows based on callable regions annotated by the pipeline, using only regions which required to be at least ≥ 20 kb from any annotated exon to approximate neutrality.

Across both species, nucleotide diversity (π) was lowest on W, intermediate on Z and highest on autosomes (A/Z/W; Raso lark: 0.0013/6.2×10^-4^/7.8×10^-5^; Skylark: 0.016/0.0099/4.5×10^-5^). The π_Z_/π_A_-ratio was lower in the Raso lark (0.49) than in the Skylark (0.61), and in neither species did the distributions overlap the neutrally expectation of 0.75. In contrast, the π_W_/π_A_-ratio was substantially higher in the Raso lark (0.062) than in the Skylark (0.0028), although both values were far below the expected ratio of 0.25.

These estimates broadly agree with previously reported values based on the same individuals, but derived from a scaffold-level assembly, using strictly autosomal variants, and an earlier, non-Snakemake version of PhaseWY; SJ Ellerstrand et al., unpublished). However, the present analyses yielded generally lower π values across most genomic categories and lower π_W_/π_A_-ratio for both species. Notably, whereas π_Z_/π_A_ in the Skylark was previously indistinguishable from the neutrally expected ratio, it was clearly reduced in the present analysis (Raso lark new/old: 0.49/0.57, Skylark new/old: 0.61/0.75).

The reduced π_Z_/π_A_-ratios in both species may reflect male-biased sexual selection, which is known to reduce effective population size of the Z chromosome. The particularly low ratio observed in the Raso lark likely also reflects historically skewed sex ratios (1.4–3.0 males per female; Donald et al., 2007, 2005, 2003), which would further amplify the effects of male sexual selection. In contrast, the much lower π_W_/π_A_-ratio in the Skylark relative to Raso lark likely reflect stronger purifying selection acting on the completely linked W-chromosome, leading to more efficient removal of deleterious variation (SJ Ellerstrand et al., unpublished).

### Sexually antagonistic polymorphisms on Z

We investigated signs of segregating sexually antagonistic polymorphisms on the Skylark Z chromosome (**Figure 6E**). Specifically, we conducted a genome-wide association study (GWAS) with sex as a response variable and performed pairwise sliding-window analyses of F_ST_ and D_XY_ between females and males using non-overlapping 10 kb-windows. Sliding window analyses were applied to predefined windows based on callable regions annotated by the pipeline, in order to avoid biases associated with missing data.

Across analyses, we detected a single outlying region on the ancestral Z chromosome based on elevated F_ST_. This region overlaps the genes *PRLR* and *SPEF2* within stratum S2, both of which have been associated with traits such as egg production, spermatogenesis, behaviour, immunoregulation and moulting (Elferink et al., 2008) – functions that are plausible candidates for sexual conflict. However, this signal requires further validation, including careful exclusion of potential pipeline artifact. Neither the GWAS nor the Dxy analysis provided additional evidence of sexual polymorphism, likely reflecting the limited statistical power of the small dataset (10 females and 8 males). More generally, sexually antagonistic polymorphisms are expected to be difficult to detect and would require substantially larger sample sizes of individuals with extreme phenotypes.

## Conclusion

The rapid advancement of sequencing technologies and the growing availability of genomic data in public repositories underscore the need for bioinformatic tools that make studies of genome evolution more accessible to a broad research community. At the same time, the genomics era has greatly expanded our understanding of the diversity of sex-chromosome systems across the tree of life. Whereas the identification and analysis of sex chromosomes previously required specialised expertise and equipment, such analyses can now be pursued by most researchers in eco-evolutionary genomics as a complementary component of broader genomic studies.

Despite this progress, the extraction and analysis of sex-limited Y/W sequences remain technically challenging. Here, we present PhaseWY, a Snakemake-based pipeline that uses WGS data from multiple female and male individuals to detect sex chromosomes and extract Y/W sequences. PhaseWY is designed to be straightforward to use, adaptable to a wide range of non-model organisms, and flexible enough to accommodate different study designs and data qualities. By lowering the technical barriers to sex-chromosome analysis, we anticipate that PhaseWY will facilitate investigations into long-standing and emerging questions in evolutionary biology, particularly those concerning the origin, diversification and degeneration of sex chromosomes. Given that sex chromosomes both influence and are affected by evolutionary processes, ecology and life history variation, we hope that PhaseWY will encourage broader exploration of sex-chromosome evolution across diverse taxa.

## Availability of PhaseWY

PhaseWY is available on GitHub (https://github.com/sjellerstrand/Snakemake_PhaseWY) and can be used as a ready-to-run Snakemake pipeline or as a modular code that users may adapt and extend for customised extraction of Y/W sequences. An example config file, contig list and sample information is provided with the pipeline. Moreover, a corresponding example dataset (3.5 Gb) from the Raso lark (*A. razae*) is available at https://doi.org/10.5281/zenodo.19050140, and includes a reference genome assembly, read alignments, variant calls, masked regions and the Great tit (*Parus major*) genome for use as a synteny reference.

## Supplementary Material

Supplementary material is available at the journal website:

- Supplementary Text: Documentation, Material and Methods
- Supplementary Figures S1-S15
- Supplementary Tables S1-S6

## Code availability

PhaseWY and additonal code and scripts used in this study are provided at GitHub: https://github.com/sjellerstrand/Snakemake_PhaseWY.

## Supporting information

Supplementary Text: Documentation, Material and Methods

## Acknowledgements

We want to thank Zachary Nolen for discussions about Snakemake workflows, and Fredrik Andreasson for discussions about statistical analyses. Bioinformatics analyses were performed on computational infrastructure provided by the National Academic Infrastructure for Supercomputing in Sweden (NAISS) at Uppsala Multidisciplinary Center for Advanced Computational Science (UPPMAX). This work was supported by the SciLifeLab & Wallenberg Data Driven Life Science Program, Knut and Alice Wallenberg Foundation (grants: KAW 2020.0239 and KAW 2017.0003), and by the National Bioinformatics Infrastructure Sweden (NBIS) at SciLifeLab.

## Author contribution

SJE and BH conceived the study. SJE developed the code of the pipeline and performed the analyses with input from BH. AC and VEK converted the code into a Snakemake pipeline with input from SJE. SJE and BH wrote the paper with input from AC and VEK.

## References

1. Almeida, P., Sandkam, B.A., Morris, J., Darolti, I., Breden, F., Mank, J.E., 2021. Divergence and Remarkable Diversity of the Y Chromosome in Guppies. Molecular Biology and Evolution 38, 619–633. 10.1093/molbev/msaa257

2. Bachtrog, D., 2008. The Temporal Dynamics of Processes Underlying Y Chromosome Degeneration. Genetics 179, 1513–1525. 10.1534/genetics.107.084012

3. Bachtrog, D., Mank, J.E., Peichel, C.L., Kirkpatrick, M., Otto, S.P., Ashman, T.-L., Hahn, M.W., Kitano, J., Mayrose, I., Ming, R., Perrin, N., Ross, L., Valenzuela, N., Vamosi, J.C., The Tree of Sex Consortium, 2014. Sex Determination: Why So Many Ways of Doing It? PLoS Biol 12, e1001899. 10.1371/journal.pbio.1001899

4. Berlin, S., Ellegren, H., 2001. Clonal inheritance of avian mitochondrial DNA. Nature 413, 37–38.

5. Brooks, M., E., Kristensen, K., van Benthem, K., J., Magnusson, A., Berg, C., W., Nielsen, A., Skaug, H., J., Mächler, M., Bolker, B., M., 2017. glmmTMB Balances Speed and Flexibility Among Packages for Zero-inflated Generalized Linear Mixed Modeling. The R Journal 9, 378. 10.32614/RJ-2017-066

6. Brown, E.J., Nguyen, A.H., Bachtrog, D., 2020. The Y chromosome may contribute to sex-specific ageing in Drosophila. Nat Ecol Evol 4, 853–862. 10.1038/s41559-020-1179-5

7. Bull, J., 1983. Evolution of sex determining mechanisms. The Benjamin/Cummings Publishing Company, Inc., Menlo Park, California.

8. Chandler, C.H., 2017. When and why does sex chromosome dosage compensation evolve? Annals of the New York Academy of Sciences 1389, 37–51. 10.1111/nyas.13307

9. Cingolani, P., Platts, A., Wang, L.L., Coon, M., Nguyen, T., Wang, L., Land, S.J., Lu, X., Ruden, D.M., 2012. A program for annotating and predicting the effects of single nucleotide polymorphisms, SnpEff: SNPs in the genome of Drosophila melanogaster strain w ^1118^ ; iso-2; iso-3. Fly 6, 80–92. 10.4161/fly.19695

10. Collins, R.L., Glessner, J.T., Porcu, E., Lepamets, M., Brandon, R., Lauricella, C., Han, L., Morley, T., Niestroj, L.-M., Ulirsch, J., Everett, S., Howrigan, D.P., Boone, P.M., Fu, J., Karczewski, K.J., Kellaris, G., Lowther, C., Lucente, D., Mohajeri, K., Nõukas, M., Nuttle, X., Samocha, K.E., Trinh, M., Ullah, F., Võsa, U., Epi25 Consortium, Estonian Biobank Research Team, Hurles, M.E., Aradhya, S., Davis, E.E., Finucane, H., Gusella, J.F., Janze, A., Katsanis, N., Matyakhina, L., Neale, B.M., Sanders, D., Warren, S., Hodge, J.C., Lal, D., Ruderfer, D.M., Meck, J., Mägi, R., Esko, T., Reymond, A., Kutalik, Z., Hakonarson, H., Sunyaev, S., Brand, H., Talkowski, M.E., 2022. A cross-disorder dosage sensitivity map of the human genome. Cell 185, 3041–3055.e25. 10.1016/j.cell.2022.06.036

11. Cortez, D., Marin, R., Toledo-Flores, D., Froidevaux, L., Liechti, A., Waters, P.D., Grützner, F., Kaessmann, H., 2014. Origins and functional evolution of Y chromosomes across mammals. Nature 508, 488–493. 10.1038/nature13151

12. Delaneau, O., Zagury, J.-F., Robinson, M.R., Marchini, J.L., Dermitzakis, E.T., 2019. Accurate, scalable and integrative haplotype estimation. Nat Commun 10, 5436. 10.1038/s41467-019-13225-y

13. Dierickx, E.G., Sin, S.Y.W., van Veelen, H.P.J., Brooke, M. de L., Liu, Y., Edwards, S.V., Martin, S.H., 2020. Genetic diversity, demographic history and neo-sex chromosomes in the Critically Endangered Raso lark. Proc. R. Soc. B. 287, 20192613. 10.1098/rspb.2019.2613

14. Donald, P.F., Brooke, M.D.L., Bolton, M.R., Taylor, R., Wells, C.E., Marlow, T., Hille, S.M., 2005. Status of Raso Lark *Alauda razae* in 2003, with further notes on sex ratio, behaviour and conservation. Bird Con. Int. 15, 165–172. 10.1017/S0959270905000134

15. Donald, P.F., de Ponte, M., Pitta Groz, M.J., Taylor, R., 2003. Status, ecology, behaviour and conservation of Raso Lark *Alauda razae*. Bird Conservation International 13, 13–28. 10.1017/S0959270903003022

16. Donald, P.F., Hille, S., Brooke, M.D.L., Taylor, R., Wells, C.E., Bolton, M., Marlow, T., 2007. Sexual dimorphism, niche partitioning and social dominance in the feeding ecology of the critically endangered Raso Lark *Alauda razae*. Ibis 149, 848–852. 10.1111/j.1474-919X.2007.00701.x

17. Elferink, M.G., Vallée, A.A., Jungerius, A.P., Crooijmans, R.P., Groenen, M.A., 2008. Partial duplication of the PRLR and SPEF2 genes at the late feathering locus in chicken. BMC Genomics 9, 391. 10.1186/1471-2164-9-391

18. Ellegren, H., Galtier, N., 2016. Determinants of genetic diversity. Nat Rev Genet 17, 422–433. 10.1038/nrg.2016.58

19. Ellerstrand, S.J., Hansson, B., 2026. Selective regimes and evolutionary dynamics of Z and W gametologs across an expanded avian neo-sex chromosome. Genome Biology and Evolution.

20. Eyer, P.-A., Blumenfeld, A.J., Vargo, E.L., 2019. Sexually antagonistic selection promotes genetic divergence between males and females in an ant. Proc. Natl. Acad. Sci. U.S.A. 116, 24157–24163. 10.1073/pnas.1906568116

21. Fisher, R., 1930. The genetical theory of natural selection. Clarendon Press, Oxford.

22. Graves, J., Wakefield, M.J., Toder, R., 1998. The origin and evolution of the pseudoautosomal regions of human sex chromosomes. Human Molecular Genetics 7, 1991–1996. 10.1093/hmg/7.13.1991

23. Grayson, P., Wright, A., Garroway, C.J., Docker, M.F., 2022. SexFindR: A computational workflow to identify young and old sex chromosomes. bioRxiv. 10.1101/2022.02.21.481346

24. Gu, H., Wen, J., Zhao, X., Zhang, X., Ren, X., Cheng, H., 2023. Evolution, Inheritance, and Strata Formation of the W Chromosome in Duck (*Anas platyrhynchos*). Genome Biology and Evolution 15, evad183. 10.1093/gbe/evad183

25. Haldane, J.B.S., 1922. Sex ratio and unisexual sterility in hybrid animals. Journal of Genetics 12, 101–109.

26. Haller, B.C., Messer, P.W., 2023. SLiM 4: Multispecies Eco-Evolutionary Modeling. The American Naturalist 201, E127–E139. 10.1086/723601

27. Henriksen, R.A., Zhao, L., Korneliussen, T.S., 2023. NGSNGS: next-generation simulator for next-generation sequencing data. Bioinformatics 39, btad041. 10.1093/bioinformatics/btad041

28. Hoang, D.T., Chernomor, O., von Haeseler, A., Minh, B.Q., Vinh, L.S., 2018. UFBoot2: Improving the Ultrafast Bootstrap Approximation. Molecular Biology and Evolution 35, 518–522. 10.1093/molbev/msx281

29. Jayaprasad, S., Peona, V., Ellerstrand, S.J., Rossini, R., Bunikis, I., Pettersson, O.V., Olsen, R.-A., Rubin, C.-J., Einarsdottir, E., Bonath, F., Bradford, T.M., Cooper, S.J.B., Hansson, B., Suh, A., Kawakami, T., Schielzeth, H., Palacios-Gimenez, O.M., 2024. Orthopteran Neo-Sex Chromosomes Reveal Dynamics of Recombination Suppression and Evolution of Supergenes. Molecular Ecology 33, e17567. 10.1111/mec.17567

30. Jeffries, D.L., Lavanchy, G., Sermier, R., Sredl, M.J., Miura, I., Borzée, A., Barrow, L.N., Canestrelli, D., Crochet, P.-A., Dufresnes, C., Fu, J., Ma, W.-J., Garcia, C.M., Ghali, K., Nicieza, A.G., O’Donnell, R.P., Rodrigues, N., Romano, A., Martínez-Solano, Í., Stepanyan, I., Zumbach, S., Brelsford, A., Perrin, N., 2018. A rapid rate of sex-chromosome turnover and non-random transitions in true frogs. Nat Commun 9, 4088. 10.1038/s41467-018-06517-2

31. Kalyaanamoorthy, S., Minh, B.Q., Wong, T.K.F., von Haeseler, A., Jermiin, L.S., 2017. ModelFinder: fast model selection for accurate phylogenetic estimates. Nat Methods 14, 587–589. 10.1038/nmeth.4285

32. Martin, M., Patterson, M., Garg, S., O Fischer, S., Pisanti, N., Klau, G.W., Schöenhuth, A., Marschall, T., 2016. WhatsHap: fast and accurate read-based phasing. bioRxiv. 10.1101/085050

33. Meisel, R.P., 2022. Ecology and the evolution of sex chromosomes. J of Evolutionary Biology 35, 1601–1618. 10.1111/jeb.14074

34. Minh, B.Q., Schmidt, H.A., Chernomor, O., Schrempf, D., Woodhams, M.D., von Haeseler, A., Lanfear, R., 2020. IQ-TREE 2: New Models and Efficient Methods for Phylogenetic Inference in the Genomic Era. Molecular Biology and Evolution 37, 1530–1534. 10.1093/molbev/msaa015

35. Mölder, F., Jablonski, K.P., Letcher, B., Hall, M.B., Tomkins-Tinch, C.H., Sochat, V., Forster, J., Lee, S., Twardziok, S.O., Kanitz, A., Wilm, A., Holtgrewe, M., Rahmann, S., Nahnsen, S., Köster, J., 2021. Sustainable data analysis with Snakemake. F1000Res 10, 33. 10.12688/f1000research.29032.1

36. Muyle, A., Käfer, J., Zemp, N., Mousset, S., Picard, F., Marais, G.A., 2016. SEX-DETector: A Probabilistic Approach to Study Sex Chromosomes in Non-Model Organisms. Genome Biol Evol 8, 2530–2543. 10.1093/gbe/evw172

37. Ohno, S., 1967. Sex Chromosomes and Sex-linked Genes. Springer, New York.

38. Otto, S.P., Pannell, J.R., Peichel, C.L., Ashman, T.-L., Charlesworth, D., Chippindale, A.K., Delph, L.F., Guerrero, R.F., Scarpino, S.V., McAllister, B.F., 2011. About PAR: The distinct evolutionary dynamics of the pseudoautosomal region. Trends in Genetics 27, 358–367. 10.1016/j.tig.2011.05.001

39. Palmer, D.H., Rogers, T.F., Dean, R., Wright, A.E., 2019. How to identify sex chromosomes and their turnover. Molecular Ecology 28, 4709–4724. 10.1111/mec.15245

40. Paradis, E., Schliep, K., 2019. ape 5.0: an environment for modern phylogenetics and evolutionary analyses in R. Bioinformatics 35, 526–528. 10.1093/bioinformatics/bty633

41. Peona, V., Palacios-Gimenez, O.M., Blommaert, J., Liu, J., Haryoko, T., Jønsson, K.A., Irestedt, M., Zhou, Q., Jern, P., Suh, A., 2021. The avian W chromosome is a refugium for endogenous retroviruses with likely effects on female-biased mutational load and genetic incompatibilities. Philosophical Transactions of the Royal Society B: Biological Sciences 376, 20200186.

42. Purcell, S., Neale, B., Todd-Brown, K., Thomas, L., Ferreira, M.A.R., Bender, D., Maller, J., Sklar, P., de Bakker, P.I.W., Daly, M.J., Sham, P.C., 2007. PLINK: A Tool Set for Whole-Genome Association and Population-Based Linkage Analyses. The American Journal of Human Genetics 81, 559–575. 10.1086/519795

43. Revell, L.J., 2024. phytools 2.0: an updated R ecosystem for phylogenetic comparative methods (and other things). PeerJ 12, e16505. 10.7717/peerj.16505

44. Rice, W.R., 1987. The Accumulation of Sexually Antagonistic Genes as a Selective Agent Promoting the Evolution of Reduced Recombination between Primitive Sex Chromosomes. Evolution 41, 911–914.

45. Ruzicka, F., Dutoit, L., Czuppon, P., Jordan, C.Y., Li, X.-Y., Olito, C., Runemark, A., Svensson, E.I., Yazdi, H.P., Connallon, T., 2020. The search for sexually antagonistic genes: Practical insights from studies of local adaptation and statistical genomics. Evolution Letters 4, 398–415. 10.1002/evl3.192

46. Sigeman, H., Ponnikas, S., Chauhan, P., Dierickx, E., Brooke, M. de L., Hansson, B., 2019. Repeated sex chromosome evolution in vertebrates supported by expanded avian sex chromosomes. Proceedings of the Royal Society B: Biological Sciences 286, 20192051.

47. Sigeman, H., Sinclair, B., Hansson, B., 2022. Findzx: an automated pipeline for detecting and visualising sex chromosomes using whole-genome sequencing data. BMC Genomics 23, 328. 10.1186/s12864-022-08432-9

48. Smeds, L., Warmuth, V., Bolivar, P., Uebbing, S., Burri, R., Suh, A., Nater, A., Bureš, S., Garamszegi, L.Z., Hogner, S., Moreno, J., Qvarnström, A., Ružić, M., Sæther, S.-A., Sætre, G.-P., Török, J., Ellegren, H., 2015. Evolutionary analysis of the female-specific avian W chromosome. Nat Commun 6, 7330. 10.1038/ncomms8330

49. Tomaszkiewicz, M., Medvedev, P., Makova, K.D., 2017. Y and W Chromosome Assemblies: Approaches and Discoveries. Trends in Genetics 33, 266–282. 10.1016/j.tig.2017.01.008

50. Trivers, R., 1985. Social Evolution. Benjamin/Cummings, Menlo Park, California.

51. Valenzuela, N., Adams, D.C., Janzen, F.J., 2003. Pattern Does Not Equal Process: Exactly When Is Sex Environmentally Determined? The American Naturalist 161, 676–683. 10.1086/368292

52. Xirocostas, Z.A., Everingham, S.E., Moles, A.T., 2020. The sex with the reduced sex chromosome dies earlier: a comparison across the tree of life 16, 20190867. 10.1098/rsbl.2019.0867

53. Xu, L., Auer, G., Peona, V., Suh, A., Deng, Y., Feng, S., Zhang, G., Blom, M.P.K., Christidis, L., Prost, S., Irestedt, M., Zhou, Q., 2019. Dynamic evolutionary history and gene content of sex chromosomes across diverse songbirds. Nat Ecol Evol 3, 834–844. 10.1038/s41559-019-0850-1

54. Zhang, H., Lundberg, M., Tarka, M., Hasselquist, D., Hansson, B., 2023. Evidence of Site-Specific and Male-Biased Germline Mutation Rate in a Wild Songbird. Genome Biology and Evolution 15, evad180. 10.1093/gbe/evad180

55. Zhou, Q., Zhang, J., Bachtrog, D., An, N., Huang, Q., Jarvis, E.D., Gilbert, M.T.P., Zhang, G., 2014. Complex evolutionary trajectories of sex chromosomes across bird taxa. Science 346, 1246338. 10.1126/science.1246338

56. Zhu, Z., Younas, L., Zhou, Q., 2025. Evolution and regulation of animal sex chromosomes. Nat Rev Genet 26, 59–74. 10.1038/s41576-024-00757-3

