## Supplementary Text: Documentation, Material and Methods for "PhaseWY: A pipeline for haplotype phasing, sex chromosome identification and extraction of sex-limited sequences"

### 1 Detailed pipeline description

#### 1.1 Installation and usage

Instructions on how to install and run PhaseWY can be found on Github (https://github.com/sjellerstrand/Snakemake_PhaseWY) and below. All of the software and dependencies for running the pipeline can be installed using the conda environment and package manager (Anaconda Software Distribution, 2016). The pipeline has been tested using both conda version 24.11.3 and 25.11.3. It was developed and testing on a high-performance computing (HPC) platform that uses the SLURM queue system.

#### 1.2 Software's

The pipeline is run with snakemake v.8.27.1 (Mölder et al., 2021) using the snakemake executor plugin for slurm (version 0.15.0). The snakemake environment can be recreated using conda and the 'environment.yaml' file in the github repository. For reproducibility purposes and to allow for specific versions of packages to be used at different pipeline stages, an individual conda environment is created within each Snakemake rule. These environments are created the first time the pipeline is run and stored in the working directory. The software's and versions used within the pipeline are the following:

- BEDTools v.2.31.0 (Quinlan and Hall, 2010)
- BCFtools v.1.20 (Danecek et al., 2021)
- CrossMap v.0.7.3 (Zhao et al., 2014)
- MUMmer v.4.0.1 (Marçais et al., 2018)
- Perl v.5.32 (Wall et al., 2020)
- Plink v.1.90b6.21 (Purcell et al., 2007)
- Python v.3.13.2 (Python Software Foundation, 2025)
- R v.4.0 (R Core Team, 2021)
- R-packages:
    - data.table v.1.12.8 (https://Rdatatable.gitlab.io/data.table)
    - dplyr v.1.0.10 (Wickham et al., 2025)
    - ggplot2 v.3.3.6 (Wickham, 2016)
    - ggridges v.0.5.4 (Wilke, 2025)
    - gridExtra v.2.3 (Auguie and Antonov, 2017)
    - Tidyverse v.1.3.2 (Wickham et al., 2019)
    - vcfR v.1.10.0 (Knaus and Grünwald, 2017)
    - viridis v.0.6.2 (Garnier et al., 2024)
- SAMtools v.1.20 (Li et al., 2009)
- Seqtk v.1.4 (https://github.com/lh3/seqtk)
- SHAPEIT v.4.1.3 (Delaneau et al., 2019)
- vcflib v.1.0.3 (Garrison et al., 2022)
- VCFtools v.0.1.16 (Danecek et al., 2011)
- WhatsHap v.1.7 (Martin et al., 2016)

#### 1.3 Input files

##### Config-file

The pipeline is configured through a config-file, which contains paths to relevant input files required by the pipeline. It also contains filtering and run parameters that should be set by the user. First, a project name is required which will be included in the name of the top-folder in the results and intermediate folder hierarchies. This allows the user to run several projects within the same snakemake folder. Note that the top folder name will further contain other parameters that are set in the config-file, such as the sex depth threshold and the whatshap setting. Within each project folder, some steps will further be divided by haplotype cluster parameters to allow the user to explore several settings in parallel.

##### Reference assembly

The method relies on the alignment of Y/W sequences to their respective homologous X/Z reference. Therefore, the reference assembly (fasta) should be based on the homogametic sex. Alternatively, Y/W contigs should be excluded before read alignment.

##### Contig list

The user should provide a list of contigs to be analysed. For example, the user might already know which contig is sex-linked or want to exclude very short contigs, or the user might prefer to perform a genome-wide exploratory analysis. (See also "Masked regions"). The contig list is provided in a text file with one contig on every row:

CADDXX010000058.1
CADDXX010000069.1
CADDXX010000137.1

##### Read alignments

Bam-files are used to assess callable regions, sex alignment depth differences, and read-based phasing per individual. The bam-files need an @RG header with the SM-tag of the corresponding sample name. The pipeline will use all reads available in the bam-files. Therefore, it is up to the user to properly filter the bam files from duplicates and e.g. low quality or improper pairing (e.g. samtools -f2 -F260 -q20).

##### Variants

Variants are provided in a vcf-file, which contain all individuals to be used in the analysis, as well as the variants the user wants to classify. Biallelic and multiallelic SNPs, indels and MNPs are supported, as are monomorphic sites. Since the provided variants are used to infer phase and haplotype clustering, it is up to the user to properly filter variants before running the pipeline. In addition, many variants take longer to process. It is thus highly recommended that filters are applied before running the pipeline and done so with care. Since every user applies their own filtering scheme, the user should think carefully of how their filters might affect the outcome of the pipeline. For example, allelic balance filters could be unsuitable, since there are likely alignment biases between X/Z and Y/W sequence to the homologous X/Z reference. Likewise, excess heterozygosity is expected in the sex-linked region of the heterogametic sex, and too stringent filters on heterozygosity or HWE should be avoided. Furthermore, the heterogametic sex will experience half the coverage in old sex-linked regions. Therefore, too high minimum depth filters might filter out X/Z sequences, which could otherwise be extracted for analysis by the pipeline. Missing genotypes will be imputed during statistical phasing.

Imputation might be  less accurate in small datasets, or in datasets with high missingness. Therefore, it is up to the user to apply strict missingness filters. Finally, even if the user is mainly interested in specific genomic elements such as exons, flanking regions may still contain information utilised for accurate phasing and should not be filtered until after running the pipeline.

##### Sample information

Sample information is provided in a tab-delimited file with five columns. The first column contains the sample ID, which must correspond to the name in the vcf-file, as well as the SM-tag in respective bam-file. The second column contain individual mean depths, which are used to normalize depth when inferring sex differences in alignment depth. We suggest that these are estimated from variants in the input vcf-file, for example using vcftools:

```
vcftools --gzvcf  example/ Rasolark_variants.vcf.gz --depth --out Rasolark_variants
```

The third column contains the sex of each individual. To infer sex differences, each individual needs to be annotated as homogametic (HOMGAM) or heterogametic (HETGAM). The fourth column contains ploidy, which is not currently implemented. Note that the current version of the pipeline will only accept diploid individuals and is not designed to work with aneuploidies. The fifth column contains paths to individual bam-files is used for easy access in various parts of the pipeline. The samples information is provided with one individual on each row:

| sample_name | depth | sex | ploidy | bamfile |
|---|---|---|---|---|
| TT95871 | 34.8395 | HETGAM | 2 | example/TT95871.bam |
| TT95873 | 26.9165 | HOMGAM | 2 | example/TT95873.bam |

##### Callable filtering parameters

Callable regions across the genome are identified and retained if meeting a set criterion of minimum depth, minimum mean depth, maximum mean depth, and missingness. These parameter values are then applied in the config-file. We suggest that the user provides the same parameters used for variant filtering, since these filters will likewise be applied to variants provided in the vcf.

##### Masked regions (optional)

The user may want to mask some regions of the genome for various reasons. For example, repetitive regions might be unreliable and can be inferred with repeatmasker (https://www.repeatmasker.org/). These regions can be provided in a bed-file and will not be considered callable, will not be analysed for sex differences in alignment depth, and neither will any variants present in such regions be included in the output.

##### Syntenic reference genome (optional)

If the genome assembly of the study species is highly fragmented, the user might want to visualise results across the genome of a closely related species with a chromosome-level assembly. If a syntenic reference genome from a related species is provided (fasta), a genome alignment and lift-over of genome classification will be performed. Note that the quality of a cross-species alignment is highly dependent on genome divergence.

##### Example dataset

An example config-file, contig list and sample information is provided with the pipeline. A corresponding example dataset (3.5 Gb) of the Raso lark (*Alauda razae*) can be downloaded from https://zenodo.org/records/19050140, and includes a reference assembly, read alignments, variants, masked regions, and the Great tit genome (*Parus major*) to be used as a syntenic reference genome.

#### 1.4 Pipeline steps

Here follows a detailed description of the PhaseWY steps and the corresponding rules. Each step has its own snakemake file (smkfile) containing the corresponding rules. Note that various steps of the pipeline alter between parallel and separate operations per contig or individual. Furthermore, these steps don't necessarily represent the order of which they are executed.

##### Step 1: Identify callable regions

The first step of the pipeline is to determine which part of the genome to analyse, and which regions are callable (i.e. proper alignments to the reference genome), and which regions to mask.

**Rule 1.1**: The provided list of contigs (contig_list) is used to subset the reference genome (genome) with Seqtk. The genome index file (.fai) can be provided to analyse the full genome.

**Rule 1.2**: The subsetted genome is divided into large and small contigs based on the threshold set in the contig file (clump_max_len). Large contigs will in some cases be run as independent jobs, while small contigs will be analyses through a loop in a single job. Following this step, a checkpoint will dynamically update the job schedule to the total number of jobs to be run.

**Rule 1.3**: The depth in each individual and at every site of the contig is retrieved with SAMtools depth and stored in a table.

**Rule 1.4**: Each site in the depth table is evaluated and retained if a set of criteria based on filtering parameters provided in the config file are met. At each site, individuals with a depth lower than the minimum depth (min_dp) are set to missing. Then the site is evaluated for the proportion of missingness (missing), and the threshold for minimum mean (min_mean) and maximum mean (max_mean) depth.

**Rule 1.5**: The sites meeting the filtering criteria are merged into coherent regions with BEDTools , creating a summary of callable regions. If a mask file is provided, masked regions are subtracted from the callable regions.

##### Step 2: Identify sex-linked regions based on differences in sequence depth

The second step of the pipeline is to identify sites in the genome which differ in alignment depth between heterogametic and homogametic individuals, suggesting sex-linkage due to sequence degeneration/divergence of the Y/W chromosome.

**Rule 2.1**: The depth table produced in rule 1.3 is used to normalize individual depths through division by the individual mean depths provided in the sample table file (sample_table). These normalized individual averages are then further averaged across heterogametic and homogametic individuals separately. Finally, a depth score is calculated for each site by dividing heterogametic mean depth with homogametic mean depth.

**Rule 2.2**: If the calculated depth score is lower than the threshold set in the config file (sex_depth_threshold), the site is kept as an indication of Y/W degeneration. All sites are merged into coherent regions with BEDTools and is likewise intersected with the callable regions identified in rule 1.5.

**Rule 2.3**: The sex depth scores from callable regions are averaged in sliding windows. These scores will be used in the haplotype clustering in rule 6.2 and then plotted in rule 10.5. Window size (window) and step size (step) can be set in the config file and correspond to the parameters for the haplotype clustering in rule 6.2.

**Rule 2.4**: The sex depth scores are subsampled from callable regions to produce distribution plots in rule 10.9. The percentage of sites to subsample (actually the probability of sampling a site when streaming through the depth score file) is set in the config file (subsample).

##### Step 3: Prepare variants for phasing

Variants in the input vcf (input_vcf) are evaluated and filtered. At least two heterozygous genotypes are needed in a contig for compatibility with phasing and downstream analyses. Further, multiallelic sites are split into biallelic records to allow them to be phased with downstream phasing software.

**Rule 3.1**: The input vcf is evaluated to identify genotype frequencies across all contigs in the provided contig list (contig_list). Sites are excluded based on parameters specified in the config file using VCFtools. These are minimum genotype depth (min_dp), the proportion of missingness (missing), and the threshold for minimum mean (min_mean) and maximum mean (max_mean) depth. If a mask file is provided, variants in mask regions are likewise excluded.

**Rule 3.2**: Contigs with less than two heterozygous genotypes are excluded from all analyses based on phasing and phased haplotypes (step 4-7). If a contig does not have enough heterozygote sites to be phased, the sites are placed in a separate file in rule 9.2 (nonphased_variants.vcf.gz).

**Rule 3.3**: The input vcf (input_vcf) is subsetted to the contigs provided in the contig list (contig_list) and have enough heterozygous genotypes to be phased. Sites are excluded based on parameters specified in the config file using VCFtools. These are minimum genotype depth (min_dp), the proportion of missingness (missing), and the threshold for minimum mean (min_mean) and maximum mean (max_mean) depth. If a mask file is provided, variants in mask regions are likewise excluded. To save disk space, the subsetted vcf is sripped from FORMAT and INFO fields. Finally, all multiallelic sites are split into biallelic sites using BCFtools norm.

##### Step 4: Read-based phasing

Read-based phasing is performed on each individual by inspecting the bam file for read-pairs spanning as many heterozygote genotypes as can be observed in physical linkage on the same contig. These are then recorded as phase-sets. Singleton alleles that exist in both a heterogametic individual and in a sex-linked region must be included in a phase informative phase-set to be placed on either the X/Z or Y/W chromosome. If they do not have phase information, they are recorded. Rule 4.1 to 4.6 will run if Whatshap is toggled by the parameter in the config file (whatshap). Only rule 4.7 will run if Whatshap is disabled, which will record every singleton that exist in both a heterogamete individual and in a sex-linked region.

**Rule 4.1**: The filtered vcf from rule 3.3 is subset into per-individual vcf files using BCFtools.

**Rule 4.2**: Read-based phasing is performed on each individual using Whatshap. It does this by inspecting the individual bam-file for reads which link alleles in at least two heterozygous genotypes physically. Alleles linked in this way are recorded as phase-sets, with each phase-set spanning as many heterozygous sites that can be physically linked. Only heterozygous variants in the individual vcf file from rule 4.1 are evaluated.

**Rule 4.3**: All individual vcf files from rule 4.2 are merged into a single vcf file using BCFtools.

**Rule 4.4**: All singleton alleles in heterogametic individuals and within any potentially sex-linked regions (as determined in rule 8.4) are extracted using BCFtools and vcflib.

**Rule 4.5**: Each singleton, its corresponding individual, as well as phase-set information is extracted using BCFtools and vcflib.

**Rule 4.6**: Singletons without phase-set information are extracted. First, singletons not included in phase-sets are extracted. Second, singletons included in a phase-set are evaluated whether that phase-set is informative using BCFtools and vcflib. Variants within the same individual and within 2 500 bases at each site of the variant and within the same phase-set are extracted. If none of these sites are heterozygous in any other individual, the placement of the singleton on X/Z or Y/W cannot be inferred and is likewise extracted. None of these singletons are excluded from the vcf-file at this point but are placed in a separate file in rule 9.7 (heterogametic_sexlinked_singletons.vcf.gz).

**Rule 4.7**: All singleton alleles in heterogametic individuals and within any potentially sex-linked regions (as determined in rule 8.4) are extracted using BCFtools and vcflib. None of these singletons are excluded from the vcf-file at this point but are placed in a separate file in rule 9.7 (heterogametic_sexlinked_singletons.vcf.gz).

##### Step 5: Statistical phasing

Statistical phasing is performed to produce a best solution phase across each contig. If Whatshap was used to produce phase-sets, these will be applied to inform the phasing inference. Further, statistical phasing will impute missing genotypes based on haplotype information. Note that singletons without any phase-set information will be placed randomly on either haplotype.

**Rule 5.1**: Statistical phasing is performed with SHAPEIT4. If Whatshap was used to produce phase-sets, the vcf file from rule 4.4 is used. If Whatshap was not used, the vcf file from rule 3.3 is used. This step runs per contig and produces one phased vcf per contig.

**Rule 5.2**: All multiallelic variants are reconstructed from biallelic records into multiallelic records with BCFtools norm.

##### Step 6. Haplotype clustering based on sex-linkage

A sliding window analysis is performed on each contig to infer sex-linkage through haplotype clustering. First, low frequency alleles are excluded. Then haplotypes in each window is forced into two clusters with k-means. If these clusters pass a set of criteria, they are classified as sex-linked, and the respective Y/W haplotypes are recorded. Several clustering statistics are outputted with respective window, as is the regions classified as sex-linked. Finally, some variants that indicate poor phasing or incomplete lineage sorting are recorded.

**Rule 6.1**: The phased vcf file is filtered to exclude multiallelic sites and low frequency alleles using BCFtools, VCFtools and vcflib. The allele frequencies to exclude can be set by the minor allele count parameter in the config file (mac). Note that these are not excluded from the final dataset, only from the haplotype clustering inference.

**Rule 6.2**: The phased and mac-fitlered vcf file is analysed per contig with a custom R-script. The vcf file is uploaded with vcfR and each individual haplotype along the contig is split into the "left" and "right" side of the genotype pipe ("|"). The sex depth score averages in windows from rule 2.3 is imported, which contain the windows to analyse. The contig is then evaluated in a sliding window analysis. Window size (window) and step size (step) can be set in the config file.

Variants within the window under analysis are extracted from the haplotypes using data.table. If there are no variants within the window, the window is reported as autosomal. If there are variants present, the haplotypes are forced into two clusters with k-means (Hartigan and Wong, 1979), applying 10 random sets. Based on the number of males and females in the dataset, there is a predefined expectation on the number of haplotypes in the X/Z-haplotype cluster ($2*N_{homogametes} + 1*N_{heterogametes}$) and the Y/W-haplotype cluster ($1*N_{heterogametes}$) respectively. The smallest cluster (containing the putative Y/W-haplotypes) is classified as autosomal if failing any of three criteria that are evaluated in the following order: i) only haplotypes from heterogametic individuals are present in the smallest cluster, ii) only one haplotype per heterogametic individual is present in the smallest cluster, and iii) the smallest cluster correspond to the expected number of Y/W haplotypes. If the window is classified as sex-linked, the haplotypes within the smallest cluster is annotated as Y/W. The default model for clustering is using absolute distances (or Hamming distances). Alternatively, the importance of a variant can be weighted by the inverse of the minor allele frequency (Inverse MAF), making rare variants more important since they are likely to share recent identity by decent. The preferred model can be set in the contig file (dist_model).

Data from each window is recorded and include; if the window is sex-linked or autosomal (and if autosomal, which criterion the window failed), the number of haplotypes present in the smallest and largest cluster, the total sums of squares of the data, the sums of squares between the clusters, and the sums of squares of the smallest and largest cluster, the number of variants, sex differences in depth, sex differences in heterozygosity, and the Y/W haplotype of each heterogametic individual (if sex-linked, otherwise set as "Unknown"). Likewise, the window is annotated as phase informative in the column "Phase information available". These window-based statistics will be summarised across the genome in rule 8.4, and several of the statistics plotted in rule 10.5.

When an autosomal window transitions into a sex-linked window or vice versa, there will be some uncertainty in which variants are autosomal or sex-linked and were the exact border should be placed. When this occurs, the putatively autosomal window is evaluated for sex depth difference to verify that the region is not X/Z with absence of a Y/W sequence. If the region is classified as an autosomal transition, all overlapping windows are annotated in the column "Border change". If not, the window is instead set to autosomal and annotated as phase informative in the column "Phase information available". If there is a change in the Y/W haplotype for any heterogametic individual, all overlapping windows are annotated in the column "Phase switch" to indicated that these variants contain some uncertainty. Likewise, the haplotype of the corresponding individual is set to "Unknown".

Finally, three types of files are printed. One file contains the raw information from the sliding window analysis (<CONTIG>_phase_windows.bed). One file is output for every heterogametic haplotype containing the putative Y/W-variants (<IND>_het_left.bed and <IND>_het_right.bed). The third file contains a simplified summary of the analysis, containing coherent regions classified as "Autosomal", "Sex-linked", or "Unknown" (<CONTIG>_phase_info.bed). "Unknown" regions are in this file based on autosomal transitions and not phase switches.

**Rule 6.3**: Regions identified as sex-linked in rule 6.2 are extracted and merged into coherent regions using BEDTools .

**Rule 6.4**: Biallelic sites within sex-linked regions that are homozygous for both alleles in heterogametic individuals are extracted using BCFtools and VCFtools.

#### Step 7. Re-organise genotypes according to genomic region

All variants are reorganised and re-genotyped based on the classification of sex-linkage in previous steps. The genotypes of homogametic individuals are reported as diploid for the shared sex chromosome. The haplotypes of heterogametic individuals are split into haploid genotypes, reorganised, and output separately for the shared and sex-limited chromosomes. The sex-linekd variants in heterogametes are treated differently depending on whether they have been detected through a difference in alignment depth, or haplotype clustering. Importantly, sex-limited genotypes at sites that have also been detected through sex differences in depth are deemed unreliable and consequently excluded from the sex-limited file. Sites with sex depth differences in the shared sex chromosome are recoded. Homozygous genotypes are re-genotyped to haploid, and heterozygous genotypes are set to missing. Finally, some variants that indicate poor phasing or incomplete lineage sorting are recorded.

**Rule 7.1**: Regions classified with sex-depth differences in rule 2.2 and sex-linked through haplotype clustering in rule 6.2 are merged into coherent regions with BEDTools. Phased variants from rule 5.2 that are located in sex-linked regions are then extracted and subset into two vcf-files, one for heterogametic and for homogametic individuals respectively. All sex-linked variants are extracted from homogametic individuals, while sites with sex-depth differences are not included in the output for heterogametic individuals.

**Rule 7.2**: Sex-linked variants in heterogametic individuals are split into two haploid vcf-files, each containing the haplotype on the left side of the phased genotype, as well as the one on the right side. For each individual, the haplotype corresponding to the X/Z and the one corresponding to the Y/W as classified in rule 6.2 are subset into separate vcf-files. These are subsequently concatenated across contigs into two haploid vcf-files, each corresponding to the X/Z and Y/W genotypes. Each data set is merged into a single file across heterogametic individuals. Finally, any site classified with sex-depth differences in rule 2.2 is filtered from the vcf-file corresponding the sex-limited Y/W haplotype.

**Rule 7.3**: The haploid X/Z haplotype subset from heterogametic individuals in rule 7.2 is merged. Then, phased variants from rule 5.2 that are located in regions classified with sex-depth differences in rule 2.2 are extracted from heterogametic individuals. These are representing putatively haploid X/Z sequences, why homozygote genotypes are re-coded as haploid, while heterozygous genotypes are coded as missing. All haploid X/Z variants in heterogametic individuals are concatenated, then merged with the diploid genotypes of homogametic individuals subset in rule 7.1.

**Rule 7.4**: Sites that occurring in both the X/Z and the Y/W vcf-files are extracted with BEDTools. Sites that are polymorphic on both the X/Z and the Y/W are extracted.

**Rule 7.5**: This rule more or less performs the same procedure as rules 7.1-7.4. However, it specifically performs on contigs were regions have been classified with sex-depth differences in rule 2.2, but appear completely autosomal based on classification through haplotype clustering in rule 6.2.

**Rule 7.6**: Autosomal regions are extracted by excluding regions classified with sex-depth differences in rule 2.2 and sex-linked through haplotype clustering in rule 6.2. Phased variants from rule 5.2 that are located in autosomal regions are then extracted.

##### Step 8. Summarise genome with bed-files

The result of the pipeline is summarised in genome-wide bed files using BEDTools . These correspond to regions which are callable or missing, which have been phased, which are classified as autosomal or X/Z and Y/W, and individual sites which are considered unreliable for various reasons. It also summaries statistics that are output from the haplotype clustering algorithm in rule 6.2.

**Rule 8.1**: Callable regions from rule 1.5 are combined into a genome-wide bed file.

**Rule 8.2**: Regions with a sex depth difference from rule 2.2 are combined into a genome-wide bed file.

**Rule 8.3**: The callable regions from rule 8.1 are filtered from non-phased contigs identified in rule 3.2. Non-phased contigs are output in a separate file.

**Rule 8.4**: Output from the haplotype clustering analysis in rule 6.2 is retained:

- The per window data is combined into a genome-wide bed file. All sex-linked regions are output in a separate bed file.
- The classification of coherent regions is likewise combined into a genome-wide bed file containing regions that are “Autosomal”, “Sex-linked” and “Unknown”. All sex-linked regions are output in a separate bed file and differ from the previous sex-linked regions file in that it does not contain “Unknown” regions, i.e. autosomal transitions in which classification is uncertain. It is also intersected with phased callable regions from rule 8.3.
- A bed file is produced containing all sex-linked regions identified through phasing, excluding autosomal transitions and regions that have been sex-linked through windowed sex depth differences .

**Rule 8.5**: All “Unknown” regions are extracted from the coherent regions file in rule 8.4. These regions correspond to autosomal transitions identified in rule 6.2. The file is intersected with phased and callable regions from rule 8.3. Further, regions with sex depth difference are subtracted from these unknown regions since they are considered confident.

**Rule 8.6**: Unknown regions from rule 8.5 are subtracted from phased and callable regions from rule 8.3 to create the final callable regions file with confidence regions.

**Rule 8.7**: A bed file representing genome-wide missing regions is created as the complement of the final callable regions from rule 8.6.

**Rule 8.8**: The final bed files corresponding to genomic regions are made:

- Regions with a sex depth difference from rule 8.2 are intersected with the final callable regions from rule 8.6 to exclude non-phased contigs.
- Regions with a sex depth difference from rule 8.2 are merged with phased and filtered sex-linked regions from rule 8.4 to create a file containing all sex-linked regions identified through both sex depth differences and haplotype clustering. These regions also correspond to regions where X/Z data can be retained.
- The regions with sex depth difference are subtracted from the total sex-linked regions to represent sex-linked regions with even depth between the sexes. These regions also correspond to regions where Y/W data can be retained.
- Sex linked regions are subtracted from the final target regions from rule 8.6. These correspond to regions classified as autosomal.

**Rule 8.9**: Variants which may be problematic due to unsuccessful phasing or incomplete lineages sorting are concatenated:

- All variants identified as problematic in rule 6.4 are concatenated.
- Regions with sex depth difference from rule 8.2 are subtracted from the variants identified in rule 6.4, since they may also correspond to absence of Y/W sequences.
- All variants identified as problematic in rule 7.4 are concatenated.

**Rule 8.10**: If Whatshap is enabled, the position of all non-phased singleton from rule 4.6 are concatenated into a bed file. If Whatshap was disabled, all singletons are instead output in the corresponding file in rule 4.7.

**Rule 8.11**: All sites in the genome considered unreliable are concatenated into a single bed file. These correspond to autosomal transition regions from rule 8.5, non-phased contigs from rule 8.3, problematic variants from rule 6.4/8.9 with no sex depth difference, problematic variants from rule 7.4/8.9, and non-phased singletons from rule 8.10.

#### Step 9. Summarise genome with vcf-files

The classified variants are further subset in genome-wide vcf files using BCFtools. These correspond to variants in regions which have been phased, which are classified as autosomal or X/Z and Y/W, and individual variants which are considered unreliable for various reasons. All heterogametic individuals are represented as haploid in vcf files corresponding to X/Z and Y/W variants. Further, autosomal, X/Z, and Y/W variants are outputted unfiltered, involving raw classification from the haplotype clustering in step 6.2. Therefore, these files contain variants with inconsistent classification and variants classified as potentially problematic. A second set of files is therefore also produced, with overlapping and problematic sites excluded.

**Rule 9.1**: The phased variants from rule 5.2 are concatenated.

**Rule 9.2**: First, all variants organised as autosomal in rule 7.6 are concatenated. Second, these variants are further intersected with the autosomal regions from rule 8.8, with unreliable variants from rule 8.11 excluded to create a filtered autosomal vcf.

**Rule 9.3**: First, all variants organised as Y/W in rule 7.2 and X/Z in rule 7.3 are concatenated in separate vcf files. Second, these variants are further intersected with the corresponding regions from rule 8.8, with unreliable variants from rule 8.11 excluded to create a filtered X/Z and Y/W vcf.

**Rule 9.4**: Any variants present in non-phased contigs identified in rule 3.2/8.3 are extracted from the input vcf. The vcf is filtered with VCFtools to match filtering parameters applied in rule 3.3.

**Rule 9.5**: Variants present in regions with autosomal transitions from rule 6.2/8.5 are extracted from the input vcf. The vcf is filtered with VCFtools to match filtering parameters applied in rule 3.3.

**Rule 9.6**: Variants which may be problematic due to unsuccessful phasing or incomplete lineages sorting are extracted. Each set of problematic variants are extracted for all individuals from the input vcf, for all individuals from the X/Z-linked region in rule 8.8/9.3, and for all heterogametic individuals from Y/W-linked regions in rule 8.8/9.3.

- All variants identified as problematic in rule 6.4/8.9 are extracted.
- variants identified as problematic in rule 6.4/8.9, but excluding regions with sex depth difference from rule 8.2 are extracted.
- All variants identified as problematic in rule 7.4/8.9 are extracted.

**Rule 9.7**: All non-phased singletons from rule 4.6/4.7/8.10 are extracted from the input vcf.

**Rule 9.8**: All variants in the genome considered unreliable are concatenated into a single vcf file. These correspond to autosomal transition regions from rule 8.5/9.5, non-phased contigs from rule 8.3/9.4, problematic variants from rule 6.4/8.9/9.6 with no sex depth difference, problematic sites from rule 7.4/8.9/9.6, and non-phased singletons from rule 4.6/4.7/8.10/9.7. The two sets of problematic variants from rule 6.4/7.4/8.9/9.6 originate from the input vcf.

##### Step 10. Pipeline accuracy summary

A summary of the accuracy of the pipeline is visualised in various ways. Some statistics and a PCA is produced for each vcf file from step 9. Some of the per window statistics from the haplotype clustering algorithm in rule 6.2/8.4 is visualised as Manhattan plots. The classification of the genome as missing, autosomal, and methods of classifications as sex-linked are visualised along the genome. These classifications are also visualised as sex depth difference distributions. Finally, a synteny alignment to a related species can be performed which liftover the genome classification and visualises it along the syntenic reference genome.

**Rule 10.1**: For every vcf-file produced in step 9 with at least one variant, VCFtools is used to produce per variant and per individual data. These include missingness per site, missingness per individual, minor allele frequency (folded Site Frequency Spectrum), per individual Inbreeding Coefficient ($F_{IS}$), and the total number of variants.

**Rule 10.2**: The statistics from rule 10.1 are visualised through density plots for each vcf. However, these plots are only produced for vcfs with at least one variant present.

**Rule 10.3**: For every vcf-file produced in step 9 with at least two variants, Plink is used to linkage prune the data using 50 kb windows, a step sizes of 10 kb, and an r-squared value of 0.

If at least one vcf is retained, Plink is used to calculate principal components of the remaining independent variants.

**Rule 10.4**: For every vcf, the first and second principal component calculated in rule 10.3 is visualised through scatterplots, with each individual coloured by sex. However, these plots are only produced for vcfs with at least two eigenvectors present.

**Rule 10.5**: Some of the per window statistics from the haplotype clustering algorithm in rule 6.2/8.4 is visualised through two sets of Manhattan plots:

- The sum of squares from k-means are visualised in four panels, including the total sums of squares, between cluster sums of squares, and within cluster sums of squares for the larges and the smallest cluster.
- Other statistics are visualised in four panels and include the number of haplotypes in the smallest cluster, the average sex depth difference, the average sex heterozygosity difference, and variant density.

In all plots, points are coloured by the classification of the window as sex-linked or autosomal (and if autosomal, which criterion the window failed). Some size parameters for the output plots can be set in the config file and include the width of the plot (width) and the height of the plot (height; since several plots are stacked on top of each other and include vertical contig names at the base, each figure height is multiplied by six inside R). In addition, small contigs can be excluded from the output plots by setting a minimum contig length threshold (min_len).

**Rule 10.6**: The classification of each region of the genome is subset into five bed files, then concatenated into one single file, representing a summary of the genome. These include "Missing data" from rule 8.7, "Autosomal" from rule 8.8, "Sex depth difference" from rule 8.2, and "Sex haplotype clustering" from rule 8.4. However, a fifth file is created as the intersect between sex depth differences and sex-linked regions based on haplotype clustering, and thus represent regions identified by both methods, i.e. "Sex haplotype clustering & depth difference". These overlapping regions are consequently subtracted from the sex depth difference regions and sex haplotype clustering regions. Finally, all five categories are concatenated into a single file.

**Rule 10.7**: The genome summary from rule 10.6 is visualised along the genome, each classification visualised by a specific colour. Some size parameters for the output plot can be set in the config file and include the width of the plot (width) and the height of the plot (height; since contig names are placed vertically at the base of the figure height is multiplied by two inside R). In addition, small contigs can be excluded from the output plot by setting a minimum contig length threshold (min_len).

**Rule 10.8**: The subsampled sex depth difference scores from rule 2.4 are concatenated into a single file. Then each genome category from rule 10.6 (except for missing) is intersected with these sex depth scores using BEDTools.

**Rule 10.9**: The sex depth difference score subset by category in rule 10.8 is visualised as histograms. Two plots are produced:

- One plot is a stacked histogram with each category stacked on top of each other in different colours. The proportion of each category corresponds to relative proportions in the dataset.

- One plot contains separate distributions for each category in the same colours as the previous plots. It also contains one distribution representing all sites together. These plots do not represent relative proportions to the other categories.

**Rule 10.10**: If the path to a syntenic reference genome has been set in the config file (synteny_genome), this genome will be aligned to the reference genome using nucmer from MUMmer. Then a chain file is produced with the perl script crossmap_delta_to_chain.pl from CrossMap.

**Rule 10.11**: The genome summary from rule 10.6 is lifted over to corresponding coordinates in the syntenic reference genome with CrossMap, using the chain file from the synteny alignment in rule 10.10. BEDTools is used to create a sixth category, “No alignment”, which corresponds to regions which could no align with the syntenic reference genome. The other five categories are extracted from the lift over, then merged into coherent regions with BEDTools. Finally, all six categories are concatenated into a single file.

**Rule 10.12**: The lifted over genome summary from rule 10.11 is visualised along the genome, each classification visualised by a specific colour. Some size parameters for the output plot can be set in the config file and include the width of the plot (width) and the height of the plot (height; since contig names are placed vertically at the base of the figure height is multiplied by two inside R). In addition, small contigs can be excluded from the output plot by setting a minimum contig length threshold (min_len).

#### 1.5 Output files

The pipeline outputs classified and reorganised variants, and reports information from the analysis in various ways. Thes include genomic regions in bed-files, and genotypes in vcf-files. Further, statistics are calculated for variants in respective vcf-file and visualised. Likewise, a PCA is performed on respective vcf-file and visualised. Finally, several genome classification summary statistics are output and visualised.

##### 1.5.1 bed-files

- **callable_regions.bed** – Callable regions based on depth and missingness of bam-files (rule 1.5/8.1).
- **callable_phased.bed** – Callable regions in contigs that could be phased (rule 8.3).
- **target_region.bed** – Callable regions in contigs that could be phased and with problematic regions with autosomal/sex-linked transitions excluded (rule 8.6).
- **missing_region.bed** – The inverse of target_region.bed, i.e. regions without proper alignments (uncallable), non-phased contigs, and problematic regions with autosomal/sex-linked transitions (rule 8.7).
- **hetgam_dropout.bed** – Callable regions with sex-depth differences (rule 8.2).
- **target_region_hetgam_dropout.bed** – Regions with sex-depth differences, intersected with target_region.bed, i.e. excluding non-phased contigs, and problematic regions with autosomal/sex-linked transitions (rule 8.8).
- **phased_windows.bed** – Per window statistics from the haplotype clustering algorithm in rule 6.2 (rule 8.4).
- **phased_info.bed** – Summary of the classification through the haplotype clustering algorithm in rule 6.2 in coherent genomic regions. Problematic regions with autosomal/sex-linked transitions are set as unknown (rule 8.4).

- **phased_sex_linked.bed** – All region identified as sex-linked through the haplotype clustering algorithm in rule 6.2, including problematic regions with autosomal/sex-linked transitions (rule 8.4).
- **callable_sex_linked.bed** – Callable regions that overlap with sex-linked regions identified through the haplotype clustering algorithm in rule 6.2. Problematic regions with autosomal/sex-linked transitions are excluded (rule 8.4).
- **autosomal.bed** – Regions classified as autosomal, excluding non-phased contigs and problematic regions with autosomal/sex-linked transitions (rule 8.8).
- **sex_linked.bed** – Regions classified as sex-linked, excluding non-phased contigs and problematic regions with autosomal/sex-linked transitions (note that this file corresponds to sexshared.bed; rule 8.8).
- **sexshared.bed** – Regions corresponding to the shared X/Z chromosome, excluding non-phased contigs and problematic regions with autosomal/sex-linked transitions (note that this file corresponds to sex_linked.bed; rule 8.8).
- **sexlimited.bed** – Regions corresponding to the sex-limited Y/W chromosome, excluding non-phased contigs and problematic regions with autosomal/sex-linked transitions (i.e. sex-linked sites with sex-depth differences excluded; rule 8.8).
- **nonphased_contigs.bed** – Contigs that had too few variants to be phased (rule 3.2/8.3).
- **border_variants.bed** – Problematic regions with autosomal to sex-linked transitions (rule 8.5).
- **sex_linked_ILS1.bed** – Sites classified as ILS1 (rule 8.9).
- **sex_linked_ILS1_not_hetgam_dropout.bed** – Sites classified as ILS1 but not present in regions with sex-depth differences (rule 8.9).
- **sex_linked_ILS2.bed** – Sites classified as ILS2 (rule 8.9).
- **nonphased_singletons_htgm_singletons.bed** – Sites with singletons present in sex-linked regions within heterogametic individuals, and which could not be phased by inclusion in a phase-set (rule 8.10).
- **all_unreliable_sites.bed** – Sites present in the files nonphased_contigs.bed, border_variants.bed, sex_linked_ILS1_not_hetgam_dropout.bed, sex_linked_ILS2.bed, and sex_linked_ILS1_not_hetgam_dropout.bed (rule 8.11).

##### 1.5.2 vcf-files

Note that files with the suffix "_filtered" have been filtered from variants present within the file all_unreliable_sites.vcf.gz. Also note that many variants may appear monomorphic once subset into a sex-shared and sex-limited sex chromosome file and may be filtered by the user if necessary.

- **phased_all_variants.vcf.gz** – All variants that could be phased variants (rule 9.1). Note that singletons without any phase-set information will be placed randomly on either haplotype.
- **autosomal.vcf.gz** – Unfiltered autosomal variants (rule 9.2).
- **autosomal_filtered.vcf.gz** – Filtered autosomal variants (rule 9.2).
- **sexshared.vcf.gz** – Unfiltered variants from the shared X/Z chromosome (rule 9.3).
- **sexshared_filtered.vcf.gz** – Filtered variants from the shared X/Z chromosome (rule 9.3).
- **sexlimited.vcf.gz** – Unfiltered variants from the sex-limited Y/W chromosome (rule 9.3).

- **sexlimited_fitlered.vcf.gz** – Filtered variants from the sex-limited Y/W chromosome (rule 9.3).
- **nonphased_variants.vcf.gz** – Variants on contigs that had too few variants to be phased (rule 9.4).
- **border_variants.vcf.gz** – Variants present in regions with autosomal to sex-linked transitions (rule 9.5).
- **all_inds_ILS1_not_hetgam_dropout.vcf.gz** – Sites classified as ILS1 with the genotypes from the input vcf-file.
- **sexshared_ILS1_not_hetgam_dropout.vcf.gz** – Sites classified as ILS1 with the genotypes subset to the shared sex-chromosome.
- **sexlimited_ILS1_not_hetgam_dropout.vcf.gz** – Sites classified as ILS1 with the genotypes subset to the sex-limited chromosome.
- **all_inds_ILS2.vcf.gz** – Sites classified as ILS2 with the genotypes from the input vcf-file.
- **sexshared_ILS2.vcf.gz** – Sites classified as ILS2 with the genotypes subset to the shared sex-chromosome.
- **sexlimited_ILS2.vcf.gz** – Sites classified as ILS2 with the genotypes subset to the sex-limited chromosome.
- **heterogametic_sexlinked_singletons.vcf.gz** – Singletons present in sex-linked regions within heterogametic individuals, and which could not be phased by inclusion in a phase-set (rule 9.7).
- **all_unreliable_sites.vcf.gz** – Variants present in the files nonphased_variants.vcf.gz, border_variants.vcf.gz, all_inds_ILS1_not_hetgam_dropout.vcf.gz, all_inds_ILS2.vcf.gz, and heterogametic_sexlinked_singletons.vcf.gz (rule 9.8). These are used to produce the files autosomal_filtered.vcf.gz, sexshared_filtered.vcf.gz, and sexlimited_fitlered.vcf.gz.

##### 1.5.3 vcf-statistics

For every output vcf-file with at least one variant, per variant and per individual statistics are output (rule 10.1-2):

- **<vcf-name>.lmiss** – Missingness per site.
- **<vcf-name>.imiss** – Missingness per individual.
- **<vcf-name>.frq** – Minor allele frequency (folded Site Frequency Spectrum).
- **<vcf-name>.het** – Per individual Inbreeding Coefficient ($F_{IS}$).
- **<vcf-name>_number_variants.txt** – The total number of variants.
- **<vcf-name>_stats.txt** – The minimum, median, mean, maximum and quantiles of each statistic.
- **<vcf-name>_stats.png** – The distribution of each statistic is visualised.

##### 1.5.4 PCA

For every output vcf-file with at least two variants, a PCA was performed (rule 10.3-4):

- **<vcf-name>_PCA.png** – The outcome of the PCA is visualised in a scatterplot along the first two PCs, with each sample coloured by sex.

##### 1.5.5 Genome alignment

- **<synteny_name>.delta** – The synteny alignment between the reference and query sequences in delta-format, which is output by the nucmer command from MUMmer.

- **<synteny_name>.filtered.delta** – A filtered synteny alignment in delta-format, which only includes 1-to-1 alignments.
- **fwd_<synteny_name>.chain** – Forward alignments in chain-format. The synteny alignment in delta-format from MUMmer converted by crossmap_delta_to_chain.pl from CrossMap
- **rev_<synteny_name>.chain** – Reverse alignments in chain-format. The synteny alignment in delta-format from MUMmer converted by crossmap_delta_to_chain.pl from CrossMap

##### 1.5.6 Genome summary

- **Cluster_info.png** – Per window statistics from the haplotype clustering algorithm in rule 6.2 is visualised in Manhattan plots in rule 10.5. The statistics are visualised in four panels and include the number of haplotypes in the smallest cluster, the average sex depth difference, the average sex heterozygosity difference, and variant density. In all panels, points are coloured by the classification of the window as sex-linked or autosomal (and if autosomal, which criterion the window failed).
- **Cluster_sums_of_squares.png** – Per window statistics from the haplotype clustering algorithm in rule 6.2 is visualised in Manhattan plots in rule 10.5. The sums of squares from k-means are visualised in four panels, including the total sums of squares, between cluster sums of squares, and within cluster sums of squares for the larges and the smallest cluster. In all panels, points are coloured by the classification of the window as sex-linked or autosomal (and if autosomal, which criterion the window failed).
- **Genome_summary bed** – The classification of every region of the genome from rule 10.6.
- **Genome_summary png** – The classification of every region of the genome is visualised along the genome in rule 10.7, each classification with different colour.
- **sexdiff_genome_summary_subset_<%>_percent_separate.png** – A set of histograms of sex depth difference in regions with various classifications produced in rule 10.9. Each classification is plotted separately, including one distribution representing all sites together. These plots do not represent relative proportions to the other categories.
- **sexdiff_genome_summary_subset_<%>_percent_stacked.png** – A stacked histogram of sex depth difference in regions with various classifications produced in rule 10.9. Each classification is stacked on top of each other in different colours. The proportion of each category corresponds to relative proportions in the dataset.
- **<synteny_name>_liftover_Genome_summary.bed** – The genome summary produced in rule 10.6 is lifted over to the syntenic reference genome (rule 10.11).
- **<synteny_name>_liftover_genome_summary.png** – The classification of every region of the genome is visualised along the syntenic reference genome in rule 10.12, each classification with different colour.

#### 1.6 Simple phasing with companion script

In addition to PhaseWY, we supply phasing_female_male_pair.py, a python companion script that extracts Y/W sequences from a single female-male pair based on sex-specific allele patterns. The script takes a vcf file with two individuals as input. The first individual should be the heterogamete, and the second the homogamete. The output then contains the homogamete as is, but with the heterogamete split into two haploid sequences representing the putative X/Z and Y/W. Note that INFO-fields for the heterogametic individual are removed, since they are genotype specific and not applicable once the genotype is split. Also note that the naming convention for the heterogametic haplotypes follows the Z/W system (this can be modified in the last few lines of the script).

At each site, the heterogametic and homogametic alleles are evaluated. If the heterogamete is homozygote, the site is considered to be non-variable between sex chromosomes and therefore retained. If the heterogamete is heterozygous, the algorithm checks whether one allele is shared between the female and male. If so, the allele is considered the putative X/Z-linked allele, and the second heterogametic allele is considered to be Y/W-linked. If both individuals are heterozygous for the same alleles, the phase cannot be inferred, and the genotype is set to missing in the heterogamete. Likewise, phasing cannot be inferred if the pair carries four unique alleles and is therefore to missing in the heterogamete.

Note that the script does not identify sex-linked regions and is applied to all sites in the provided vcf file. Since many non-sex-linked genotypes may conform to this sex-specific pattern in a single female male pair, it is up to the user to only retain genotypes within known sex-linked regions for downstream analyses.

#### 1.7 Parameter refinement

There are a few parameters that can be adjusted in the config file to improve the outcome of the pipeline. For the identification of sex-linked regions based on differences in sequence depth in step 2, the threshold at which such are classified can be adjusted (sex_depth_threshold). Changing this threshold will affect how many autosomal sites are classified as sex-linked. It will also affect how many Y/W variants identified through haplotype clustering will be retained or determined as unreliable due to low depth.

For phasing in step 4, WhatsHap can be enabled (whatshap != ”OFF”) or disabled (whatshap = ”OFF”). Read-based phasing should be useful for inferring phase in populations with high background genomic variation, abundant recombination, and in sex-linked regions with little divergence. In severely bottlenecked populations with high linkage, WhatsHap will have a relatively lesser effect on phasing. However, it is important to note that the time it takes to run WhatsHap will scale with the amount of individual heterozygosity.

For haplotype clustering in step 6 the parameters that can be adjusted include the minor allele count filter (mac), which is used filter low frequency alleles before inference of sex-linkage. This can be useful for reducing noise from species with high background genomic diversity. The window size (window) and sliding step (step) can likewise be adjusted to optimise the retrieval of sex-linked regions. It is also possible to give rare variants more weight by changing the model (dist_model) to Inverse_MAF.

For haplotype clustering in step 6, the parameters that can be adjusted include the minor allele count filter (mac), which is used to filter low frequency alleles before inference of sex-linkage. Although this does not show and impact on accuracy in our simulations, it could be worth experimenting with to reducing noise from species with high background genomic diversity. The window size (window) and sliding step (step) can likewise be adjusted to optimise the retrieval of sex-linked regions, and large window sizes seem to overall improve accuracy based on our simulations. It is also possible to give rare variants more weight by changing the model (dist_model) to Inverse_MAF. However, this does not seem to have any major effect on the outcome based on our simulations.

The user may want to run the pipeline with various setting in the haplotype clustering step. For this reason, step 6 and proceeding steps are possible to rerun with various settings and with output in separate folders, as long as the intermediate files from previous steps are retained. However, steps 1 to 5 are relatively computer intensive, and any modifications require a complete rerun.

#### 1.8 Troubleshooting know issues

It is also important to note that while sex-linkage inference based on sequence depth is relatively robust to misclassification of sex or aneuploidy in a couple of individuals, haplotype clustering is not and require precise knowledge of the frequency of sex-chromosomes in the dataset. Therefore, if known sex-linked regions are not identified as expected, it can be possible to troubleshoot such an issue by using the pipeline summary from step 10. For example, some structure due to sex-bias should be evident. Falsely sexed individuals could potentially be identified by inspecting the PCA-output of all phased variants. Likewise, the Inbreeding Coefficient can often show two separate distributions of males and females, with the heterogamete having a lower value due to inflated heterozygosity. Likewise, the Manhattan-plots containing haplotype clustering statistics should show scatter in autosomal regions, and a clear line conforming to the known number of haplotypes and individuals in the dataset. If such a line is seen, but falsely classified, it could at what criterion the algorithm failed to classify the region as sex-linked. For example, if one heterogametic individual is falsely classified as homogametic, the line conforming to sex-linkage should be classified as autosomal due to haplotypes from a homogametic individual present in the smallest cluster. If one homogametic individual is falsely classified as heterogametic, the line conforming to sex-linkage should be classified as autosomal due to heterogametic individuals are missing from the smallest cluster.

Another known issue is an undesirable bias arising when linkage pruning sex-linked data. Linkage pruning is often performed to reduce non-independence od SNPs due to linkage disequilibrium before analyses of population structure and admixture. In one dataset we had combined Z and W haplotypes among several species, and found that while these haplotypes separated well when using the full data, young strata had a tendency to cluster together following linkage pruning. We believe that the effect likely originates from a biased overrepresentation of erroneously phased Z an W SNPs. Such SNPs are effectively independent from SNPs on the gametologous haplotype, and are thus less likely to be pruned. Consequently, even a few misassigned SNPs can have an impact on analyses following linkage-pruning. We therefore evaluated the effect of alternative approaches for reducing SNP non-independence and found that thinning the dataset by picking one random SNP per X bp did not change the underlying phasing error rate. We therefore suggest thinning as an alternative approach to linkage pruning in such situations.

### 2 Datasets

#### 2.1 Simulated data

##### 2.1.1 Background

The accuracy of detection of sex-linkage and extraction of sex-linked variants may depend on a number of factors. These may be biological, such as time since recombination cessation (increases signatures of divergence), background genetic diversity (may cause noise in detecting signatures of divergence), etc. In addition, technical factors may interact with biological factors, such as various parameter settings (e.g. sliding windows size and the use of read-based phasing), or study design (i.e. the number of sequenced females and males respectively). Therefore, benchmarking of such factors is useful in informing and guiding the user on expectations based on biological factors, study design, and parameter settings on pipeline accuracy.

##### 2.1.2 Data processing

To benchmark the accuracy of PhaseWY, we performed individual-based forward simulations with SLiM 4.2.2 (Haller and Messer, 2023). We simulated a 200 kb neutrally evolving autosome, whereof half becomes XY sex-linked through complete recombination suppression in the first generation (Script based on recipe 14.5 Modeling both X and Y Chromosomes with a Pseudo-Autosomal Region (PAR) and 18.13 Tree-sequence recording and nucleotide-based models). We used a mutation rate and a recombination rate, both at 1e-8. We ran simulations at different constant effective population sizes (Ne: 100, 1,000, 10,000, 100,000, 1,000,000). To simulate different times since recombination cessation, a tree sequence of the population was outputted at different generations within the same simulation (Generation: 1,000, 10,000, 100,000, 1,000,000, 10,000,000). However, we did not run population sizes of 1,000,000 to generations longer than 1,000,000. We ran 10 replicate simulations of each effective population size. For each simulation we generated a random ancestral nucleotide sequence. We then used the tree sequence to reconstruct the ancestral recombination graph through recapitation with pyslim v.1.0.4 (https://github.com/tskit-dev/pyslim), and simulated neutral mutations with msprime v.1.3.1 (Kelleher et al., 2016) which were overlayed on the ancestral sequence. We then randomly sampled 10 females and 10 males from each tree sequence, and one additional female to use as reference sequence for that simulation and generation. These were outputted in vcf-format as a phased-truth set for later evaluation. Each individual haplotype was then converted to a chromosome length fasta-sequence using vcf2fasta from vcflib v.1.0.1 (Garrison et al., 2022). The reference allele in the vcf was then corrected based on the reference female using norm --check-ref from BCFtools v.1.14.

For each simulation and generation, we simulated paired-end 150 bp reads with a 350 bp insert size using NGSNGS v.0.9.2 (Henriksen et al., 2023). Each haplotype was simulated at 10x coverage, then concatenated to a total of 20x coverage per individual. Reads were then aligned to the first haplotype of the reference female with BWA-MEM v.0.7.17 (Li, 2013). SAMtools v.1.14 (Li et al., 2009) was used to exclude reads that were not mapped in proper pairs, not primarily aligned, or with a mapping quality score lower than 20 (view -f2 -F260 -q20). Variants were called for 10 females and 10 males with BCFtools v.1.22 multiallelic caller (Li et al., 2009), excluding alignments with a mapping quality lower than 30, alleles with a supporting base quality lower than 20, and sites with a depth lower than 3. Variants were filtered with a combination of VCFtools v.0.1.16 (Danecek et al., 2011), vcffilter from vcflib v.1.0.1, and BCFtools v.1.14.

We excluded variants with a quality score lower than 30, variants with a missingness of 0.95, variants with only heterozygous genotypes, variants with a mean depth lower than 3 and higher than 30, and genotypes with a depth lower than 3. The minimum mean depth was set low to account for lower heterogametic coverage on X when the corresponding Y region has degenerated (mean taken of female individual depth and half male depth, then divided by 3. Further, complex alleles were decomposed and normalized into their constituent SNPs and indels with a combination of vcfallelicprimitives from vcflib v.1.0.1, decompose_blocksub from vt v.0.5772 (http://github.com/atks/vt), normalize from vt v.0.5772 (Tan et al., 2015) and duplicate records removed with BCFtools v.1.14. Only sites remaining polymorphic or fixed for the alternate allele were kept. Mean individual depths for the inference of normalized sex depth differences were calculated with VCFtools on the autosomal regions of each simulated chromosome.

The simulated data was run through PhaseWY on several settings to benchmark accuracy. In total, we generated data for six statistical tests. The first two were produced to evaluate the biological factors determining when young sex-linked regions can be discovered with population genomic methods, and when they are too old for Y/W sequences to properly align to their homologous X/Z reference. Note that for neither of these tests does the parameter choice much affect the outcome other than the parameters mentioned. To test the ability to discover young sex chromosomes with sex depth differences in heterozygosity we used a single PhaseWY run with all 10 female-male pairs, MAC=1, and Window size=10 000. To test when sex chromosomes are too old for Y/W sequences to properly align to their homologous X/Z reference we used a single PhaseWY run with the sex sequence depth threshold was set to 0.75 (this setting was used in all runs). Second, we produced four datasets to evaluate how different parameter settings and study design impact successful extraction of Y genotypes. First, we used all 10 female-male pairs, the Hamming model for k-means clustering, and all 18 combinations of WhatsHap (OFF, ON), MAC (1, 5, 9), and Window size (100, 1 000, 10 000). Second, we used all 10 female-male pairs, MAC=1, WhatsHap=ON, and all six combinations of Window size (100, 1 000, 10 000), and k-means clustering model (Hamming, Inverse MAF). Third, we used WhatsHap=ON, MAC=1, the Hamming model for k-means clustering, and all 15 combinations of female-male pairs (2, 4, 6, 8, 10), and Window size (100, 1 000, 10 000). Fourth, we wanted to evaluate the successful extraction of Y genotypes in the event of very few individuals (1 female and 2 males, 2 females and 1 male). Since SHAPEIT4 does not run with less than three individuals, we ran all simulations with one male and two females (MAC=0), and all simulations with one female and two males (MAC=1). We used WhatsHap=ON, Windows size=10 000, and the Hamming model for k-means clustering. To compare the success of extracting Y/W sequences with few individuals, we also used a single female-male pair with phasing_female_male_pair.py, a companion script to PhaseWY (see section 1.6 Simple phasing with companion script). In all runs the sex sequence depth threshold was set to 0.75 and the step size ¼ that of the window size (i.e. 25, 250, 2 500).

##### 2.1.3 Analyses

We made six statistical models to evaluate how biological factors, parameter settings, and study design might influence the detection and successful extraction of Y/W sequences. In all models the predictors of population size (100, 1,000, 10,000, 100,000, 1,000,000) and generations (1,000, 10,000, 100,000, 1,000,000, 10,000,000) were included.

For the two tests evaluating how biological factors impact the  potential to discover young sex chromosomes with sex differences in heterozygosity and when sex chromosomes are too old for Y/W sequences to properly align to their homologous X/Z reference we performed. Multiple paired one-sided Wilcoxon signed-rank test (R Core Team, 2021) to assess differences were greater in the sex-linked region compared to the pseudoautosomal region. We then applied Bonferroni correction to correct for multiple testing.

For the four tests evaluating parameter settings and study design affecting the accuracy of Y/W sequence extraction we compared our simulated data to the phased-truth set. For each simulation, we extracted the Y-linked variants from the non-recombining region of the truth-set, as well as all Y-linked variants estimated from our analyses. We only used variants not considered unreliable (see Pipeline steps, Rule 8.11/9.3). Further, we did not compare variants excluded by PhaseWY due to sex depth difference. This means that our comparison only includes variants which may be extracted following alignment to a homologous reference. We calculated the total proportion of Y-linked genotypes (across all males) that were successfully extracted by PhaseWY (a measure for true positives, i.e. "sensitivity") with the help of BCFtools stats. Likewise, we calculated the proportion of inferred Y-linked genotypes that matched the truth set (a measure for false positives, i.e. "precision"). If the truth set did not contain any Y-linked variants, the proportions were set to NA to avoid division by zero. The same approach was performed on pseudoautosomal markers and X-linked markers. For these tests, co-linearity among predictors was assessed by evaluating variance inflation factors using *check_collinearity* from *performance* v.0.13.0 (Lüdecke et al., 2021). All models were built using *gamlss* v.5.5-0 (Rigby and Stasinopoulos, 2005), with a beta inflated distribution (Ospina and Ferrari, 2010). Model adequacy was assessed using randomized quantile residuals. The models were diagnosed by evaluating randomized quantile residuals. Effect sizes were calculated as $\log_{10}$ of odds ratios ($\log_{10}$OR). Furthermore, the effect of each parameter was quantified using average predictive contrasts as the average change in predicted response when moving from the minimum to the maximum simulated value, while retaining all other predictors at their observed values. Predicted responses were calculated for every observation and averaged across the dataset, yielding response-scale effect sizes representative of the entire simulated parameter space. For binary parameters, effect sizes were calculated as the average difference in predicted response between factor levels. Interactions were similarly calculated as the difference between predictor effects estimated at the minimum and maximum values of the interacting parameters.

**Sex heterozygosity difference depending on population size and generations**

We extracted the mean sex heterozygosity difference across the pseudoautosomal region, and for the sex-linked region respectively. These metrics are calculated as the mean number of heterozygote sites in heterogametes divided by the mean number in homogametes. If a simulation could not be analysed due to too little genetic variation (see rule 3.2), these metrics were both set to 1 to represent no difference.  For each combination of population size and generation we performed a paired one-sided Wilcoxon signed-rank test to assess whether sex heterozygosity differences were greater in the sex-linked region compared to the pseudoautosomal region. Each test had 10 paired observations of sex heterozygosity difference. Bonferroni correction was applied to correct for 24 tests.

**Sex depth difference depending on population size and generations**

We calculated the proportion of sites that had been excluded due to sex depth difference across the pseudoautosomal region, and for the sex-linked region respectively. These metrics are calculated as the proportion of sites were the mean normalized depth in heterogametes divided by the mean normalized in homogametes is less than 0.75 (and in Y/W linked genotypes are not considered to be extractable, see rule 2.1). For each combination of population size and generation we performed a paired one-sided Wilcoxon signed-rank test to assess whether the proportion of excluded sites were greater in the sex-linked region compared to the pseudoautosomal region. Each test had 10 paired observations of excluded sites. Bonferroni correction was applied to correct for 24 tests.

**Parameter settings for read based phasing, minor allele count filter, and window size**
We used generalized additive models for location scale and shape (GAMLSS) with a beta inflated distribution and a logit link function to model the accuracy of genome classification. We used the predictor variables: Measure (categorical variable; 6 categories: Sensitivity and Precision for Pseudoautosomal, X and Y-linked variants), population size (continuous variable), generations (continuous variable), the use of read based phasing (categorical variable; 2 categories: WhatsHap ON and WhatsHap OFF. WhatsHap OFF was set as reference), window size (continuous variable), and minor allele count filter (continuous variable). The population size, generation, and window size were $\log_{10}$ transformed. We assessed co-linearity with *performance* and found no signs of co-linearity (highest VIF = 1.52). We constructed our model by incorporating all predictors and second-degree interactions, with Measure set to interact as a third-degree term with all other predictors. The model was thus run on all data, but once with each Measure set as reference to get Measure specific coefficients. Effect sizes were calculated as $\log_{10}$ of odds ratios.

**Parameter settings for clustering model and window size**
We used generalized additive models for location scale and shape (GAMLSS) with a beta inflated distribution and a logit link function to model the accuracy of genome classification. We used the predictor variables: Measure (categorical variable; 6 categories: Sensitivity and Precision for Pseudoautosomal, X and Y-linked variants), population size (continuous variable), generations (continuous variable), window size (continuous variable), and the use of k-means clustering model (categorical variable; 2 categories: Hamming and Inverse MAF. Hamming was set as reference). The population size, generation, and window size were $\log_{10}$ transformed. We assessed co-linearity with *performance* and found no signs of co-linearity (highest VIF = 1.54). We constructed our model by incorporating all predictors and second-degree interactions, with Measure set to interact as a third-degree term with all other predictors. The model was thus run on all data, but once with each Measure set as reference to get Measure specific coefficients. Effect sizes were calculated as $\log_{10}$ of odds ratios.

**Study design for number of female-male pairs and parameter settings for window size**
We used generalized additive models for location scale and shape (GAMLSS) with a beta inflated distribution and a logit link function to model the accuracy of genome classification. We used the predictor variables: Measure (categorical variable; 6 categories: Sensitivity and Precision for Pseudoautosomal, X and Y-linked variants), population size (continuous variable), generations (continuous variable), the number of sequenced female-male pairs (continuous variable), and window size (continuous variable). The population size, generation, and window size were $\log_{10}$ transformed. We assessed co-linearity with *performance* and found no signs of co-linearity (highest VIF = 1.49). We constructed our model by incorporating all predictors and second-degree interactions, with Measure set to interact as a third-degree term with all other predictors. The model was thus run on all data, but once with each Measure set

as reference to get Measure specific coefficients. Effect sizes were calculated as $\log_{10}$ of odds ratios.

**Methods for sex-linked variant extractions using two to three individuals**
We used generalized additive models for location scale and shape (GAMLSS) with a beta inflated distribution and a logit link function to model the accuracy of genome classification. One goal of this analysis is to test how study design of few individuals limits the use of method to a simple approach (Phasing and Y extraction with a companion python script: see section 1.6 Simple phasing with companion script) and running PhaseWY with few individuals. The python script does not perform classification of sex-linkage but is intended to extract sex-linked variants in a region already known to be sex-linked. Therefore, we do not evaluate accuracy of autosomal classification with this model.

We used the predictor variables: Measure (categorical variable; 4 categories: Sensitivity and Precision for X and Y-linked variants), population size (continuous variable), generations (continuous variable), the method and number of female and male individuals (categorical variable; 3 categories: Python script 1 female and 1 male, PhaseWY 1 female and 2 males, PhaseWY 2 females and 1 male. Python script 1 female and 1 male was set as reference), and window size (continuous variable). The population size, generation, and window size were $\log_{10}$ transformed. We assessed co-linearity with *performance* and found no signs of co-linearity (highest VIF = 1.17). We constructed our model by incorporating all predictors and second-degree interactions, with Measure set to interact as a third-degree term with all other predictors. The model was thus run on all data, but once with each Measure set as reference to get Measure specific coefficients. Effect sizes were calculated as $\log_{10}$ of odds ratios.

#### 2.2 Larks (*Alauda*)

##### 2.2.1 Background

Bird have a ZW sex chromosome system, with the ancestral avian sex chromosomes formed ca 135 mya, which is shared by all present-day living birds (Neornithes; Cortez et al., 2014; Wright et al., 2012; Zhou et al., 2014). Evolutionary strata have subsequently formed independently in various bird lineages. The largest known avian sex chromosomes are described in the widespread Eurasian Skylark (*Alauda arvensis*) and the critically endangered Raso lark (*A. razae*). These sex-chromosomes are formed by the ancestral bird chromosomes, including four evolutionary strata shared by all passerines (Stratum S0, S1, S2 and S3), with at least three chromosomal translocations to the ancestral avian sex chromosomes, showing evidence of four to five additional evolutionary strata (Stratum 4A, 3a, 3b, 5 and 3c; Brooke et al., 2010; Dierickx et al., 2020; Sigeman et al., 2021, 2019). Based on evidence of sex specific patterns expected following recombination cessation, the sex-linked regions are estimated to as much as 195.3 Mbp, which constitutes 16.3 % of a 1.2 Gbp size bird genome (Sigeman et al., 2019). The nine strata vary widely in divergence time between ca 6 and 135 MYA (corresponding to ca 3 to 35 million generations; Ellerstrand and Hansson, 2026). Likewise, the extreme demographic differences between these two species has led to difference levels of background nucleotide diversity (0.01 vs 0.001 in the Skylark and Raso lark respectively; Dierickx et al., 2020). Thus, this system highlights the impact of various divergence times and background genomic diversity on successful phasing and haplotype clustering.

##### 2.2.2 Data processing

The larks were processed as part of another project including other lark species (SJ Ellerstrand et al., unpublished). Variants were called for all species together, then the relevant species were subset to per species files before PhaseWY.

The scaffolds were annotated for protein coding sequences by lift-over from the well-annotated Zebra finch assembly (Taeniopygia guttata; NCBI RefSeq accession GCF_003957565.2; Rhie et al., 2021) using Liftoff v. 1.6.3 (Shumate and Salzberg, 2021). Repetitive elements in the Skylark assembly were masked using RepeatMasker v. 4.1.5 (Smit et al., 2013).

Raw reads were quality trimmed with Trimmomatic v.0.36 (Bolger et al., 2014). Bases with a Phred score lower than 20 were trimmed from the leading and trailing ends, and in averages of 4 bases in sliding windows, finally excluding reads shorter than 100 bp. The reads were aligned to a new chromosome level genome of the Skylark (SJ Ellerstrand et al., unpublished) with BWA-MEM v.0.7.17. Duplicates were removed with MarkDuplicates from Picard v.2.23.4 (http://broadinstitute.github.io/picard). SAMtools v.1.14 was used to exclude reads that were not mapped in proper pairs, not primarily aligned, or with a mapping quality score lower than 20 (view -f2 -F260 -q20). Variants were called for the nuclear genome and mitochondrion (as haploid) separately with BCFtools v.1.22 multiallelic caller (Li et al., 2009), excluding alignments with a mapping quality lower than 20, with a depth higher than 1,000, and bases with a quality lower than 30.

Variants were then filtered with a combination of VCFtools v.0.1.16 (Danecek et al., 2011), vcffilter from vcflib v.1.0.1 (Garrison et al., 2022), and BCFtools v.1.14 (Danecek et al., 2021). Repeat regions were identified with RepeatMasker v.4.1.0 (Smit et al., 2013), using the repeat database of "Aves", and all variants within such regions were removed. We filtered variants with low quality score (< 30), high missingness (< 0.95), low depth genotypes (< 3), and maximum mean variant depth of 44 (one and a half times the mean depth of all individuals). The minimum mean depth was set low to account for lower female coverage on Z when the corresponding W region has degenerated 7 (mean taken of male individual depth and half female depth, then divided by 3). We further removed alleles which were not represented at least once on both strands, and biallelic sites which where heterozygote in all individuals. Further, haplotype variants and complex alleles were decomposed and normalized into their constituent SNPs and indels with a combination of vcfallelicprimitives from vcflib, decompose_blocksub from vt v.0.5772 (http://github.com/atks/vt), and normalize from vt (Tan et al., 2015). Only sites remaining polymorphic or fixed for the alternate allele were kept. For the mitochondrion we applied similar filters but with minimum genotype depth at 5, minimum mean depth at 5, and maximum mean depth four times the mean depth. For the mitochondrion, we only used the first 18,000 bases in the Skylark assembly due to the circular overlap of the assembly. We filtered variants with low quality score (< 30), high missingness (< 0.95), and low depth genotypes (< 3).

PhaseWY was run on the Skylark (10 females, 8 males) and the Raso lark (9 females, 9 males), using a sex sequence depth threshold of 0.75. Settings for haplotype clustering were set with mac = 1, window size = 10,000, and sliding step = 2,500. Annotated repeat regions were subtracted from the output PhaseWY bed-files of genome classification. However, note that in the final version of PhaseWY, repeat regions, etc. can be provided as a bed-file to be masked by the pipeline.

##### 2.2.3 Analyses

We performed a collection of downstream analyses based on the lark datasets to showcase examples of how the output can be applied.

**Gametolog divergence in strata of various ages**

We investigated the divergence of strata of various ages by building phylogenetic trees of respective stratum. We only used SNPs in coding regions with a minimum missingness of 0.95. We only used regions callable on both Z and W (no sex-depth difference) based on the pipeline. Genes located on the ancestral Z chromosome were further organised into evolutionary strata according to Xu et al. (2019) by cross-referencing gene names from the Zebra finch lift-over. For each stratum and individual a full sequence including invariant sites was extracted with consensus from BCFtools v1.14. Sequences from males were analysed as diploid with IUPAC ambiguity codes, and non-callable and non-coding positions were excluded. We built maximum-likelihood trees for each stratum with IQ-TREE v.2.4.0 (Minh et al., 2020), under the GTR+F+G model (GTR: general time-reversible substitution, F: use observed base frequencies, G: among-site rate heterogeneity) with 1,000 ultrafast bootstraps (Hoang et al., 2018). The branch-lengths from respective bootstrap tree were halved and used as estimate of confidence of the divergence time. The divergence times were scaled with a mutation rate of $7.16\times10^{-9}$ m/s/g (Great reed warbler; Zhang et al., 2023).

**Loss of W gametologs over time**

We previously modelled the probability of W gene-loss over time in Ellerstrand and Hansson (2026). Due to the comprehensive post-PhaseWY procedure to estimate loss of W function in that study, we here chose to only reproduce a sub-figure for illustration and refer to the methods in that paper. Briefly, W gametologs were classified as non-functional based on sex-depth differences and predicted mutational impacts using SnpEff v.4.3t (Cingolani et al., 2012). Gametologs were classified as non-functional if they had lost at least one exon, or if the fully retained transcript was fixed for one or more loss-of-function mutation (i.e. mutations causing loss of start codons, gain or loss of stop codons, or frameshifts). We then used generalized linear models (GLMs) with binomial error distribution and logit link function to test whether the probability of W non-functionality varied with stratum age (million generations, MG), species (Skylark vs. Raso lark), gene haploinsufficiency (high haploinsufficiency indicates that two copies are necessary for normal gene function; Collins et al., 2022), gene length, and their interactions. These analyses were performed on the same individuals as in this study, but using a scaffold level assembly, and an older, non-snakemake version of PhaseWY.

**Co-segregation of the W chromosome and mitochondrion**

We investigated co-segregation of the W chromosome and mitochondrion, including the Z chromosome as a control. We only used SNPs from females with a minimum missingness of 0.95. These were converted into PHYLIP format with vcfphylip.py (https://github.com/edgardomortiz/vcf2phylip). We built maximum-likelihood trees using IQ-TREE v.2.4.0 (Minh et al., 2020), including model selection (Kalyaanamoorthy et al., 2017) with ascertainment bias correction and 1,000 ultrafast bootstraps. The maximum-likelihood trees were converted to ultrametric trees using chronos from ape v.5.8.1 (Paradis and Schliep, 2019). Co-segregation patterns were visualised using cophylo from phytools v.2.4.4 (Revell, 2024).

**Deviations in nucleotide diversity between autosomes and sex chromosomes**

We investigated deviations in autosomal to Z and W nucleotide diversity ratio expected under neutral evolution, random mating, and even sex ratios (A:Z:W = 4:3:1). We only used biallelic

SNPs with a minimum missingness of 0.95. We estimated nucleotide diversity of pseudoautosomal, Z and W regions in sliding windows using the python script popgenWindows.py popgenWindows.py (https://github.com/simonhmartin/genomics_general, accessed 2025-11-10). We used the slop function from BEDTools v.2.29.2 (Quinlan and Hall, 2010) to define regions of lifted-over exon annotations and 20 kb adjoining regions. We then applied predefined windows only based on callable regions based on the pipeline, which were located ≥ 20 kb from any exon to approximate neutral evolution. We used non-overlapping windows of 10,000 b. We used the nucleotide diversity data to estimate the A:Z and A:W ratio of genetic diversity. We used 1,000 bootstraps to sample windows from each dataset and calculate ratios, which produced bootstrap distributions of ratios.

**Sexually antagonistic polymorphisms on Z**

We investigated signs of segregating sexually antagonistic polymorphisms on the Z chromosomes. We only used biallelic SNPs with a minimum missingness of 0.95. We estimated pairwise $F_{ST}$ and $D_{XY}$ between females and males in sliding windows using the python script popgenWindows.py popgenWindows.py (https://github.com/simonhmartin/genomics_general, accessed 2025-11-10). We applied predefined windows only based on callable regions annotated by the pipeline to avoid bias of missing data. We used a window size of 10,000 b with a sliding step of 2,500 b. Further, we performed a genome-wide association study (GWAS) using GEMMA v.0.98.1 (Zhou and Stephens, 2012). We estimated a centred genetic relatedness matrix (-gk 1) which was used to account for relatedness between individuals. We then run a linear mixed model (-lmm 4) and only included SNPs with a maf ≥ 0.05.

### References

Anaconda Software Distribution, 2016. Anaconda Vers. 2-2.4.0. https://anaconda.com.

Auguie, B., Antonov, A., 2017. gridExtra: Miscellaneous Functions for “Grid” Graphics.

Bolger, A.M., Lohse, M., Usadel, B., 2014. Trimmomatic: a flexible trimmer for Illumina sequence data. Bioinformatics 30, 2114–2120. https://doi.org/10.1093/bioinformatics/btu170

Brooke, M.D.L., Welbergen, J.A., Mainwaring, M.C., van der Velde, M., Harts, A.M.F., Komdeur, J., Amos, W., 2010. Widespread Translocation from Autosomes to Sex Chromosomes Preserves Genetic Variability in an Endangered Lark. J Mol Evol 70, 242–246. https://doi.org/10.1007/s00239-010-9333-3

Cingolani, P., Platts, A., Wang, L.L., Coon, M., Nguyen, T., Wang, L., Land, S.J., Lu, X., Ruden, D.M., 2012. A program for annotating and predicting the effects of single nucleotide polymorphisms, SnpEff: SNPs in the genome of Drosophila melanogaster strain w $^{1118}$ ; iso-2; iso-3. Fly 6, 80–92. https://doi.org/10.4161/fly.19695

Collins, R.L., Glessner, J.T., Porcu, E., Lepamets, M., Brandon, R., Lauricella, C., Han, L., Morley, T., Niestroj, L.-M., Ulirsch, J., Everett, S., Howrigan, D.P., Boone, P.M., Fu, J., Karczewski, K.J., Kellaris, G., Lowther, C., Lucente, D., Mohajeri, K., Nõukas, M., Nuttle, X., Samocha, K.E., Trinh, M., Ullah, F., Võsa, U., Epi25 Consortium, Estonian Biobank Research Team, Hurles, M.E., Aradhya, S., Davis, E.E., Finucane, H., Gusella, J.F., Janze, A., Katsanis, N., Matyakhina, L., Neale, B.M., Sanders, D., Warren, S., Hodge, J.C., Lal, D., Ruderfer, D.M., Meck, J., Mägi, R., Esko, T., Reymond, A., Kutalik, Z., Hakonarson, H., Sunyaev, S., Brand, H., Talkowski, M.E., 2022. A cross-disorder dosage sensitivity map of the human genome. Cell 185, 3041-3055.e25. https://doi.org/10.1016/j.cell.2022.06.036

Cortez, D., Marin, R., Toledo-Flores, D., Froidevaux, L., Liechti, A., Waters, P.D., Grützner, F., Kaessmann, H., 2014. Origins and functional evolution of Y chromosomes across mammals. Nature 508, 488–493. https://doi.org/10.1038/nature13151

Danecek, P., Auton, A., Abecasis, G., Albers, C.A., Banks, E., DePristo, M.A., Handsaker, R.E., Lunter, G., Marth, G.T., Sherry, S.T., McVean, G., Durbin, R., 1000 Genomes Project Analysis Group, 2011. The variant call format and VCFtools. Bioinformatics 27, 2156–2158. https://doi.org/10.1093/bioinformatics/btr330

Danecek, P., Bonfield, J.K., Liddle, J., Marshall, J., Ohan, V., Pollard, M.O., Whitwham, A., Keane, T., McCarthy, S.A., Davies, R.M., Li, H., 2021. Twelve years of SAMtools and BCFtools. GigaScience 10, giab008. https://doi.org/10.1093/gigascience/giab008

Delaneau, O., Zagury, J.-F., Robinson, M.R., Marchini, J.L., Dermitzakis, E.T., 2019. Accurate, scalable and integrative haplotype estimation. Nat Commun 10, 5436. https://doi.org/10.1038/s41467-019-13225-y

Dierickx, E.G., Sin, S.Y.W., van Veelen, H.P.J., Brooke, M. de L., Liu, Y., Edwards, S.V., Martin, S.H., 2020. Genetic diversity, demographic history and neo-sex chromosomes in the Critically Endangered Raso lark. Proc. R. Soc. B. 287, 20192613. https://doi.org/10.1098/rspb.2019.2613

Ellerstrand, S.J., Hansson, B., 2026. Selective regimes and evolutionary dynamics of Z and W gametologs across an expanded avian neo-sex chromosome. Genome Biology and Evolution.

Garnier, S., Ross, N., Rudis, R., Camargo, A.P., Sciaini, M., Scherer, C., 2024. viridis(Lite) - Colorblind-Friendly Color Maps for R. viridis package version 0.6.5. https://doi.org/10.5281/zenodo.4679423

Garrison, E., Kronenberg, Z.N., Dawson, E.T., Pedersen, B.S., Prins, P., 2022. A spectrum of free software tools for processing the VCF variant call format: vcflib, bio-vcf, cyvcf2, hts-nim and slivar. PLoS Comput Biol 18, e1009123. https://doi.org/10.1371/journal.pcbi.1009123

Haller, B.C., Messer, P.W., 2023. SLiM 4: Multispecies Eco-Evolutionary Modeling. The American Naturalist 201, E127–E139. https://doi.org/10.1086/723601

Hartigan, J.A., Wong, M.A., 1979. A *K*-Means Clustering Algorithm. Applied Statistics 28, 100–108. https://doi.org/10.2307/2346830

Henriksen, R.A., Zhao, L., Korneliussen, T.S., 2023. NGSNGS: next-generation simulator for next-generation sequencing data. Bioinformatics 39, btad041. https://doi.org/10.1093/bioinformatics/btad041

Hoang, D.T., Chernomor, O., von Haeseler, A., Minh, B.Q., Vinh, L.S., 2018. UFBoot2: Improving the Ultrafast Bootstrap Approximation. Molecular Biology and Evolution 35, 518–522. https://doi.org/10.1093/molbev/msx281

Kalyaanamoorthy, S., Minh, B.Q., Wong, T.K.F., von Haeseler, A., Jermiin, L.S., 2017. ModelFinder: fast model selection for accurate phylogenetic estimates. Nat Methods 14, 587–589. https://doi.org/10.1038/nmeth.4285

Kelleher, J., Etheridge, A.M., McVean, G., 2016. Efficient Coalescent Simulation and Genealogical Analysis for Large Sample Sizes. PLoS Comput Biol 12, e1004842. https://doi.org/10.1371/journal.pcbi.1004842

Knaus, B.J., Grünwald, N.J., 2017. VCFR: a package to manipulate and visualize variant call format data in R. Molecular Ecology Resources 17, 44–53. https://doi.org/10.1111/1755-0998.12549

Li, H., 2013. Aligning sequence reads, clone sequences and assembly contigs with BWA-MEM. arXiv 1303.3997.

Li, H., Handsaker, B., Wysoker, A., Fennell, T., Ruan, J., Homer, N., Marth, G., Abecasis, G., Durbin, R., 1000 Genome Project Data Processing Subgroup, 2009. The Sequence

Alignment/Map format and SAMtools. Bioinformatics 25, 2078–2079. https://doi.org/10.1093/bioinformatics/btp352
Lüdecke, D., Ben-Shachar, M., Patil, I., Waggoner, P., Makowski, D., 2021. performance: An R Package for Assessment, Comparison and Testing of Statistical Models. JOSS 6, 3139. https://doi.org/10.21105/joss.03139
Marçais, G., Delcher, A.L., Phillippy, A.M., Coston, R., Salzberg, S.L., Zimin, A., 2018. MUMmer4: A fast and versatile genome alignment system. PLoS Comput Biol 14, e1005944. https://doi.org/10.1371/journal.pcbi.1005944
Martin, M., Patterson, M., Garg, S., O Fischer, S., Pisanti, N., Klau, G.W., Schöenhuth, A., Marschall, T., 2016. WhatsHap: fast and accurate read-based phasing. bioRxiv. https://doi.org/10.1101/085050
Minh, B.Q., Schmidt, H.A., Chernomor, O., Schrempf, D., Woodhams, M.D., von Haeseler, A., Lanfear, R., 2020. IQ-TREE 2: New Models and Efficient Methods for Phylogenetic Inference in the Genomic Era. Molecular Biology and Evolution 37, 1530–1534. https://doi.org/10.1093/molbev/msaa015
Mölder, F., Jablonski, K.P., Letcher, B., Hall, M.B., Tomkins-Tinch, C.H., Sochat, V., Forster, J., Lee, S., Twardziok, S.O., Kanitz, A., Wilm, A., Holtgrewe, M., Rahmann, S., Nahnsen, S., Köster, J., 2021. Sustainable data analysis with Snakemake. F1000Res 10, 33. https://doi.org/10.12688/f1000research.29032.1
Ospina, R., Ferrari, S.L.P., 2010. Inflated beta distributions. Stat Papers 51, 111–126. https://doi.org/10.1007/s00362-008-0125-4
Paradis, E., Schliep, K., 2019. ape 5.0: an environment for modern phylogenetics and evolutionary analyses in R. Bioinformatics 35, 526–528. https://doi.org/10.1093/bioinformatics/bty633
Purcell, S., Neale, B., Todd-Brown, K., Thomas, L., Ferreira, M.A.R., Bender, D., Maller, J., Sklar, P., de Bakker, P.I.W., Daly, M.J., Sham, P.C., 2007. PLINK: A Tool Set for Whole-Genome Association and Population-Based Linkage Analyses. The American Journal of Human Genetics 81, 559–575. https://doi.org/10.1086/519795
Python Software Foundation, 2025. Python.
Quinlan, A.R., Hall, I.M., 2010. BEDTools: a flexible suite of utilities for comparing genomic features. Bioinformatics 26, 841–842. https://doi.org/10.1093/bioinformatics/btq033
R Core Team, 2021. R: A language and environment for statistical computing. R Foundation for Statistical Computing, Vienna, Austria.
Revell, L.J., 2024. phytools 2.0: an updated R ecosystem for phylogenetic comparative methods (and other things). PeerJ 12, e16505. https://doi.org/10.7717/peerj.16505
Rhie, A., McCarthy, S.A., Fedrigo, O., Damas, J., Formenti, G., Koren, S., Uliano-Silva, M., Chow, W., Fungtammasan, A., Kim, J., Lee, C., Ko, B.J., Chaisson, M., Gedman, G.L., Cantin, L.J., Thibaud-Nissen, F., Haggerty, L., Bista, I., Smith, M., Haase, B., Mountcastle, J., Winkler, S., Paez, S., Howard, J., Vernes, S.C., Lama, T.M., Grutzner, F., Warren, W.C., Balakrishnan, C.N., Burt, D., George, J.M., Biegler, M.T., Iorns, D., Digby, A., Eason, D., Robertson, B., Edwards, T., Wilkinson, M., Turner, G., Meyer, A., Kautt, A.F., Franchini, P., Detrich, H.W., Svardal, H., Wagner, M., Naylor, G.J.P., Pippel, M., Malinsky, M., Mooney, M., Simbirsky, M., Hannigan, B.T., Pesout, T., Houck, M., Misuraca, A., Kingan, S.B., Hall, R., Kronenberg, Z., Sović, I., Dunn, C., Ning, Z., Hastie, A., Lee, J., Selvaraj, S., Green, R.E., Putnam, N.H., Gut, I., Ghurye, J., Garrison, E., Sims, Y., Collins, J., Pelan, S., Torrance, J., Tracey, A., Wood, J., Dagnew, R.E., Guan, D., London, S.E., Clayton, D.F., Mello, C.V., Friedrich, S.R., Lovell, P.V., Osipova, E., Al-Ajli, F.O., Secomandi, S., Kim, H., Theofanopoulou, C., Hiller, M., Zhou, Y., Harris, R.S., Makova, K.D., Medvedev, P., Hoffman, J., Masterson, P., Clark, K., Martin, F., Howe, Kevin, Flicek, P., Walenz, B.P., Kwak, W., Clawson, H., Diekhans, M., Nassar, L., Paten, B., Kraus, R.H.S., Crawford, A.J.,

Gilbert, M.T.P., Zhang, G., Venkatesh, B., Murphy, R.W., Koepfli, K.-P., Shapiro, B., Johnson, W.E., Di Palma, F., Marques-Bonet, T., Teeling, E.C., Warnow, T., Graves, J.M., Ryder, O.A., Haussler, D., O'Brien, S.J., Korlach, J., Lewin, H.A., Howe, Kerstin, Myers, E.W., Durbin, R., Phillippy, A.M., Jarvis, E.D., 2021. Towards complete and error-free genome assemblies of all vertebrate species. Nature 592, 737–746. https://doi.org/10.1038/s41586-021-03451-0

Rigby, R.A., Stasinopoulos, D.M., 2005. Generalized Additive Models for Location, Scale and Shape. Journal of the Royal Statistical Society Series C: Applied Statistics 54, 507–554. https://doi.org/10.1111/j.1467-9876.2005.00510.x

Shumate, A., Salzberg, S.L., 2021. Liftoff: accurate mapping of gene annotations. Bioinformatics 37, 1639–1643. https://doi.org/10.1093/bioinformatics/btaa1016

Sigeman, H., Ponnikas, S., Chauhan, P., Dierickx, E., Brooke, M. de L., Hansson, B., 2019. Repeated sex chromosome evolution in vertebrates supported by expanded avian sex chromosomes. Proceedings of the Royal Society B: Biological Sciences 286, 20192051.

Sigeman, H., Strandh, M., Proux-Wéra, E., Kutschera, V.E., Ponnikas, S., Zhang, H., Lundberg, M., Soler, L., Bunikis, I., Tarka, M., Hasselquist, D., Nystedt, B., Westerdahl, H., Hansson, B., 2021. Avian Neo-Sex Chromosomes Reveal Dynamics of Recombination Suppression and W Degeneration. Molecular Biology and Evolution 38, 5275–5291. https://doi.org/10.1093/molbev/msab277

Smit, A., Hubley, R., Green, P., 2013. RepeatMasker Open-4.0. https://doi.org/http://www.repeatmasker.org

Tan, A., Abecasis, G.R., Kang, H.M., 2015. Unified representation of genetic variants. Bioinformatics 31, 2202–2204. https://doi.org/10.1093/bioinformatics/btv112

Wall, L., Christiansen, T., Orwant, J., Foy, B., Schwartz, R.L., 2020. Perl. The Perl Foundation.

Wickham, H., 2016. ggplot2: Elegant Graphics for Data Analysis. Springer-Verlag New York.

Wickham, H., Averick, M., Bryan, J., Chang, W., McGowan, L., François, R., Grolemund, G., Hayes, A., Henry, L., Hester, J., Kuhn, M., Pedersen, T., Miller, E., Bache, S., Müller, K., Ooms, J., Robinson, D., Seidel, D., Spinu, V., Takahashi, K., Vaughan, D., Wilke, C., Woo, K., Yutani, H., 2019. Welcome to the Tidyverse. JOSS 4, 1686. https://doi.org/10.21105/joss.01686

Wickham, H., François, R., Henry, L., Müller, K., Vaughan, V., Davis, 2025. dplyr: A Grammar of Data Manipulation.

Wilke, C.O., 2025. ggridges: Ridgeline Plots in "ggplot2."

Wright, A.E., Moghadam, H.K., Mank, J.E., 2012. Trade-off Between Selection for Dosage Compensation and Masculinization on the Avian Z Chromosome. Genetics 192, 1433–1445. https://doi.org/10.1534/genetics.112.145102

Xu, L., Auer, G., Peona, V., Suh, A., Deng, Y., Feng, S., Zhang, G., Blom, M.P.K., Christidis, L., Prost, S., Irestedt, M., Zhou, Q., 2019. Dynamic evolutionary history and gene content of sex chromosomes across diverse songbirds. Nat Ecol Evol 3, 834–844. https://doi.org/10.1038/s41559-019-0850-1

Zhang, H., Lundberg, M., Tarka, M., Hasselquist, D., Hansson, B., 2023. Evidence of Site-Specific and Male-Biased Germline Mutation Rate in a Wild Songbird. Genome Biology and Evolution 15, evad180. https://doi.org/10.1093/gbe/evad180

Zhao, H., Sun, Z., Wang, J., Huang, H., Kocher, J.-P., Wang, L., 2014. CrossMap: a versatile tool for coordinate conversion between genome assemblies. Bioinformatics 30, 1006–1007. https://doi.org/10.1093/bioinformatics/btt730

Zhou, Q., Zhang, J., Bachtrog, D., An, N., Huang, Q., Jarvis, E.D., Gilbert, M.T.P., Zhang, G., 2014. Complex evolutionary trajectories of sex chromosomes across bird taxa. Science 346, 1246338. https://doi.org/10.1126/science.1246338

Zhou, X., Stephens, M., 2012. Genome-wide efficient mixed-model analysis for association studies. Nat Genet 44, 821–824. https://doi.org/10.1038/ng.2310
