## Supplementary Figures S1-S15 for "PhaseWY: A pipeline for haplotype phasing, sex chromosome identification and extraction of sex-limited sequences"

Figure S12-S15: Methods for sex-linked variant extractions using two to three individuals .. 11

Figure S1: Inbreeding Coefficient ($F_{IS}$)

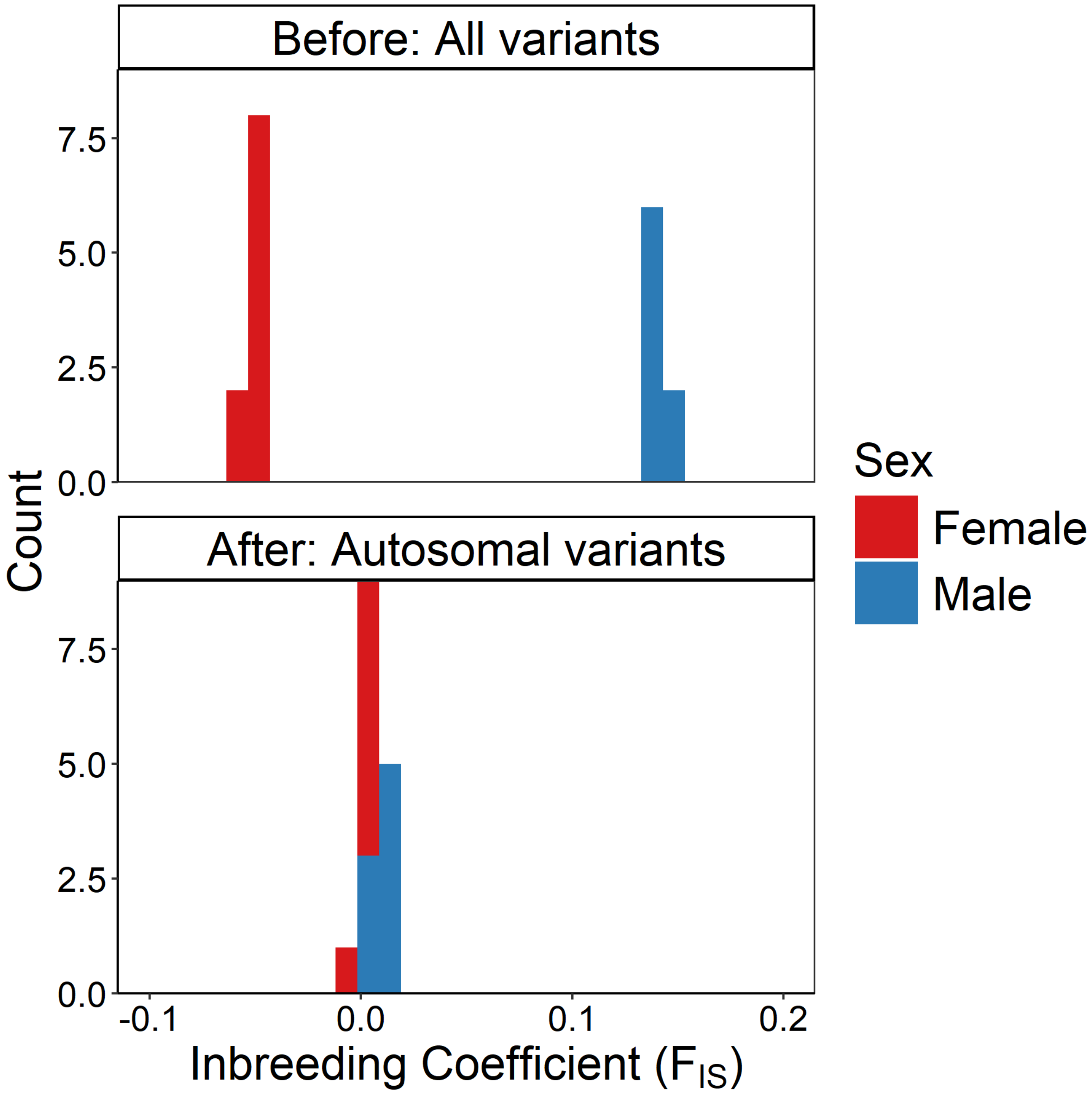

***Figure S1.*** *The inbreeding coefficient ($F_{IS}$) of male and female Skylarks based on all variants prior to PhaseWY processing, and of autosomal variants following genome classification and data subsetting with PhaseWY.*

#### Figure S2: Sex heterozygosity difference

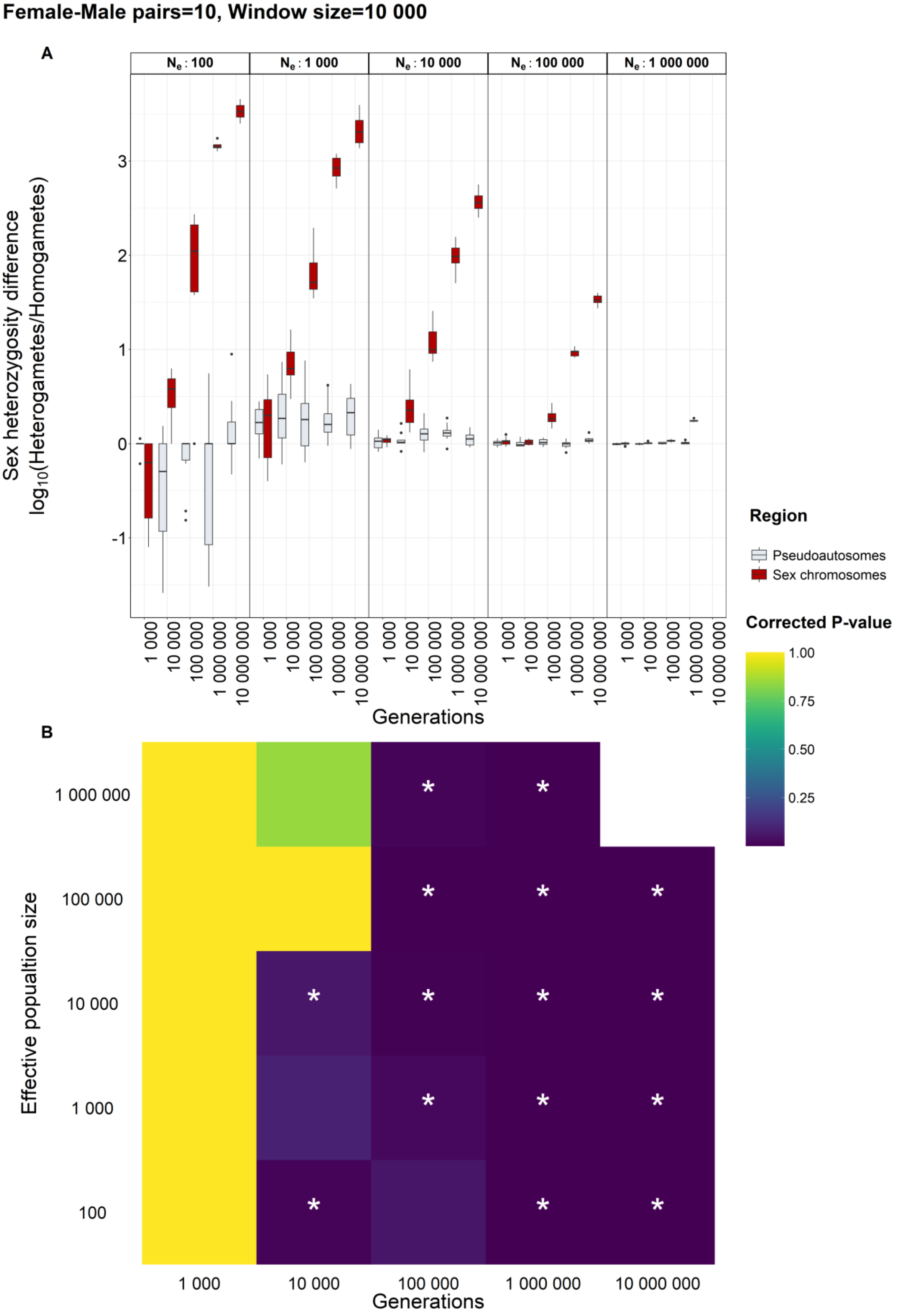

***Figure S2.*** *Simulated sex heterozygosity differences.* ***(A)*** *Raw heterozygosity differences for various effective population sizes and generations.* ***(B)*** *Bonferroni corrected P-values for respective combination of effective population size and Generations. Labels mark significance level (P < 0.1: ., P < 0.05: *, P < 0.01: **, P < 0.001: ***).*

### Figure S3: Sex depth difference

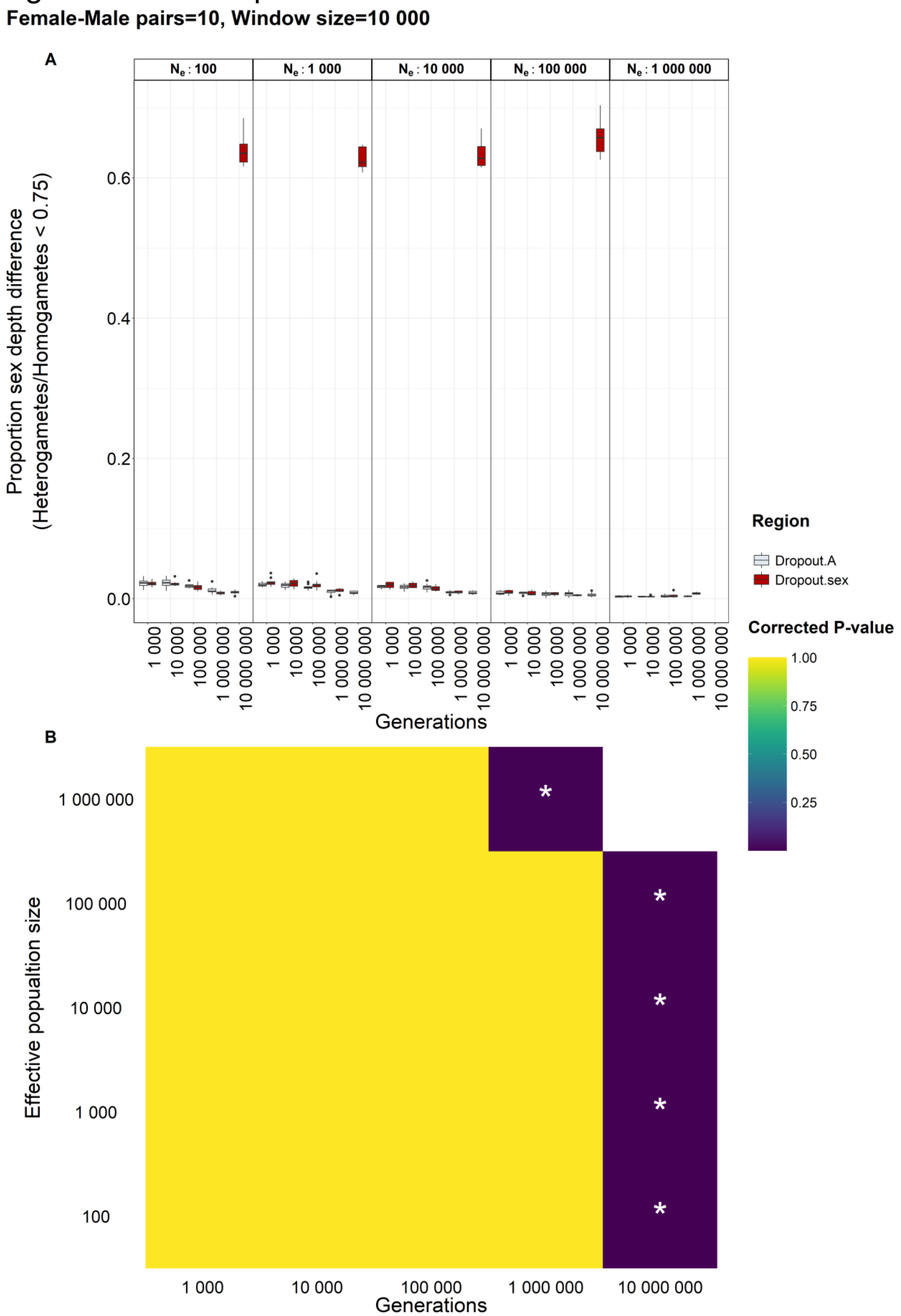

***Figure S3.*** *Simulated sex depth differences.* ***(A)*** *Raw sex depth differences for various effective population sizes and generations.* ***(B)*** *Bonferroni corrected P-values for respective combination of effective population size and Generations. Labels mark significance level (P < 0.1: ., P < 0.05: *, P < 0.01: **, P < 0.001: ***).*

### Figure S4-S6: Parameter settings for read based phasing, minor allele count filter, and window size

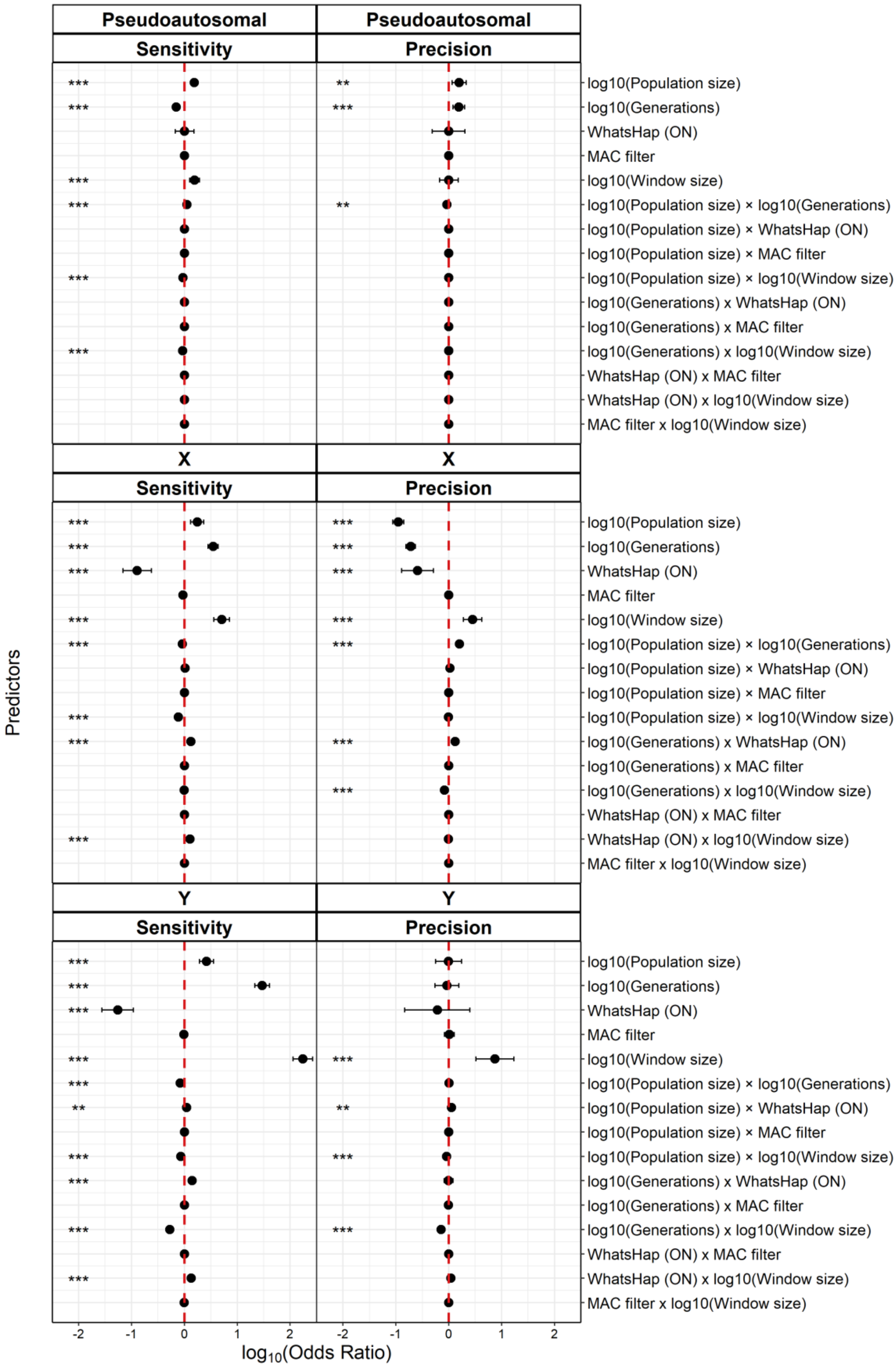

***Figure S4.*** *Effect sizes (log10 of odds ratios) for all predictors in the model on simulated data describing the test "Parameter settings for read based phasing, minor allele count filter, and window size". Estimates are derived from a generalized additive model for location scale and shape (GAMLSS) of sensitivity (true positives) and precision (false positives) in pseudoautosomal, X and Y regions.*

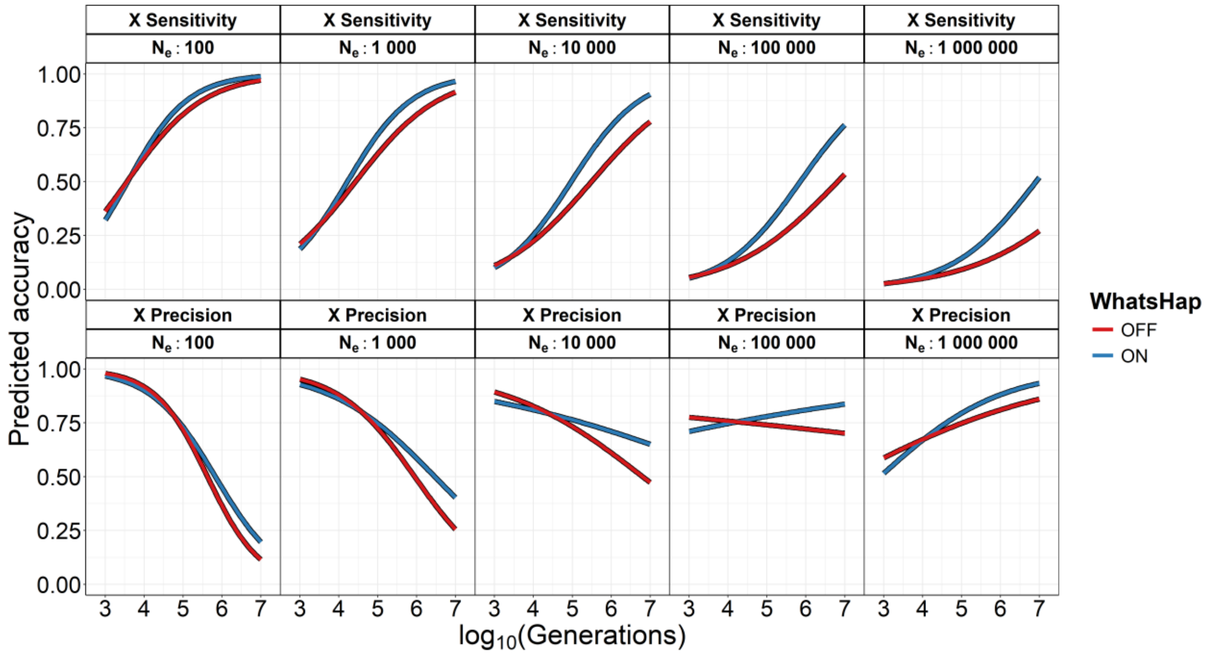

***Figure S5.*** *Sensitivity and precision in retaining X-linked variants as a function of population size, generations, and enabling or disabling WhatsHap for read-based phasing. Values are predicted with window size = 10 000, and MAC = 1.*

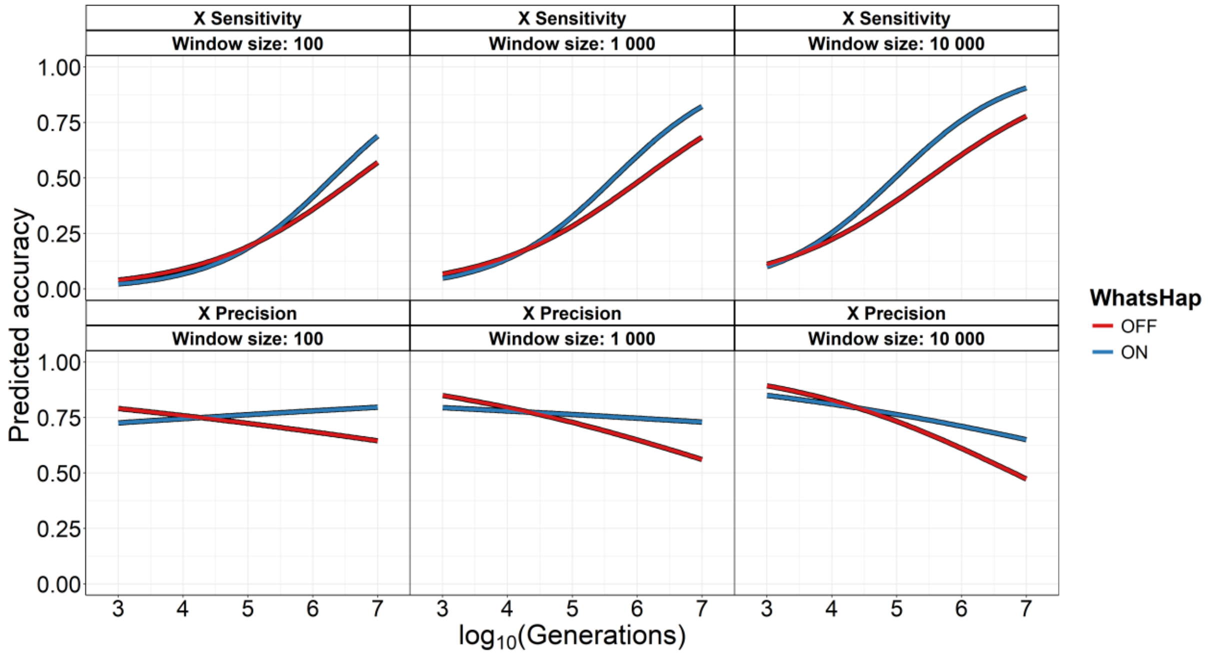

***Figure S6.*** *Sensitivity and precision in X-linked variants as a function of window size, generations, and enabling or disabling WhatsHap for read-based phasing. Values are predicted with population size = 10 000, and MAC = 1.*

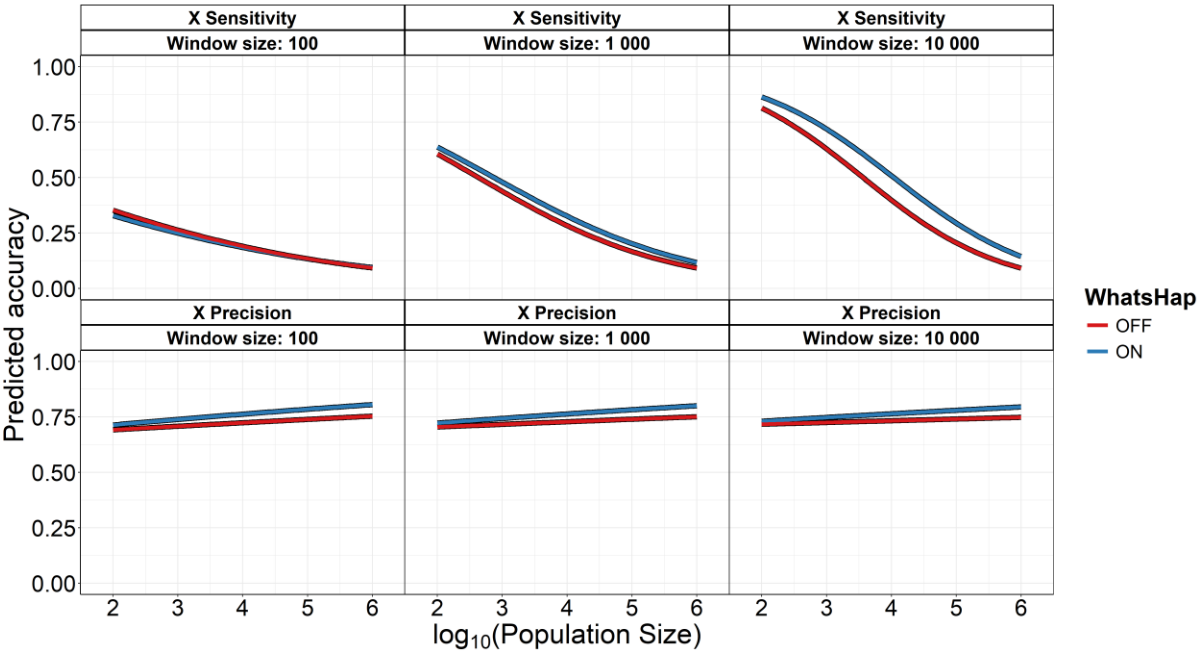

***Figure S7.*** *Sensitivity and precision in retaining X-linked variants as a function of window size, population size, and enabling or disabling WhatsHap for read-based phasing. Values are predicted with population size = 100 000, and MAC = 1.*

### Figure S8: Parameter settings for clustering model and window size

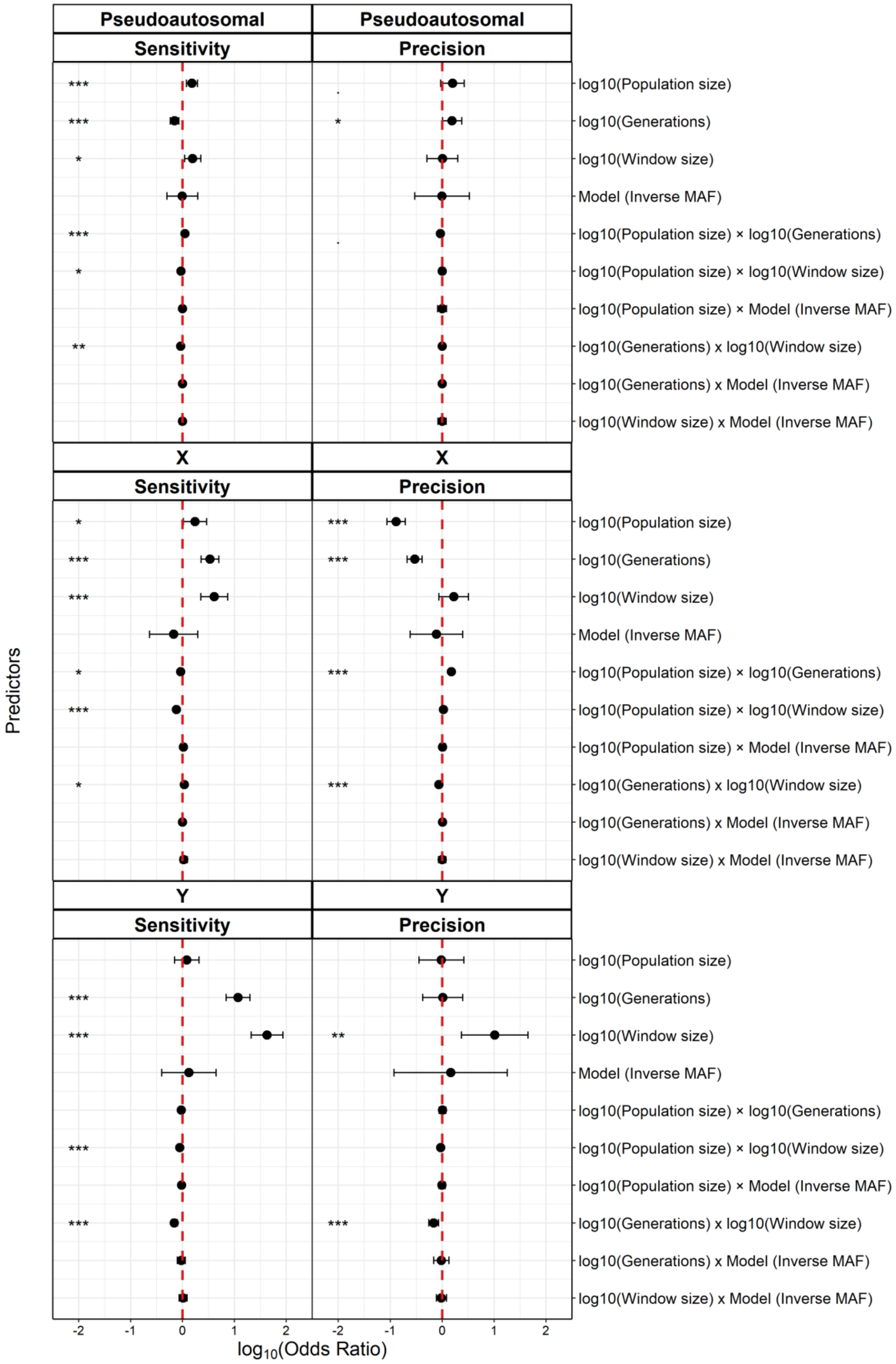

***Figure S8.*** *Effect sizes (log10 of odds ratios) for all predictors in the model on simulated data describing the test "Parameter settings for clustering model and window size". Estimates are derived from a generalized additive model for location scale and shape (GAMLSS) of sensitivity (true positives) and precision (false positives) in pseudoautosomal, X and Y regions.*

### Figure S9-S11: Study design for number of female-male pairs and parameter settings for window size

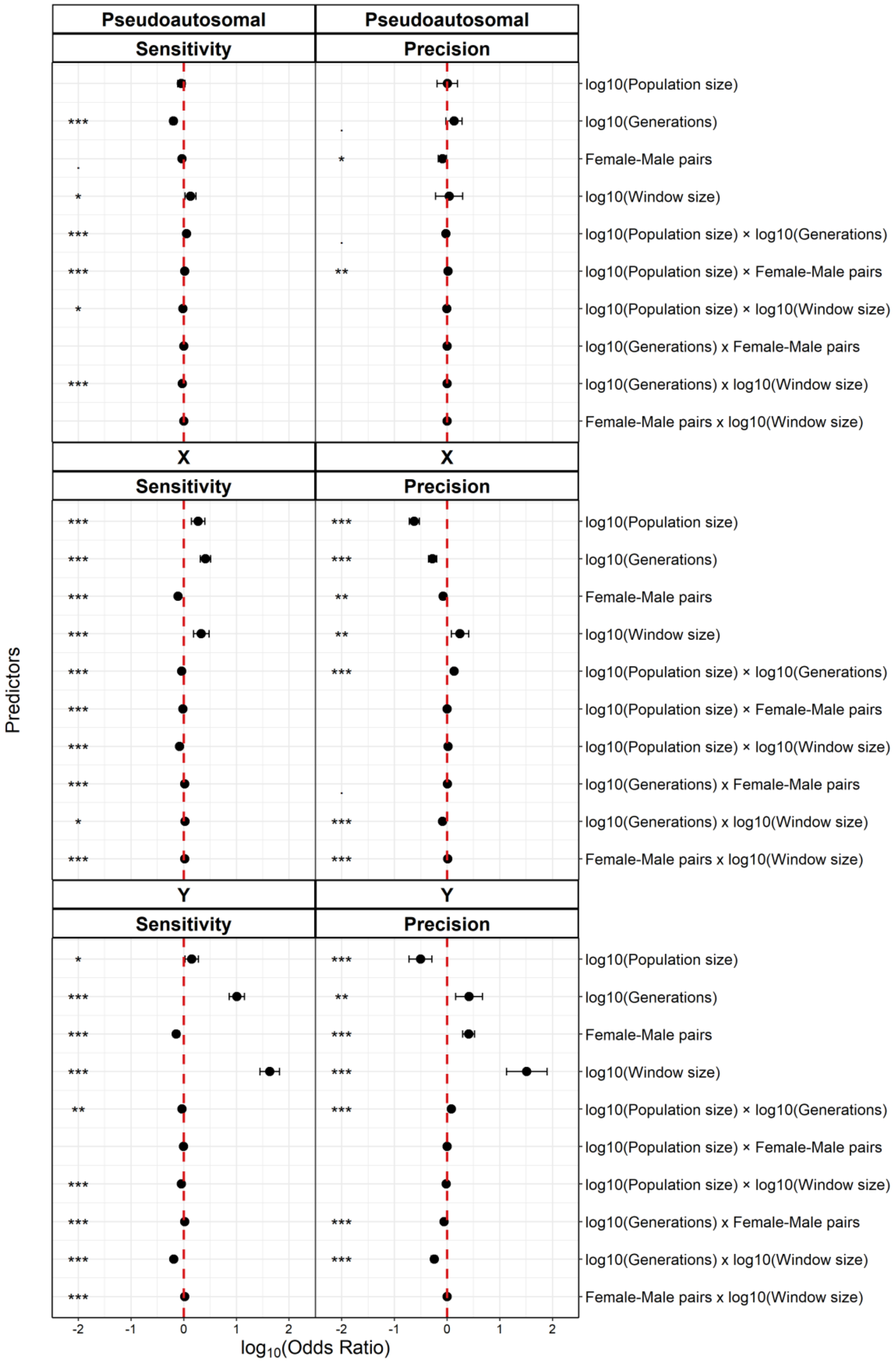

***Figure S9.*** *Effect sizes (log10 of odds ratios) for all predictors in the model on simulated data describing the test "Study design for number of female-male pairs and parameter settings for window size". Estimates are derived from a generalized additive model for location scale and shape (GAMLSS) of sensitivity (true positives) and precision (false positives) in pseudoautosomal, X and Y regions.*

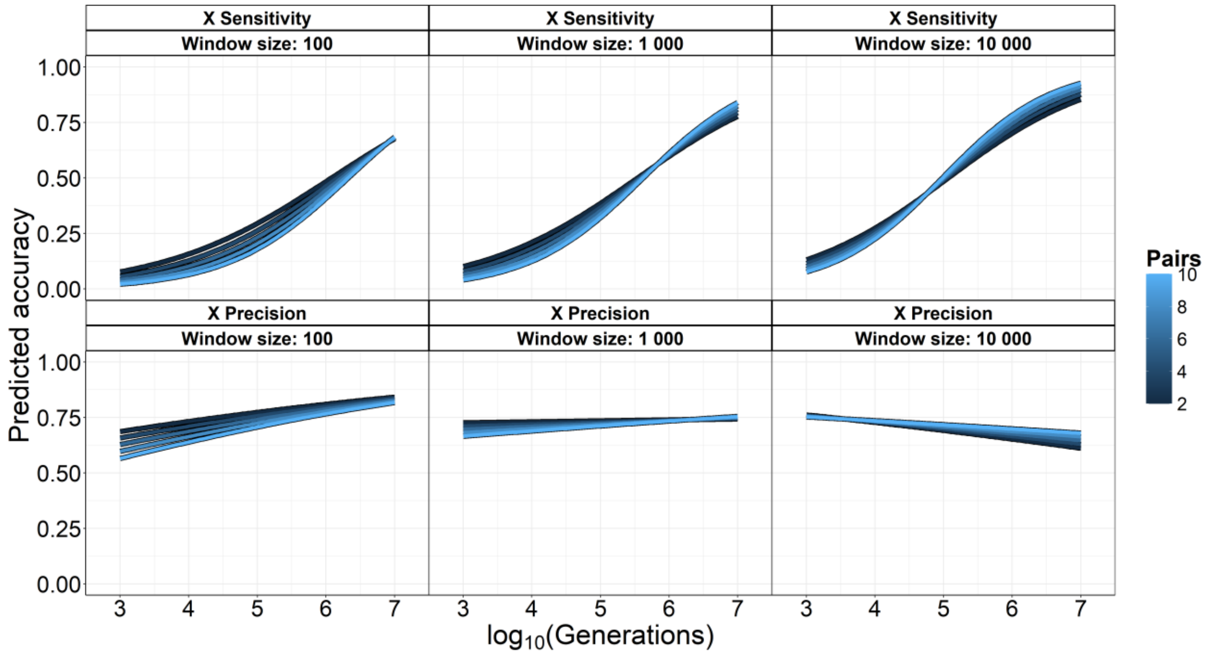

***Figure S10.*** *Sensitivity and precision in retaining X-linked variants as a function of population size, generations, and the number of female-male pairs. Values are predicted with population size = 10 000, and MAC = 1.*

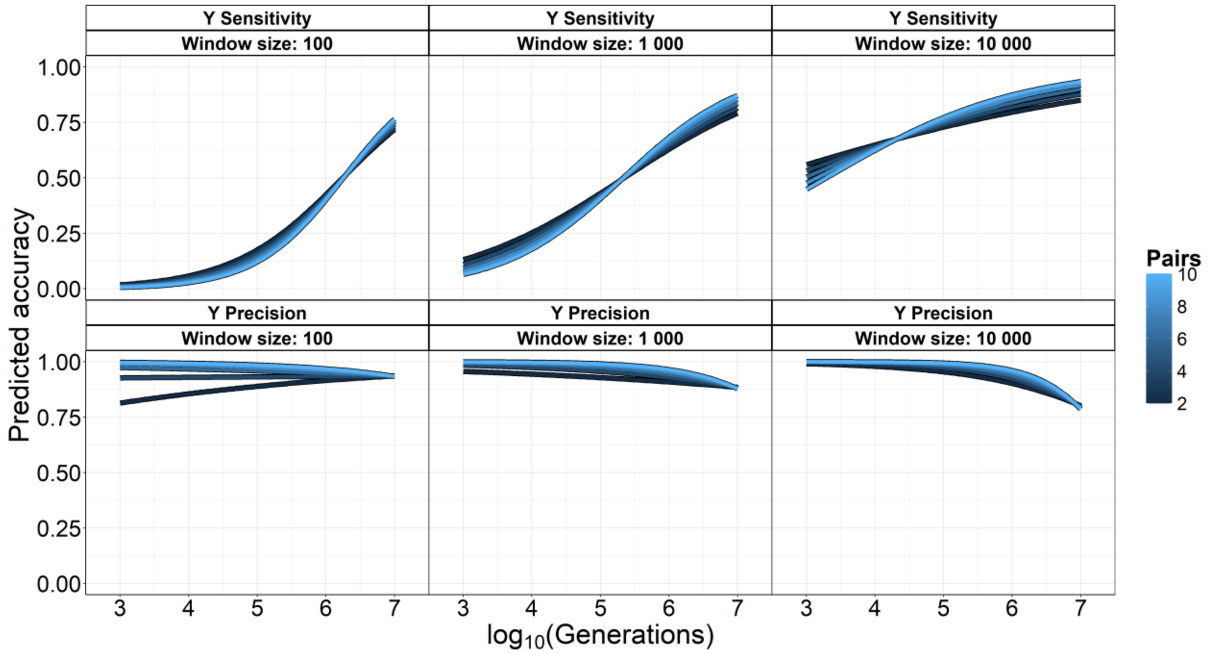

***Figure S11.*** *Sensitivity and precision in retaining Y-linked variants as a function of window size, generations, and the number of female-male pairs. Values are predicted with population size = 10 000, and MAC = 1.*

### Figure S12-S15: Methods for sex-linked variant extractions using two to three individuals

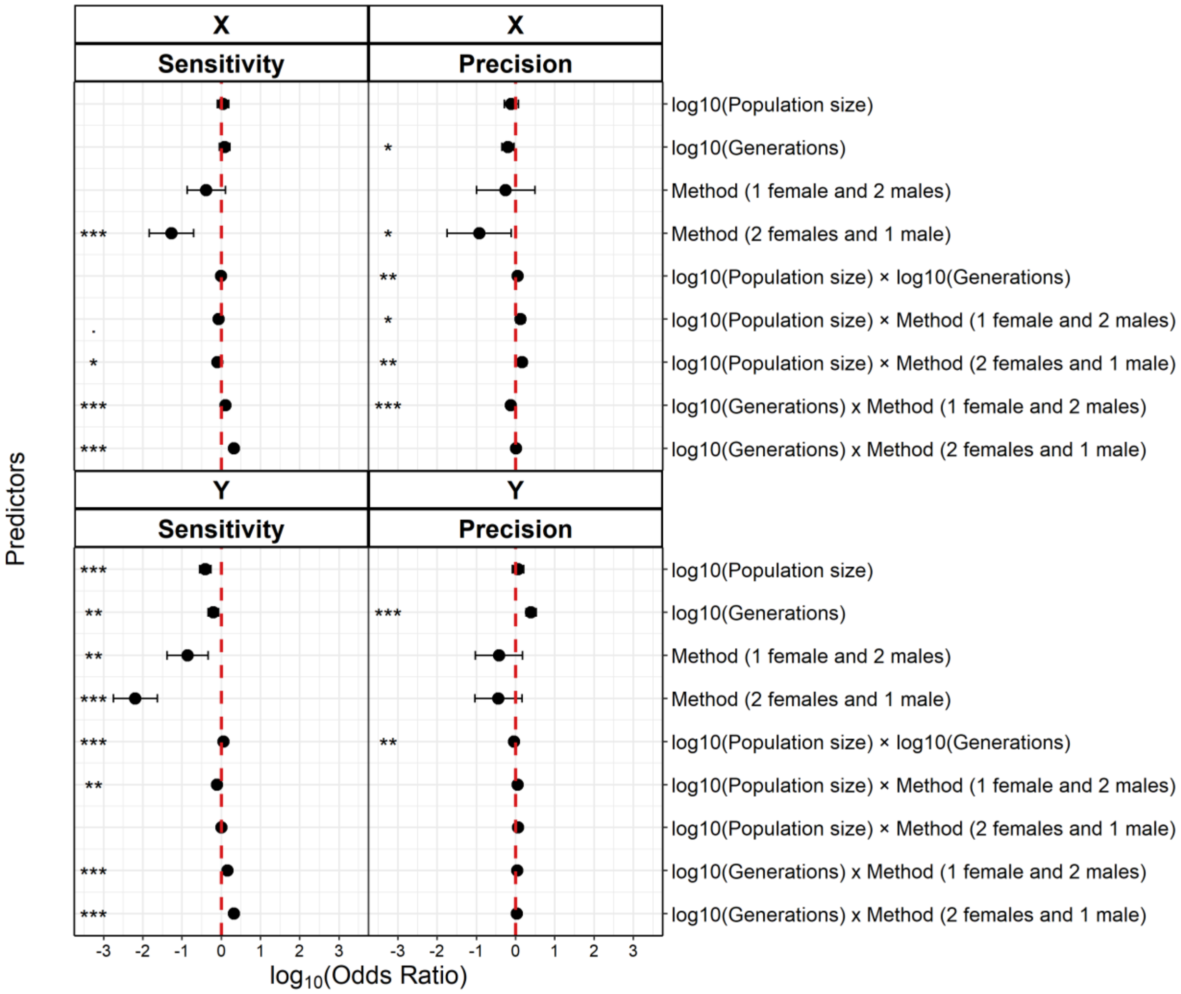

***Figure S12.*** *Effect sizes (log10 of odds ratios) for all predictors in the model on simulated data describing the test "Methods for sex-linked variant extractions using two to three individuals". Estimates are derived from a generalized additive model for location scale and shape (GAMLSS) of sensitivity (true positives) and precision (false positives) in pseudoautosomal, X and Y regions.*

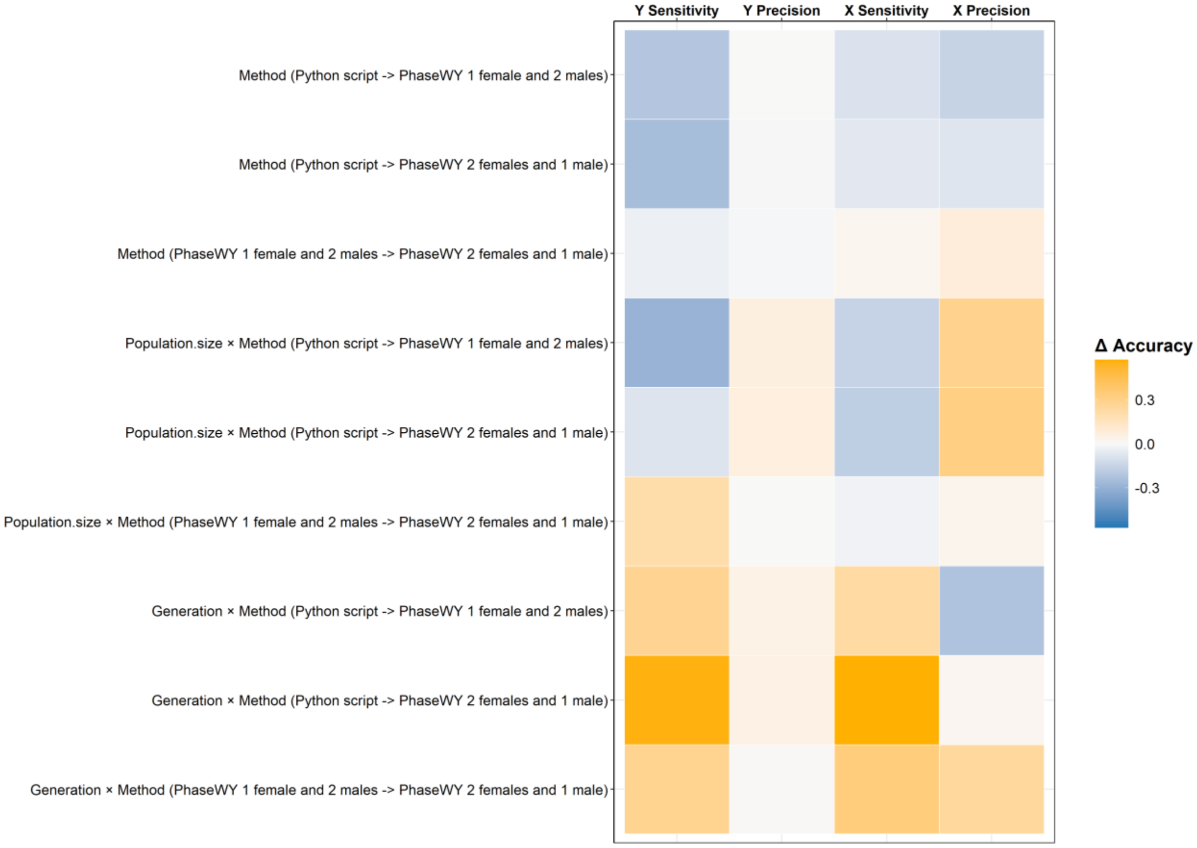

***Figure S13.*** *The average change in accuracy between the minimum and maximum value tested for respective parameter. Orange values represent an increase in accuracy, and blue values represent a decrease. Accuracy is shown for Y and X linked markers, and measured as the proportion of variants classified as true positives (Sensitivity), and the proportion of true positives out of all both true and false positives (Precision).*

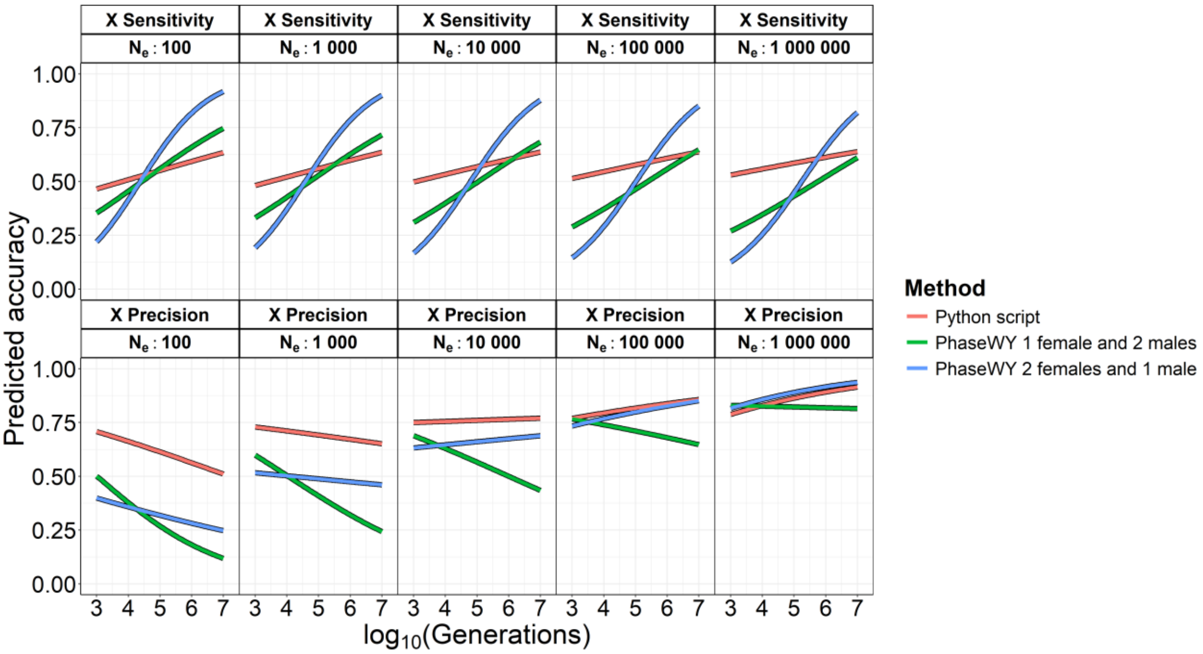

***Figure S14.*** *Sensitivity and precision in retaining X-linked variants as a function of generations, population size, and method.*

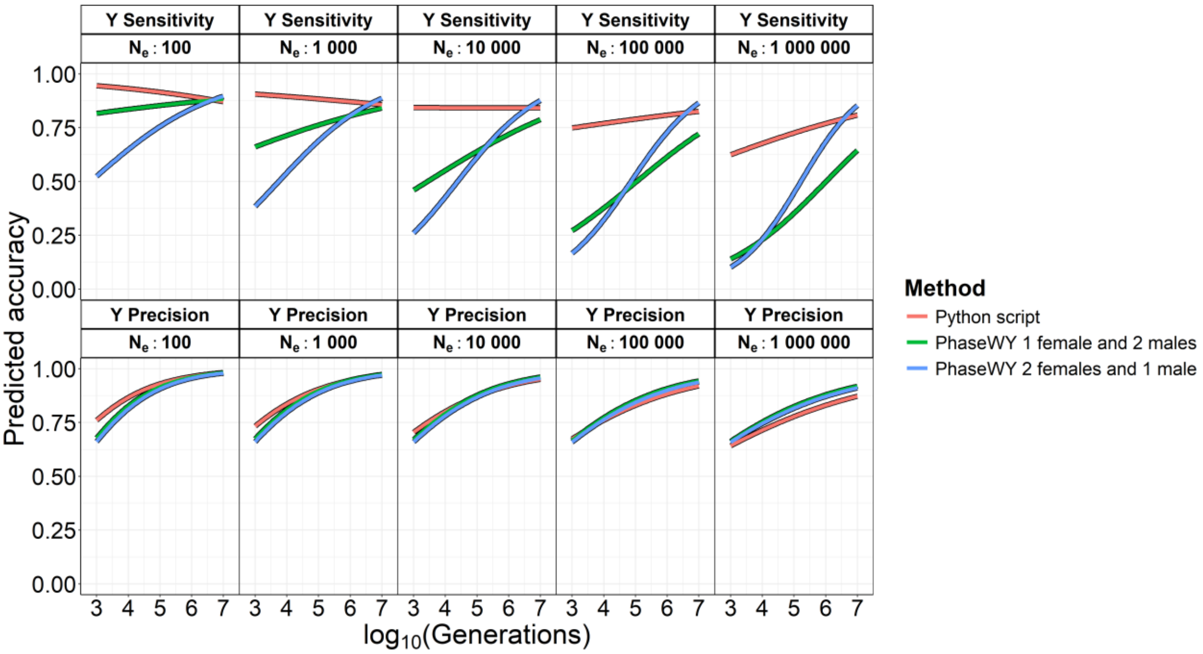

***Figure S15.*** *Sensitivity and precision in retaining Y-linked variants as a function of generations, population size, and method.*
